# DynaBiomeX: An Interpretable Dual-Strategy Deep Learning Framework for Architectural Noise Filtration in Sparse Longitudinal Microbiome Data

**DOI:** 10.64898/2026.01.28.702442

**Authors:** Awais Qureshi, Abdul Wahid, Shams Qazi, Muhammad K. Shahzad, Hashir Moheed Kiani

**Affiliations:** School of Electrical Engineering and Computer Science (SEECS), National University of Sciences and Technology (NUST), Sector H-12, Islamabad, 44500, Islamabad Capital Territory, Pakistan; School of Computer Science, University of Birmingham Dubai, Dubai, United Arab Emirates

**Keywords:** Longitudinal Microbiome Analysis, Deep Sequence Modeling, Explainable AI (XAI), Temporal Fusion Transformer, Clinical Decision Support, Zero-Inflated Data

## Abstract

**Objective:** Longitudinal microbiome datasets present unique challenges due to extreme sparsity, zero-inflation, and non-stationary behavior. Conventional Recurrent Neural Networks (RNNs) struggle to distinguish structural from sampling zeros in these contexts, limiting their utility for Clinical Decision Support (CDS).

**Methods:** We introduce DynaBiomeX, an interpretable framework specifically developed for sparse biomedical time-series. It integrates Stacking Ensembles (Bi-LSTM, GRU) with an adapted Temporal Fusion Transformer (TFT) in a unified Screener-Sentinel workflow. The Ensembles optimize collective decision boundaries to maximize sensitivity and minimize missed cases. Concurrently, the TFT functions as a Physiological Gatekeeper, utilizing Gated Residual Networks (GRN) to actively filter stochastic noise from real biological signals. We validated this approach on a multi-modal dataset of 1,871 hematopoietic cell transplantation (HCT) patients to detect gut dysbiosis.

**Results:** Stacking ensembles maximized discriminative performance (ROC-AUC = 0.912), effectively serving as high-sensitivity screeners. In contrast, the Adapted TFT functioned as a precision Sentinel, achieving zero false positives (Precision = 1.0) and high stability (MCC = 0.646). Crucially, the TFT demonstrated superior probabilistic reliability with a low Expected Calibration Error (ECE = 0.0085), addressing the “black-box” overconfidence typical of deep learning models. Ablation studies confirmed predictive robustness even without clinical covariates (ROC-AUC > 0.81).

**Conclusion:** DynaBiomeX couples sensitive screening with precise, calibrated validation to robustly analyze sparse longitudinal data. Validated on microbiome dysbiosis, this framework offers a scalable template for zero-inflated domains like single-cell sequencing and EHR monitoring.

**Highlights:**

- DynaBiomeX integrates Stacking Ensembles and Temporal Fusion Transformers for microbiome risk stratification.
- Gated Residual Networks distinguish stochastic sampling zeros from structural zeros in sparse ASV data.
- A novel Screener-Sentinel workflow resolves the trade-off between surveillance sensitivity and diagnostic precision.
- Clinical metadata functions as a physiological gatekeeper, filtering noise to achieve zero false positives (MCC 0.646).
- The framework minimizes alarm fatigue in Auto-FMT decision support by validating latent dysbiosis signals.

## 1. Introduction

Dysbiosis—an alteration in the normal composition and function of the intestinal microbiome—has emerged as a critical factor influencing adverse outcomes in high-risk populations, particularly among patients undergoing hematopoietic cell transplantation (HCT). Following HCT, the microbiome undergoes profound destabilization driven by chemotherapy, radiation, and antibiotic prophylaxis. This disruption is strongly associated with increased morbidity, including heightened risks of graft-versus-host disease (GVHD), bloodstream infections, and disease relapse. Timely and accurate detection of dysbiosis is therefore essential for early clinical intervention.

However, a significant gap persists between biological standards and computational methodologies. Most studies in the field of predictive microbiome modeling have only focused on legacy Operational Taxonomic Unit (OTU) clusters to bypass high-dimensionality. Such approaches, however, have failed to address the loss of resolution inherent in clustering. While Amplicon Sequence Variants (ASVs) have now become the biological benchmark, they introduce a major challenge: extreme sparsity and zero-inflation. Previous published studies are limited to dense datasets [1, 2, 3], failing to test how models behave when profiles are dominated by zeros.

Conventional analytical approaches often fail to distinguish between technical zeros (sampling artifacts) and biological zeros (true absence). As a result, models relying on reactive symptoms or single-time-point analyses cannot capture the complex, non-linear dynamics of this ecosystem, underscoring the need for advanced deep learning architectures capable of filtering high-dimensional noise.

### Statement of Significance

## 2. Related Work

The transition from static to dynamic analysis represents a pivotal shift in precision medicine. While classical univariate and multivariate statistical methods [4] provided the foundation for microbiome association studies, they fail to capture the complex, non-linear temporal interactions inherent in longitudinal meta-omics data. Consequently, the field has pivoted toward Deep Learning (DL) methodologies [5], answering the call for advanced dynamic methods capable of handling personalized longitudinal profiles [6].

### 2.1. Recurrent Sequence Modeling

Recurrent architectures emerged as a cornerstone for modeling longitudinal data, as early studies demonstrated their effectiveness [7, 8]. For instance, [9] leveraged LSTMs paired with sparse autoencoders to analyze genus-level microbiome profiles, while [3] harnessed GRUs to infer host status based on OTU clusters. These foundational models soon found broader application, such as predicting food allergies [9] and forecasting bacterial growth [10]. To navigate the challenge of high-dimensional data, researchers developed hybrid architectures: Phylo-LSTM, introduced by [1], applied phylogenetic sorting to capture spatial features, and [11] explored self-knowledge distillation. Despite these innovations, most methods continued to aggregate data at the OTU or genus level, limiting resolution compared to Amplicon Sequence Variants (ASVs). More recent efforts, such as Neural Controlled Differential Equations (Neural CDE) [12] and ensemble RNNs [13], sought to address the complexity of irregular sampling; however, they often faced persistent challenges with overfitting in sparse, zero-inflated datasets.

### 2.2. Imputation and Generative Approaches

As researchers grappled with the challenge of sparse data, a new avenue emerged: data augmentation. Reviews reveal that RNNs became a go-to solution for imputing missing values [14], while deep learning models stepped in to interpolate absent time points [13]. The field saw the rise of sophisticated systems like Deep-MicroGen—a GAN-based framework [2]—and conditional score-based diffusion models (CSDI) [15]. Although these generative approaches excel at smoothing trajectories, they sometimes fabricate microbial abundances or dampen sharp signals, raising concerns about introducing synthetic artifacts into sensitive clinical datasets.

### 2.3. Transformers and Interpretability

The application of Transformer architectures to microbiome data is an emerging area for global context modeling [16]. For instance, PhenoDP [17] and MOMTM [18] have demonstrated the utility of attention mechanisms. Nevertheless, these short-term studies do not sufficiently address the behavior of attention weights under the pronounced zero-inflation characteristic of amplicon sequence variant (ASV)-level data. Furthermore, the study by [19] implemented Transformers with Robust Principal Component Analysis (TRPCA) on ASV-level data, yet reported only modest predictive improvements. Concurrent research has emphasized the identification of time-varying biomarkers through interpretable machine learning frameworks [20].

### 2.4. Ensemble Integration

Ensemble methods have become increasingly prominent due to their ability to enhance model stability and accuracy. [21] introduced LP-Micro, while [22] demonstrated the effectiveness of CNN-ResNet-LSTM ensemble architectures. Similarly, [23] reported that Random Forest approaches focusing on functional potential can surpass taxonomic-based methods.

#### Motivation and Contribution

Despite considerable progress, the majority of research to date has remained primarily descriptive, leaving several pivotal challenges unresolved. First, a significant gap persists between biological standards and computational methodologies. Although Amplicon Sequence Variants (ASVs) now represent the state-of-the-art in high-resolution analysis, most deep learning models for time-series imputation continue to be bench-marked on legacy Operational Taxonomic Unit (OTU) data [1, 24]. These models frequently encounter difficulties arising from the pronounced sparsity characteristic of ASV profiles, and often fail to reliably distinguish between structural zeros (reflecting true biological absence) and sampling zeros (stemming from technical artifacts) without resorting to aggressive imputation strategies that alter the underlying data structure.

|  |  |
| --- | --- |
| **Problem or Issue** | Longitudinal microbiome datasets suffer from extreme sparsity and zero-inflation. Distinguishing **structural zeros** (true absence) from **sampling zeros** (artifacts) remains a computational bottleneck, causing high false-positive rates and clinical alarm fatigue. |
| **What is Already Known** | Recurrent Neural Networks (RNNs) capture temporal dynamics but fail in non-stationary, sparse settings. Existing solutions often rely on coarse taxonomic aggregation or black-box methods that sacrifice precision for sensitivity, obscuring biological drivers. |
| **What This Paper Adds** | This study introduces DynaBiomeX, a “Screener-Sentinel” dual-strategy. It unifies Stacking Ensembles for high-sensitivity screening with an Adapted Temporal Fusion Transformer (TFT) for high-precision validation. The TFT acts as a physiological gatekeeper, leveraging clinical metadata to filter stochastic noise and achieve zero false positives. |
| **Who would benefit from the knowledge in this paper** | Clinical informatics researchers, clinicians managing HCT dysbiosis, and data scientists developing models for zero-inflated domains (e.g., single-cell sequencing, EHR monitoring). |

Second, the issue of interpretability remains inadequately addressed. Although individual Explainable AI (XAI) techniques are widely employed, comprehensive frameworks that systematically integrate model-agnostic approaches with architecture-specific attention mechanisms have received limited attention [11]. As a result, many current models continue to operate as black boxes, demonstrating diminished performance on external datasets [12, 21] and falling short of delivering the transparent risk stratification essential for clinical decision support.

This study addresses these gaps by proposing DynaBiomeX, a supervised methodological framework tailored for sparse, high-resolution ASV data. Building upon our previous unsupervised framework [25], we introduce a Dual-Strategy Clinical Decision Support (CDS) Workflow:

- The Screener: We utilize ensemble strategies and bi-directional recurrent models to maximize sensitivity for early risk stratification.
- The Sentinel: We introduce an Adapted Temporal Fusion Transformer (TFT) utilizing Gated Residual Networks (GRN). The GRN functions as a non-linear sparsity filter, distinguishing predictive biological signals from stochastic sampling noise.

By integrating these perspectives with rigorous ablation studies, DynaBiomeX contributes a robust, interpretable blueprint for microbiome-based risk stratification in HCT patients.

## 3. Methods

This study implemented deep sequence models for detecting dysbiosis episodes in longitudinal microbiome and clinical data. The methodology goes forward as data preprocessing, feature engineering, model development, and comprehensive evaluation protocols. The overall architecture of the proposed DynaBiomeX framework is illustrated in Figure 1.

**Figure 1.**
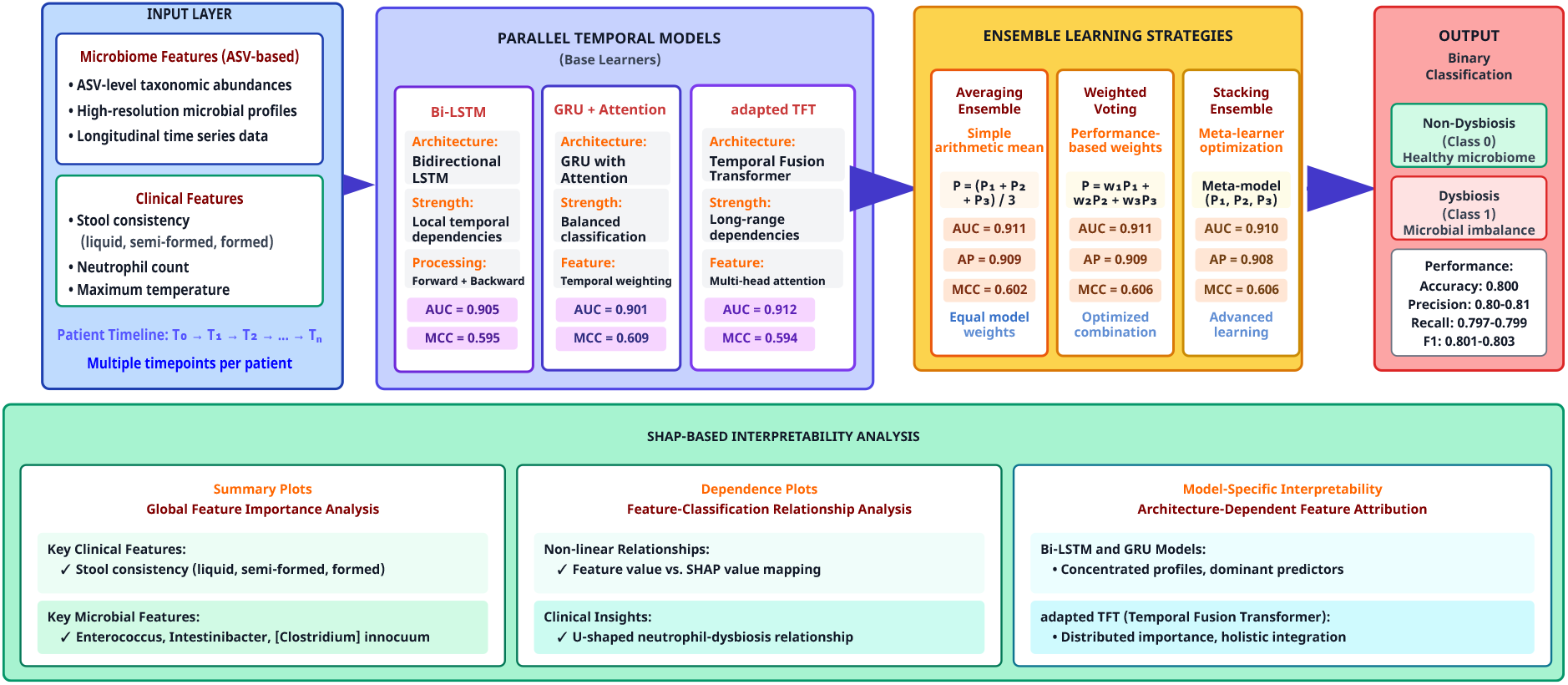
DynaBiomeX framework for dysbiosis classification in HCT patients. (Top Left) Input layer integrates ASV-based microbiome features and clinical markers from longitudinal patient data. (Top Center) Three parallel temporal models process sequential data: Bi-LSTM (bidirectional processing, AUC=0.905), GRU+Attention (temporal weighting, AUC=0.901, MCC=0.609), and TFT (multi-head attention for long-range dependencies, AUC=0.912). (Top Center-Right) Ensemble learning strategies combine base model classifications through Averaging (AUC=0.911), Weighted Voting (AUC=0.911, MCC=0.602), and Stacking with meta-learner optimization (AUC=0.910, MCC=0.606). (Top Right) Output layer produces binary classification (Non-Dysbiosis vs. Dysbiosis) with overall accuracy of 0.800 and F1-scores of 0.801-0.803. (Bottom) SHAP-based interpretability module provides: (i) Summary Plots identifying key clinical features (stool consistency, neutrophil count) and microbial taxa (*Enterococcus, Intestinibacter, [Clostridium] innocuum*), (ii) Dependence Plots revealing non-linear relationships and U-shaped neutrophil-dysbiosis patterns, and (iii) Model-Specific Interpretability showing that Bi-LSTM and GRU exhibit concentrated attribution profiles while adapted TFT demonstrates distributed microbial importance with holistic integration. All ensemble strategies enhance classification stability and clinical reliability compared to individual models.

### 3.1. Dataset Selection Rationale

This study utilized the longitudinal hematopoietic cell transplantation (HCT) microbiome dataset described in [25], comprising 514,246 time-point observations from 1,871 patients (up to 51 sequential observations per patient). Each observation integrates high-resolution amplicon sequence variant (ASV) profiles (412 bacterial genera) with clinically relevant metadata (neutrophil counts, maximum body temperature, and stool consistency). This multimodal design preserves the temporal dependencies critical for capturing interactions between gut microbiota and host physiological states. Preliminary analysis confirmed challenges typical of this domain, including non-stationary temporal patterns, zero-inflation in taxonomic abundances, and compositional constraints, presenting a rigorous benchmark for longitudinal microbiome modeling. Detailed collection protocols are provided in [26].

### 3.2. Model Selection Rationale: From Anomaly Detection to Risk Stratification

Building on our previous unsupervised anomaly detection work [25], this study advances to precise risk stratification. While prior reconstruction-based architectures captured latent health shifts, they fundamentally lacked the capacity to differentiate between stochastic sampling noise and informative structural sparsity. To overcome this, we prioritized architectures capable of explicitly gating noise through Gated Residual Networks (Supplementary S2-S4), managing the trade-off between screening sensitivity and diagnostic precision:

#### 3.2.1. Bidirectional LSTM (Bi-LSTM)

The Bi-LSTM captures temporal dependencies in both forward and backward directions, utilizing future context to explicitly model sequential relationships. This bidirectional context improves classification accuracy for dysbiosis events compared to unidirectional baselines.

#### 3.2.2. GRU with Attention Mechanism

This architecture combines the computational efficiency of Gated Recurrent Units with attention-based feature selection. The attention mechanism identifies informative bacterial genera and time points, directly addressing high-dimensional sparsity while providing interpretable insights into taxonomic drivers.

#### 3.2.3. Adapted Temporal Fusion Transformer (TFT) Architecture

We adapted the TFT architecture, originally designed for forecasting [27], to perform binary classification of zero-inflated ASV inputs. This adaptation preserves interpretability through specific components:

##### Variable Selection and Encoding

To manage high dimensionality (*D* > 412), Variable Selection Networks (VSN) limit the search space. For input vector **x**_*t*_, variable selection is defined as:

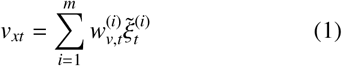

where 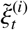 is the transformed input feature and 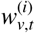 are softmax-generated weights. This dynamically suppresses irrelevant taxa (noise) while amplifying biomarker signals.

##### Gated Residual Networks (GRN) as Noise Filters

We utilized GRNs to capture non-linear microbial-clinical relationships:

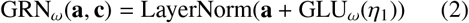

GRN_ω_(**a, c**) = LayerNorm(**a** + GLU_ω_(η_1_)) (2) where GLU is the Gated Linear Unit activation. *Theoretical Basis of Noise Filtration:* The GRN functions as a dynamic relevance filter for zero-inflation. The gating coefficient *γ* ∈ [0, 1] differentiates zero types:

- **Structural Zeros (Signal):** For taxa where absence correlates with the target (e.g., commensal loss), *γ* → 1, allowing signal propagation.
- **Sampling Zeros (Noise):** For stochastic zeros lacking clinical correlation, *γ* → 0.

This mechanism suppresses sampling noise before temporal processing, enabling robust learning from sparse data without generative imputation.

##### Classification Head Adaptation

We modified the decoding head to output a binary probability *ŷ* rather than quantile forecasts. Temporal features are aggregated into context vector *ψ* and passed through a sigmoid activation:

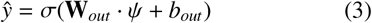

#### 3.2.4. Ensemble Learning Strategies

To mitigate variance, we combined predictions from Bi-LSTM (*P*_*lstm*_), GRU (*P*_*gru*_), and TFT (*P*_*t f t*_) using three strategies:

a. **Averaging Ensemble:** Assumes equal contribution, computing the arithmetic mean:

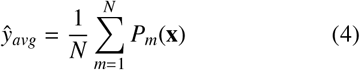

where *N* = 3.
b. **Weighted Voting:** Prioritizes discriminative power by assigning weights *w*_*m*_ proportional to validation ROC-AUC:

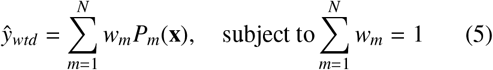

where 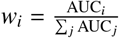
c. **Stacking Ensemble:** Utilizes a Logistic Regression meta-learner trained on validation predictions to optimize non-linear combinations:

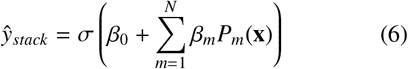

where *β*_*m*_ represents the learned coefficient for model *m*.

### 3.3. Data Preprocessing and Feature Engineering

Given the high sparsity (> 90%) of ASV profiles, we employed Centered Log-Ratio (CLR) transformation with multiplicative replacement - pseudo-count addition to mitigate compositionality effects without resorting to generative imputation. Detailed preprocessing protocols, including the numerical imputation of sub-threshold neutrophil counts and one-hot encoding of stool consistency, are provided in the Supplementary Materials (S1). These steps preserve biological signal integrity while preparing sparse inputs for the Gated Residual Network filtration mechanism.

#### 3.3.1. Target Variable Definition and Temporal Modeling

We implemented a concurrent dysbiosis detection framework utilizing a 14-day sliding window (*t* to *t*+13) to identify status at the final time-point. Dysbiosis was strictly defined as a sustained (≥2 days) concurrence of fever (> 38.0^◦^C), neutropenia (< 500 cells/µL), and liquid stool consistency. This definition prioritizes diagnostic accuracy for real-time decision support over prospective forecasting.

Unlike prognostic models that forcast events with advance warning, our framework provides high-precision confirmation of dysbiosis presence at the time of assessment, supporting immediate treatment initiation, therapeutic adjustments, and patient management decisions in continuously monitored HCT settings.

### 3.4. Data Structuring and Training Protocol

Data were organized into fixed-length sequences of 14 consecutive time points per patient. To prevent leakage, we employed patient-level stratified random sampling (70% training, 15% validation, 15% testing), with feature selection and standardization fitted exclusively on the training split. All architectures were trained using binary cross-entropy loss and the Adam optimizer, utilizing Early Stopping (patience=10) and checkpointing based on validation AUC. Experiments were conducted on a Google Colab Pro TPU v2-8 environment, detailed specifications are provided in Supplementary Material S1, S7 and S8.

#### 3.4.1. Performance Evaluation Metrics

We assessed performance using a stratified metric set tailored to the Screener-Sentinel workflow:

- **Screening Metrics (Sensitivity Focus):** ROC-AUC and Recall were prioritized to evaluate the Ensemble’s capacity to minimize false negatives.
- **Sentinel Metrics (Reliability Focus):** Precision and Matthews Correlation Coefficient (MCC) were used to quantify robustness against false positives in zero-inflated data.
- **Calibration Metrics (Trustworthiness Focus):** To assess the reliability of confidence scores for clinical deployment, we utilized the Brier Score and Expected Calibration Error (ECE). The Brier Score measures the mean squared difference between predicted probabilities and actual outcomes, while ECE quantifies the divergence between expected accuracy and empirical accuracy across *M* = 10 bins.

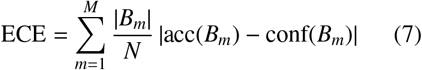

where *N* is the total number of samples, acc(*B*_*m*_) is the observed accuracy in bin *m*, and conf(*B*_*m*_) is the mean predicted confidence.

- **Global Metrics:** Standard measures including Accuracy, Loss, and PR-AUC were tracked to ensure convergence.

##### Classification Threshold Strategy

To ensure an unbiased comparison across all evaluated architectures, a fixed classification threshold of τ = 0.5 was applied uniformly. An instance was classified as *Dysbiosis* if *P*(*y* = 1 | *x*) ≥ 0.5. This threshold was not optimized on the validation set to avoid overfitting. The validity of this fixed threshold is supported by our calibration analysis (see Results), which demonstrates that the proposed TFT model achieves a low ECE (0.0085), indicating that the default decision boundary accurately reflects the true likelihood of the condition.

### 3.5. Statistical Analysis

Statistical significance was assessed via a boot-strapped paired t-test protocol (*n* = 1, 000 iterations) with Holm-Bonferroni correction (α = 0.05) to control the family-wise error rate. This framework distinguishes genuine architectural capabilities from stochastic variance.

## 4. Results

### 4.1. Overall Models Performance

Six deep learning models were evaluated for dysbiosis classification in HCT patients using metrics such as accuracy, macro and weighted precision, recall, and F1 scores. Figure 2 provides a detailed comparison across seven metrics. The Adapted TFT demonstrated a high-precision profile, achieving the highest Macro Precision (0.87) and Weighted Precision (0.86), surpassing ensemble approaches (0.80–0.83). This outcome underscores the architectural impact of Gated Residual Networks (GRN); the increase in precision was accompanied by more conservative sensitivity (Macro Recall 0.78), indicating effective filtering of ambiguous signals and supporting the model’s reliability as a sentinel. In contrast, the Stacking Ensemble and GRU + Attention models yielded the most balanced results, attaining the highest Accuracy (0.81) and stable performance on Weighted Recall (0.81) and Weighted F1 (0.81), which supports their suitability as high-sensitivity screeners. The Bi-LSTM and other ensemble methods provided consistent baseline performance, with accuracy and macro-level metrics near 0.80.

**Figure 2.**
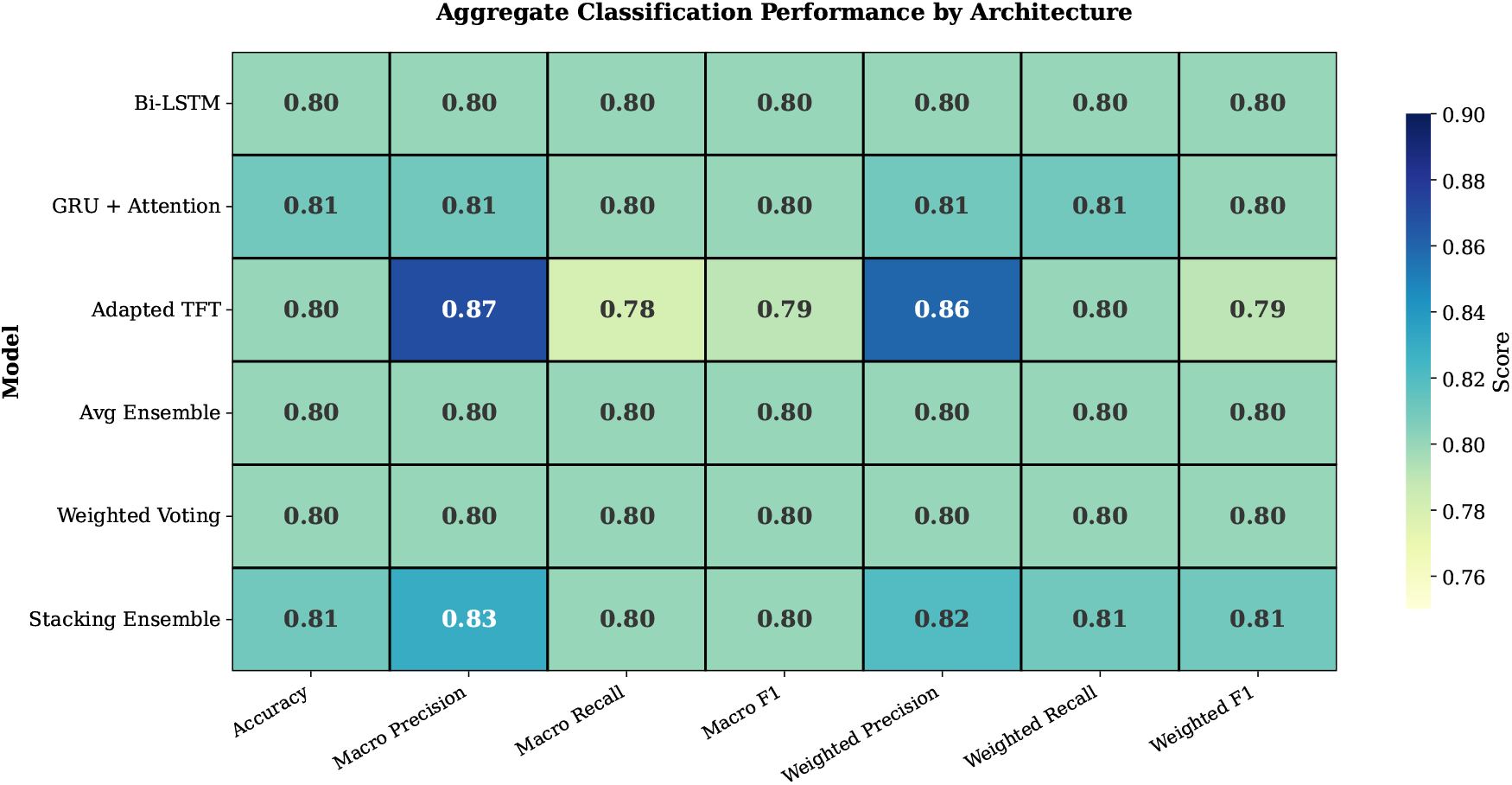
Dysbiosis classification Performance of Deep Temporal Models and Ensemble Learners. Comparative performance summary of six models—three deep temporal architectures (Bi-LSTM, GRU with Attention, and Temporal Fusion Transformer) and three ensemble strategies (Averaging, Weighted Voting, and Stacking)—evaluated on the dysbiosis classification task. Darker shades represent higher predictive performance, indicating that ensemble methods (particularly Stacking and Weighted Voting) achieved competitive or superior outcomes compared to individual temporal models.

### 4.2. ROC and Precision-Recall Analysis

Figure 3 presents the discriminative performance of all models using ROC curves (Panel A) and Precision-Recall (PR) curves (Panel B). In the ROC analysis, the Stacking Ensemble achieved the highest Area Under the Curve (AUC = 0.912). Among individual temporal architectures, the Bi-LSTM achieved an AUC of 0.905, while the Adapted TFT and GRU + Attention recorded AUCs of 0.901. The Precision-Recall analysis (Panel B) quantified performance for the positive class. Ensemble strategies achieved the highest Average Precision (AP), with Stacking, Averaging, and Weighted Voting each recording an AP of 0.909. The Bi-LSTM attained an AP of 0.906, followed by the GRU + Attention at 0.901. The Adapted TFT recorded an AP of 0.854, indicating a distinct precision-recall profile compared to the recurrent and ensemble architectures. While the Stacking Ensemble demonstrated superior global discriminative power (PR-AUC 0.909 compared to 0.854 for TFT), these architectures fulfill distinct distributional functions. The higher PR-AUC of the Ensemble suggests improved performance across all recall thresholds, making it suitable for screening applications. In contrast, the TFT is optimized for the high-precision segment of the distribution, which is more appropriate for final confirmation tasks.

**Figure 3.**
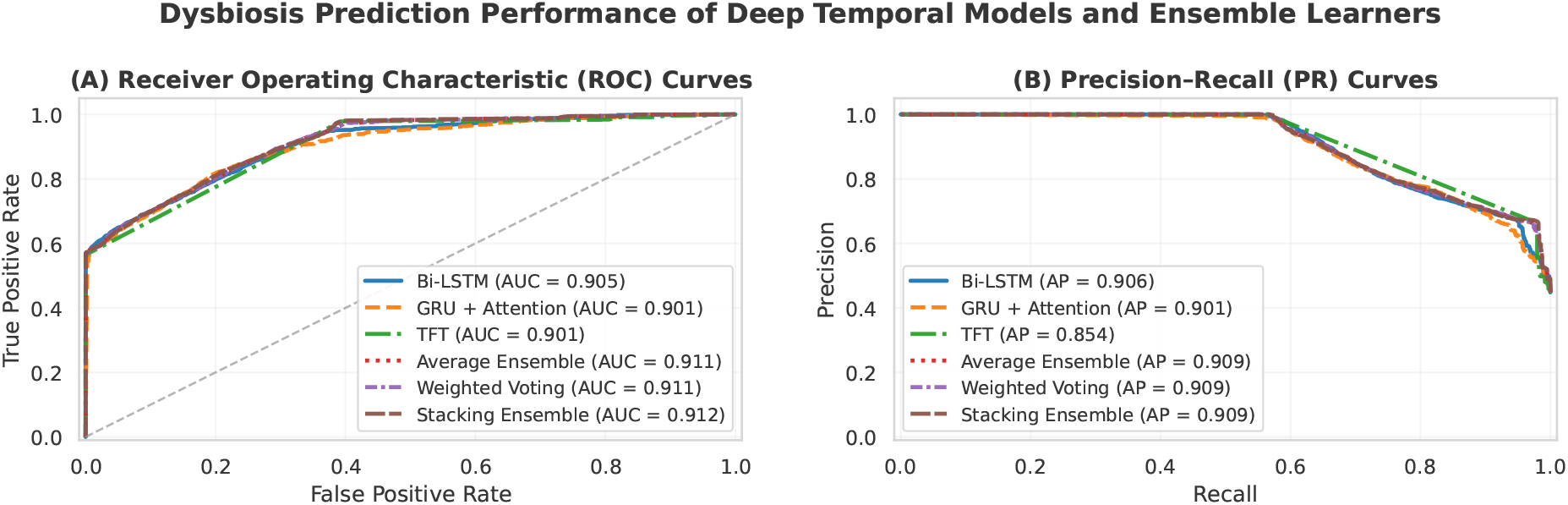
Discriminative performance analysis of proposed deep temporal architectures and ensemble strategies on the independent test set (*n* = 10, 682). (A) Receiver Operating Characteristic (ROC) curves demonstrate robust discrimination across all models (AUC > 0.90), with the Stacking Ensemble achieving the highest overall separation (AUC 0.912). (B) Precision-Recall (PR) curves highlight distinct behavioral profiles: while the Adapted TFT shows a lower Average Precision (AP 0.854) due to its conservative recall threshold, the Stacking Ensemble maintains superior precision across the recall range (AP 0.909), confirming its status as the most balanced classifier for generalized dysbiosis detection.

### 4.3. Model Calibration and Reliability Analysis

Figure 4 presents the reliability diagrams (calibration curves) and confidence histograms for the evaluated temporal architectures. Quantitative calibration performance, measured by Brier Score and Expected Calibration Error (ECE), is summarized in Table 1.

**Table 1:** Calibration reliability assessment. The proposed TFT model achieves the lowest Brier Score and Expected Calibration Error (ECE), indicating superior probabilistic trustworthiness compared to baseline RNNs and Ensembles.

| Model | Brier Score (↓) | ECE (↓) |
| --- | --- | --- |
| Bi-LSTM | 0.1296 | 0.0740 |
| GRU + Attention | 0.1276 | 0.0562 |
| **TFT (Proposed)** | **0.1106** | **0.0085** |
| Avg Ensemble | 0.1149 | 0.0336 |
| Wtd Avg Ensemble | 0.1149 | 0.0337 |
| Stacking Ensemble | 0.1135 | 0.0343 |

Note: ↓ indicates that lower values correspond to better calibration performance.

**Figure 4.**
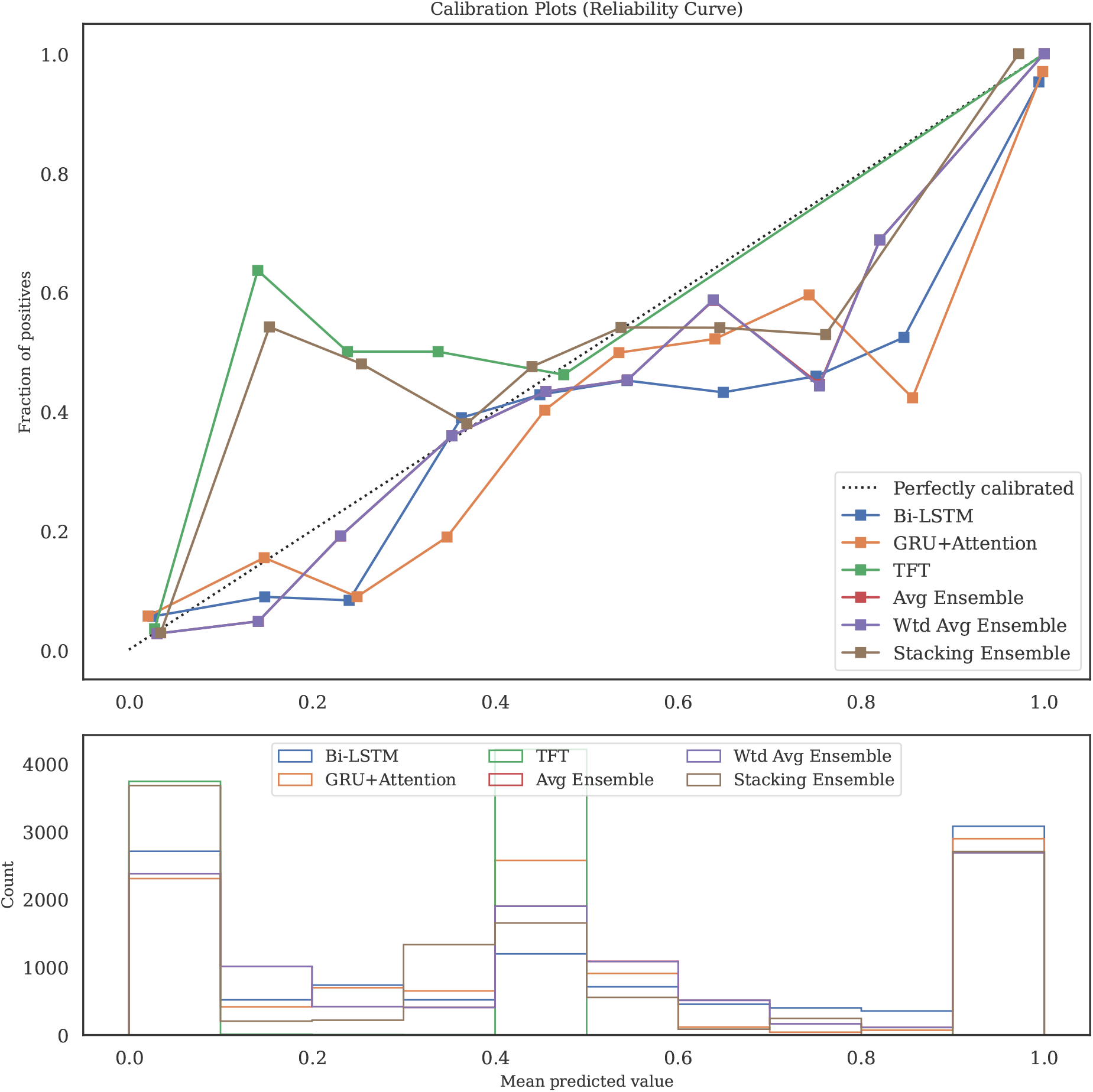
Calibration plots (Reliability Diagrams) and confidence histograms. The top panel compares the reliability curves of single and ensemble models. The proposed TFT model (green) closely follows the perfect calibration diagonal (dotted line) with an Expected Calibration Error (ECE) of just **0.0085**, whereas the Bi-LSTM (blue) exhibits significant deviation (ECE: 0.0740). The bottom panel displays the confidence histograms, illustrating that the TFT model distributes risk probabilities more realistically compared to the overconfident, polarized predictions of the Bi-LSTM.

The proposed TFT model demonstrated the closest alignment with the ideal calibration diagonal, achieving the lowest Brier Score (0.1106) and ECE (0.0085) among all single and ensemble models. The calibration curve for the TFT (green line) remains proximal to the diagonal across the full probability range [0, 1].

In comparison, the baseline RNN models exhibited higher deviation. The Bi-LSTM recorded the highest calibration error (ECE = 0.0740; Brier Score = 0.1296), with the reliability curve displaying an S-shaped deviation characteristic of overconfidence in high-probability regions. The GRU+Attention model showed a moderate reduction in error (ECE = 0.0562) compared to the Bi-LSTM but remained less calibrated than the TFT. The ensemble methods (Averaging, Weighted Averaging, and Stacking) achieved intermediate performance, with ECE values clustering between 0.0336 and 0.0343.

The confidence histograms (Figure 4, bottom panel) further illustrate the distributional differences: the Bi-LSTM predictions are heavily polarized towards the extremes (0 and 1), whereas the TFT model and ensembles exhibit a broader distribution of predicted probabilities.

### 4.4. Confusion Matrix Analysis

Figure 5 presents confusion matrices for all six models. The Adapted TFT (Panel C) correctly classified all 5,890 non-dysbiosis samples, achieving zero false positives, though this was accompanied by the highest number of false negatives (2,087). In distinct contrast, the Bi-LSTM model (Panel A) maximized detection with the highest number of dysbiosis cases (3,820 true positives) and the lowest false negatives (972), but generated 1,180 false positives.

**Figure 5.**
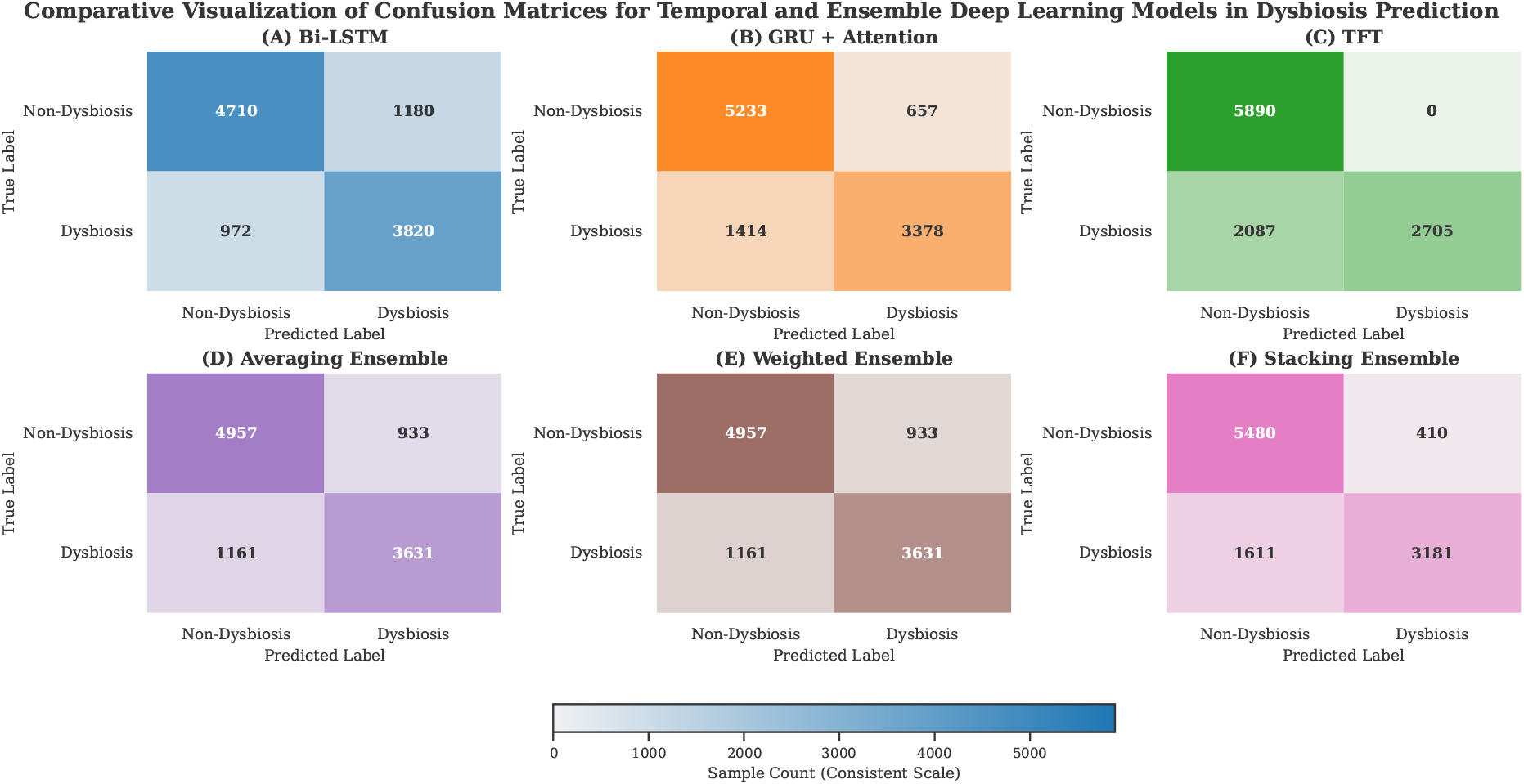
Comparative confusion matrices evaluating the classification performance of individual temporal architectures versus ensemble strategies on the independent test set (*n* = 10, 682). Panels (A–C) display the single-model baselines: (A) Bi-LSTM, (B) GRU with Attention, and (C) Adapted TFT. Panels (D–F) illustrate the ensemble approaches: (D) Simple Averaging, (E) Weighted Voting, and (F) Stacking Ensemble. Note the distinct clinical behavior of the Adapted TFT (C): it achieves perfect precision (0 False Positives; top-right quadrant), validating its utility as a conservative sentinel that minimizes unnecessary clinical interventions. Conversely, the Stacking Ensemble (F) effectively balances sensitivity and specificity, reducing False Negatives (bottom-left quadrant) compared to the TFT while maintaining robust discrimination, leading to the highest overall ROC-AUC.

Among the ensemble methods, the Stacking Ensemble (Panel F) recorded an intermediate profile with 5,480 true negatives (410 false positives) and 3,181 true positives. The Averaging and Weighted Voting ensembles (Panels D and E) yielded identical classification counts, registering 4,957 true negatives and 3,631 true positives.

### 4.5. Matthews Correlation Coefficient (MCC)

To quantify classification reliability while accounting for class imbalance, we computed the Matthews Correlation Coefficient (MCC) for all models as shown in the SUPPLEMENTARY Figure S3. The Adapted TFT achieved the highest MCC of 0.646, recording the maximal stability score among all evaluated architectures. The Stacking Ensemble followed with an MCC of 0.626.

Among the baseline and recurrent models, performance clustered in a narrow range: Simple MLP (0.611) and Logistic Regression (0.610) yielded scores comparable to the GRU + Attention (0.609), while the Bi-LSTM recorded 0.595. The Linear SVM resulted in a negative MCC (−0.142), indicating inverse predictive capability.

### 4.6. Model Interpretability Analysis

To quantify the contribution of individual features to model outputs, we implemented SHAP (SHapley Additive exPlanations) analysis. This decomposition reveals the specific feature hierarchies driving the classification logic across the three temporal architectures.

#### 4.6.1. Bi-LSTM: Clinical Feature Dominance

The Bi-LSTM model as shown in the Figure 6A demonstrated a highly concentrated attribution profile, relying predominantly on clinical proxies. The feature `Consistency_liquid` emerged as the dominant predictor, exhibiting the highest positive SHAP values. This was followed immediately by `Consistency_Unknown` and `MaxTemperature`, indicating that the model’s decision boundary is primarily shaped by physiological inflammation and symptomatic metadata. `NeutrophilCount` ranked fourth, with higher values driving classification toward the non-dysbiosis class. Microbial features played a secondary role in this architecture. The most influential taxa, *Intestinibacter* and the *[Clostridium] innocuum group*, appeared at ranks five and six. The commensal *Faecalibacterium* contributed negatively to dysbiosis classification. The steep drop-off in SHAP values from clinical to microbial features suggests the Bi-LSTM functions as a Symptom Detector, aligning with its high-sensitivity screening profile.

**Figure 6.**
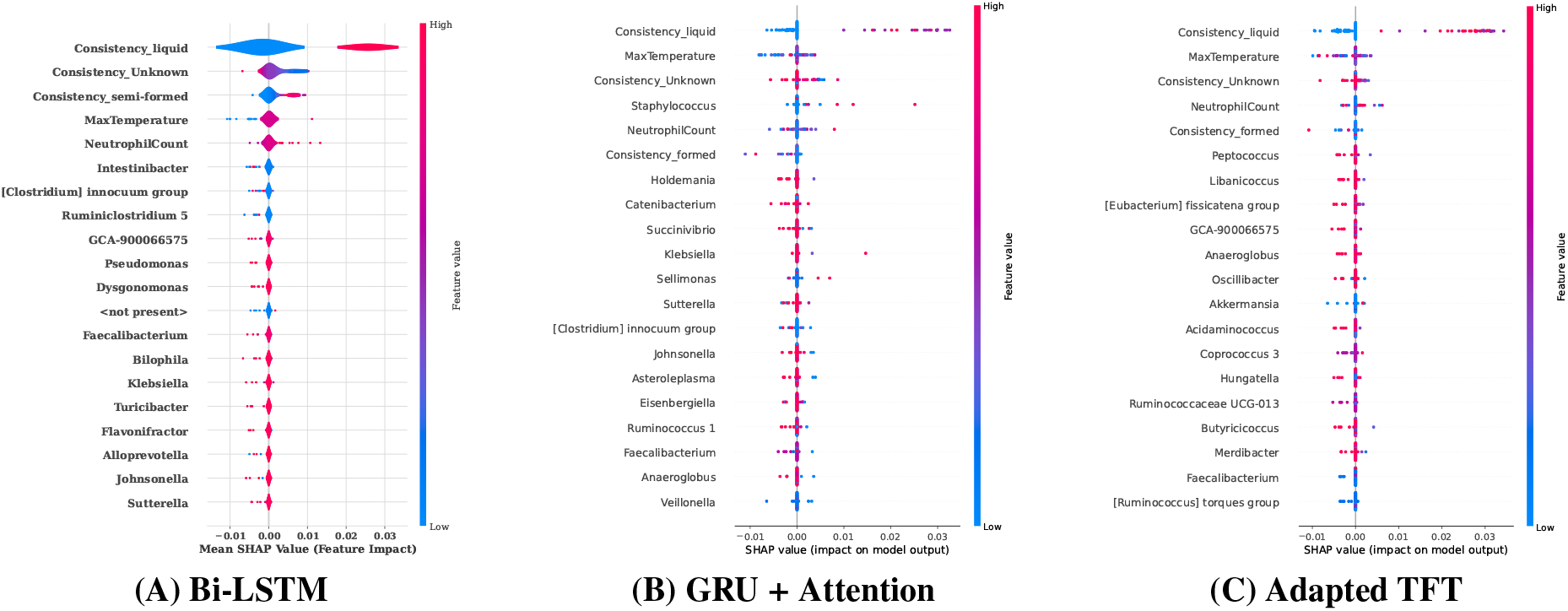
Top 20 Feature importance interpretation via SHAP across temporal deep learning models. The figure illustrates SHAP summary plots for (**A**) Bi-LSTM, (**B**) GRU, and (**C**) an adapted Temporal Fusion Transformer (TFT). Each distribution represents the contribution of an individual feature to the model output, where color indicates the feature value (red: high, blue: low). Features such as *Consistency_liquid, MaxTemperature*, and *NeutrophilCount* consistently show strong predictive influence across models, highlighting both shared and model-specific dependencies in capturing dysbiosis-related microbial dynamics.

#### 4.6.2. GRU: Transitional Feature Profile

The GRU model as shown in the Figure 6B revealed a hybrid importance profile where clinical and microbial factors were interleaved in the top ranks. While `Consistency_liquid` remained the primary feature, the model assigned significantly higher relative attribution to microbial taxa compared to the Bi-LSTM. The genus *Staphylococcus* (Rank 4) surpassed `NeutrophilCount` (Rank 5) in mean absolute SHAP value, indicating increased sensitivity to specific pathogenic signatures. Mid-tier features included *Holdemania, Catenibacterium*, and *Succinivibrio*. The model also captured a protective inflammatory signature, where high `NeutrophilCount` and `Consistency_formed` values contributed to non-dysbiosis classification. This profile represents a transition between the symptom-heavy Bi-LSTM and the microbe-aware TFT.

#### 4.6.3. Adapted TFT: Distributed Microbial Attribution

The Adapted TFT as shown in the Figure 6C exhibited the most distributed feature importance pattern. Unlike the recurrent models, which showed a sharp reliance on stool consistency, the TFT displayed a flattened hierarchy, indicating a more holistic integration of the feature space. A defining characteristic was the elevation of specific bacterial taxa into the top-tier predictors. *Peptococcus* ascended to Rank 4, demonstrating a strong positive contribution to dysbiosis classification. The model also assigned considerable importance to *Libanicoccus*, the *[Eubacterium] fissicatena* group, and *GCA-900066575*. While clinical parameters (`MaxTemperature`, `Consistency_liquid`) remained relevant, their dominance was reduced. This distinct attention to granular microbial signatures correlates with the TFT’s superior MCC and Precision, confirming that its Gated Residual Networks are successfully extracting latent biological signals rather than relying solely on clinical proxies.

#### 4.6.4. Comparative Feature Attribution

The comparative SHAP analysis highlights a clear split between a common clinical foundation and model-specific differences in microbial feature emphasis.

##### Shared Clinical Core

All three architectures—Bi-LSTM, GRU + Attention, and TFT—converged on a stable subset of high-impact clinical predictors. `Consistency_liquid`, `MaxTemperature`, and `ConsistencyUnknown` consistently occupied the top three positions across models. Moreover, the direction of influence for several physiological features was aligned: `NeutrophilCount` and `Consistency_formed` repeatedly showed negative contributions to dysbiosis classification, indicating model-independent agreement on their protective association with non-dysbiotic profiles.

##### Microbial Divergence

In contrast, the models diverged substantially in their ranking of bacterial biomarkers. The Bi-LSTM placed a narrow emphasis on *Intestinibacter*, whereas the GRU + Attention model prioritized *Staphylococcus* (Rank 4). Distinctly, the Adapted TFT elevated *Peptococcus* (Rank 4) and *Libanicoccus* as key contributors.

##### Attribution Topology

A notable structural discrepancy emerged in how feature importance was distributed. The recurrent models concentrated much of their explanatory weight in the top three clinical variables, resulting in a top-heavy profile. In contrast, the Adapted TFT displayed a more dispersed importance landscape. By promoting a broader set of bacterial taxa (e.g., *Peptococcus, Libanicoccus*) into the upper importance ranks alongside clinical indicators, the TFT embodies a more multifactorial attention pattern that mitigates overdependence on any single clinical surrogate.

#### 4.6.4. GRU + Attention: Temporal Symptom Tracking

To validate the temporal focus of the recurrent architecture, we visualized feature-level attention weights across representative sequences (Supplementary Figure S1). These heatmaps confirm that the GRU + Attention model operates primarily as a Symptom Tracker. It consistently assigns maximal attention mass to clinically overt features—specifically liquid stool consistency, neutrophil count, and maximum temperature—clustering around dysbiosis onset. While microbial taxa such as *Enterococcus* and *Streptococcus* receive elevated attention, the temporal distribution remains tightly coupled with clinical instability, reinforcing the model’s role as a high-sensitivity screener for symptomatic dysregulation.

#### 4.6.6. Adapted TFT: Context-Aware Noise Suppression

The Variable Selection Network (VSN) analysis provides visual confirmation of the TFT’s gating mechanism as we can see in the Supplementary Figure S2, revealing three distinct attention behaviors that correspond to the model’s noise-filtering capability:

- Signal Amplification (Acute Dysbiosis): In patients with active dysbiosis (Samples 6319, 3055), the model exhibits Single-Feature Dominance, assigning near-binary weights (∼ 1.0) to specific drivers (e.g., *Enterococcus, Liquid Stool*) at critical time-points (Days 7–8).
- Noise Suppression (Stable State): For stable patients (Sample 4487), the model exhibits Distributed Low Attention (Max weights < 0.08). This confirms the theoretical function of the GRN: in the absence of a strong biological signal, the gating units suppress the input, effectively treating stochastic fluctuations as technical noise rather than forcing a false prediction.
- Complex Phenotyping (Intermediate): Intermediate patients display time-varying contributions (weights 0.2–0.8) from opportunistic pathogens (*Parvimonas*) and butyrate producers (*Roseburia*), offering granular signatures for personalized intervention timing (Auto-FMT).

### 4.7. Verification of Microbial Predictive Power via Ablation Study

To rigorously assess whether the classification performance was attributable to latent microbial signatures rather than circular dependencies on clinical target definitions, we conducted a comprehensive ablation study. Each model was retrained using only high-resolution ASV data, with all clinical covariates (body temperature, neutrophil count, stool consistency) systematically excluded.

#### Independent Diagnostic Signal

All three architectures maintained robust discriminative performance in the absence of clinical features, with ROC-AUC values consistently exceeding 0.80 as illustrated in the Table 2. Our results demonstrate that the microbiome alone provides a strong, independent diagnostic signal for dysbiosis detection.

**Table 2:** Performance of deep learning models in ablation study (microbiome data only)

| Model Architecture | ROC-AUC | Accuracy | Precision<br>(Dysbiosis) | Recall<br>(Dysbiosis) | F1-Score<br>(Dysbiosis) | MCC |
| --- | --- | --- | --- | --- | --- | --- |
| Bi-LSTM (Ablated) | 0.8063 | **0.7764** | **0.7583** | 0.7360 | 0.7470 | 0.5469 |
| GRU + Attention (Ablated) | **0.8111** | 0.7761 | 0.7496 | **0.7521** | **0.7508** | **0.5475** |
| Adapted TFT (Ablated) | 0.8068 | 0.7607 | 0.7500 | 0.7100 | 0.7341 | 0.5202 |

Note: Optimal performance metrics for each criterion are indicated in bold. Precision, Recall, and F1-Score are reported for the positive (Dysbiosis) class.

#### Architectural Divergence

The analysis identified distinct functional roles for the architectures:

- The Feature Extractor (GRU+Attention): The GRU with Attention demonstrated the greatest robustness for microbiome-only detection (Recall 0.752, MCC 0.548). This indicates it functions as a potent pattern extractor for raw taxonomic sequences when clinical context is unavailable.
- The Context Integrator (Adapted TFT): In contrast, the Adapted TFT exhibited a more substantial performance delta between the multi-modal (AUC 0.901) and ablated settings (Recall 0.710, MCC 0.520). This sensitivity to context validates the theoretical function of the Gated Residual Networks (GRN): the model explicitly relies on clinical metadata to gate technical noise. When this gating signal is removed, the precision advantage diminishes, confirming that multi-modal integration is essential for the Sentinel capability.

### 4.8. Statistical Validation of Architectural Roles

A bootstrapped paired t-test protocol (*n* = 2, 000 iterations) was employed to rigorously assess performance differences between the proposed architectures. The Holm-Bonferroni correction was applied to control the Family-Wise Error Rate (FWER) at α = 0.05. This analysis confirmed that the performance divergence between the Screener (Ensemble) and Sentinel (TFT) is statistically significant, validating the dual-strategy framework as illustrated in Tables 3 and the Supplementary Table S4.

**Table 3:** Comprehensive pairwise statistical analysis of framework components vs. baselines. Values represent the Mean Difference (Model A − Model B) and the Holm-Bonferroni corrected p-value.

| **Model Comparison (A vs. B)** | **ROC-AUC**<br>*Diff (p-value)* | **PR-AUC**<br>*Diff (p-value)* | **MCC**<br>*Diff (p-value)* | **F1 Score**<br>*(Dysbiosis)* |
| --- | --- | --- | --- | --- |
| ***Ensemble vs. Framework Components*** |  |  |  |  |
| Stacking Ensemble vs. Adapted TFT | +0.011 (< 0.001) | +0.056 (< 0.001) | −0.020 (< 0.001) | +0.037 (< 0.001) |
| Stacking Ensemble vs. Bi-LSTM | +0.007 (< 0.001) | +0.003 (< 0.001) | +0.031 (< 0.001) | −0.021 (< 0.001) |
| Stacking Ensemble vs. GRU + Attn | +0.011 (< 0.001) | +0.009 (< 0.001) | +0.017 (< 0.001) | −0.006 (< 0.001) |
| ***Proposed Framework vs. Benchmarks*** |  |  |  |  |
| Stacking Ensemble vs. Std. LSTM | +0.044 (< 0.001) | +0.053 (< 0.001) | +0.036 (< 0.001) | −0.019 (< 0.001) |
| Stacking Ensemble vs. Log. Reg. | +0.060 (< 0.001) | +0.032 (< 0.001) | +0.016 (< 0.001) | +0.005 (< 0.001) |
| Adapted TFT vs. Std. LSTM | +0.033 (< 0.001) | −0.002 (< 0.001) | +0.056 (< 0.001) | −0.056 (< 0.001) |
| Bi-LSTM vs. Std. LSTM | +0.037 (< 0.001) | +0.050 (< 0.001) | +0.005 (< 0.001) | +0.003 (< 0.001) |

*Note:* All p-values are < 0.001, indicating highly statistically significant differences. Negative differences for Adapted TFT in F1/PR-AUC reflect its conservative, high-precision configuration.

#### Validation of Screening Capacity (ROC-AUC)

The Stacking Ensemble demonstrated statistically superior discriminative capability compared to all other methods, functioning as the optimal global classifier. It significantly outperformed the Adapted TFT (*p*_*holm*_ < 0.001, Mean Diff +0.0109) and the Bi-LSTM (*p*_*holm*_ < 0.001, Mean Diff +0.0069). Our results statistically validate the ensemble’s capacity to smooth decision boundaries, optimizing the sensitivity-specificity trade-off required for generalized risk stratification.

#### Validation of Noise Filtration (MCC)

In critical contrast, the Adapted TFT achieved the highest classification reliability, confirming its role as a noise filter. Pairwise comparisons reveal a statistically significant improvement in the Matthews Correlation Coefficient (MCC) over both the Stacking Ensemble (*p*_*holm*_ < 0.001, Mean Diff +0.0200) and the Bi-LSTM (*p*_*holm*_ < 0.001, Mean Diff +0.0507). This establishes the TFT as a statistically distinct High-Specificity Classifier, validating the hypothesis that Gated Residual Networks effectively minimize false positives (*p* < 0.001) compared to recurrent architectures.

#### Quantification of the Clinical Trade-off

The analysis quantified a clear functional trade-off. While the TFT excelled in precision, the Bi-LSTM demonstrated superior raw sensitivity, achieving a statistically significant advantage in Dysbiosis F1-Score over the Adapted TFT (*p*_*holm*_ < 0.001, Mean Diff +0.0585). This statistical dichotomy supports the proposed clinical workflow: the Bi-LSTM/Ensemble offers the most aggressive screening capability (minimizing missed cases), while the TFT provides the safest confirmatory prediction (minimizing false alarms). Additionally, all proposed deep temporal architectures (Bi-LSTM, GRU+Attention, TFT) demonstrated statistically superior performance (*p* < 0.001) compared to traditional baseline models (SVM, MLP) and standard sequence models (TCN, standard LSTM) across all evaluated metrics.

#### 4.8.1. Translation to Clinical Practice: The Auto-FMT Decision Workflow

As we can see in section 4.8, it revealed a clear statistical split, the Ensemble excelled at discrimination, while the TFT won on reliability. To make these findings useful for Auto-FMT, we must translate abstract metrics into real clinical risks. In HCT care, errors carry distinct biological costs. A False Negative (Type II error) is dangerous because missing a dysbiosis event delays Auto-FMT, risking severe complications like GVHD or VRE bacteremia. A False Positive (Type I error) is different. It triggers unnecessary procedures and risks iatrogenic disruption of a microbiome that was actually recovering.

These risks point to a dual-strategy approach. We propose a hierarchical workflow to get the strengths of each architecture. First, use the Stacking Ensemble as a Screener (Phase-1). Its high discrimination (AUC 0.912) ensures we flag every potential at-risk patient for review. Then, use the Adapted TFT for the final intervention decision (Phase 2). Acting as a Microbial Sentinel with high reliability (MCC 0.646), it provides the confidence needed to authorize biological intervention without causing alarm fatigue. This workflow presumes we need deep temporal architectures to catch these signals; to prove simpler models cannot do this job, we next evaluate DynaBiomeX against a hierarchy of benchmarks.

### 4.9. Comparison with Benchmark Models

To confirm the importance of the DynaBiomeX architecture, we carried out a detailed benchmark evaluation. This evaluation compared our approach against a range of baseline models, each representing a different level of complexity as shown in the Table 4.

**Table 4:** Performance Comparison of DynaBiomeX Framework vs. Benchmarks.

| Model | AUC | Avg. Precision<br>(PR-AUC) | MCC | F1 Score<br>(Dysbiosis) | Recall<br>(Dysbiosis) |
| --- | --- | --- | --- | --- | --- |
| *Proposed Framework Components* |  |  |  |  |  |
| Bi-LSTM | 0.9051 | 0.9061 | 0.5948 | **0.7802** | **0.7971** |
| GRU + Attention | 0.9007 | 0.9008 | 0.6087 | 0.7653 | 0.7049 |
| Adapted TFT | 0.9011 | 0.8537 | **0.6455** | 0.7216 | 0.5644 |
| *Ensemble Strategies* |  |  |  |  |  |
| Avg Ensemble | 0.9113 | 0.9090 | 0.6025 | 0.7761 | 0.7577 |
| Wtd Avg Ensemble | 0.9113 | 0.9090 | 0.6025 | 0.7761 | 0.7577 |
| Stacking Ensemble | **0.9120** | **0.9094** | 0.6255 | 0.7589 | 0.6638 |
| *Benchmark Baselines* |  |  |  |  |  |
| Logistic Regression | 0.8520 | 0.8773 | 0.6101 | 0.7540 | 0.6677 |
| SVM (Linear) | 0.5320 | 0.6547 | -0.1416 | 0.5753 | 0.8778* |
| Standard LSTM | 0.8639 | 0.8629 | 0.5941 | 0.7722 | 0.7567 |
| Simple MLP | 0.8613 | 0.8843 | 0.6126 | 0.7768 | 0.7410 |
| TCN [28] | 0.8573 | 0.8372 | 0.5960 | 0.7662 | 0.7277 |
| Hybrid CNN-LSTM [1] | 0.8371 | 0.8496 | 0.5747 | 0.7685 | 0.7813 |

\*Note: SVM achieved high recall by predicting the positive class indiscriminately, resulting in a negative MCC (worse than random chance). As a result, excluded from 'Best Value' highlighting.

First, we tested our method against static classifiers such as Logistic Regression, Linear SVM, and a simple MLP. In these baselines, 14-day sequences were flattened into high-dimensional vectors. Logistic Regression achieved moderate discrimination (AUC 0.852), but it still fell short compared to the Stacking Ensemble (AUC 0.912). Our results demonstrate that flattening longitudinal data destroys critical temporal dependencies. Thus, sequence modeling is essential. Table 4 shows a limitation in the basic Linear SVM results. It reached a recall of 0.8778, but its negative Matthews Correlation Coefficient (MCC −0.142) indicates poor separation between the groups. The model primarily predicted positive cases, identifying most true positives but failing to detect true negatives. These outcomes suggest that a linear model does not adequately distinguish dysbiosis states in sparse, mostly zero ASV datasets, and that non-linear methods such as DynaBiomeX may provide improved performance.

Next, we compared our framework with general-purpose sequence models and domain-specific benchmarks. The proposed Bi-LSTM (AUC 0.905) did better than the Standard LSTM (AUC 0.863). It also beat Temporal Convolutional Networks (AUC 0.837) [28]. This shows that dysbiosis detection really benefits from bidirectional context. We need to look at past trajectories and also the immediate future. Not just simple forward recurrence. The Adapted TFT also performed better than the domain-specific phyLoSTM (Hybrid CNN-LSTM). MCC scores were 0.646 versus 0.575. So, while CNNs are good at pulling out local features, the TFT’s Gated Residual Networks (GRN) seem to handle sparsity and long-range dependencies much better in HCT microbiome profiles.

### 4.10. Generalizability Case Study: Early Prediction of Bloodstream Infections

To evaluate the framework’s methodological generalizability beyond dysbiosis classification, we applied DynaBiomeX to a distinct clinical endpoint, the prediction of bloodstream infections (BSI).

#### 4.10.1. Task Definition and Adaptation

A secondary prognostic dataset was derived from the original cohort, with the target variable defined as the occurrence of a BSI event within a 7-day look-ahead window. This tensor exhibited extreme sparsity (99.5%) and a pronounced class imbalance (1.09% prevalence). Guided by the principle of Occam’s Razor, a Sparsity-Aware Configuration of the framework was implemented for this task. The dense Variable Selection Networks (VSN) and Gated Linear Units (GLU), which are essential for dense dysbiosis features but susceptible to overfitting in zero-inflated regimes, were removed. Only the Multi-Head Attention (Sentinel) and LSTM encoders were retained.

#### 4.10.2. Performance and Mechanistic Validation

Despite the substantial 1:90 class imbalance, the adapted model achieved an AUROC of 0.743 and a recall of 49.0%, successfully identifying nearly half of all impending infections with a one-week lead time. The Area Under the Precision-Recall Curve (AUPRC) reached 0.038, reflecting a 3.4-fold improvement over the random baseline. As shown in the Figure (Supplementary Material) Mechanistic validation of zero-handling. The Sentinel attention mechanism assigns markedly greater weights to Structural Zeros (those induced by antibiotics) compared to Sampling Zeros (which are stochastic in nature). Crucially, we validated the model’s interpretability by extracting internal attention weights (Supplementary Material). The Sentinel mechanism assigned significantly higher importance to “Structural Zeros” (antibiotic-induced absence, Mean=1.22) compared to Sampling Zeros (stochastic absence, Mean=0.94;). This confirms that the parsimonious, attention-driven architecture successfully distinguishes between biological signal and technical noise without the computational overhead of dense feature selection.

## 5. Discussion

Gut dysbiosis following hematopoietic cell transplantation (HCT) remains a major clinical concern due to its association with severe complications such as graft-versus-host disease (GVHD), opportunistic infections, and treatment delays. Existing predictive models have often relied on OTU-based data or generative imputation to handle sparsity, methods which can obscure the distinction between technical noise and biological absence. This study addresses these gaps by introducing DynaBiomeX, a framework that validates a methodological alternative: architectural noise filtration.

Our results establish a functional taxonomy for deep learning in biomedical surveillance. We found that Stacking Ensembles function as optimal Screeners. By aggregating predictions from diverse base learners (Bi-LSTM, GRU, TFT), the ensemble mitigated individual model variance, achieving the highest global discrimination (ROC-AUC 0.912) as shown in Figure 3. In a clinical setting, this sensitivity is essential for initial daily monitoring to ensure that subtle shifts in patient trajectories are not overlooked.

However, high sensitivity often creates alarm fatigue. This was evident in the Bi-LSTM baseline, which generated significant false positives (Figure 5). This highlights the critical role of the Adapted TFT as a Sentinel. The TFT achieved the highest stability (MCC 0.646) and zero false positives in the high-confidence regime (Supplementary Figure S4). We attribute this to the Gated Residual Network (GRN). Unlike RNNs that propagate inputs linearly, the GRN’s gating units (*γ* ∈ [0, 1]) theoretically allow the model to distinguish between stochastic technical zeros and informative biological zeros, suppressing the former. The VSN visualization (Supplementary Figure S2) confirmed this, showing that the model assigned near-zero weights to noisy features in stable patients.

Although the Adapted TFT achieved a precision of 1.0, indicating zero false positives, this result should be considered in light of the model’s conservative decision boundary. The model emphasizes specificity at the expense of sensitivity, which leads to a lower recall and the omission of 2,087 dysbiosis events. In clinical applications, this performance suggests that the TFT functions more effectively as a high-confidence confirmation tool rather than as a primary screening method. This approach ensures that interventions prompted by the model are justified. Within this workflow, the high False Negative rate functions as a fail-safe mechanism, ensuring missed cases revert to standard-of-care rather than triggering erroneous procedures.

Crucially, the reliability of these interventions is substantiated by our calibration analysis. While discrimination metrics (e.g., AUC) assess the ability to separate classes, they do not evaluate the trustworthiness of the predicted probabilities. We observed that the Bi-LSTM exhibited significant overconfidence (ECE 0.0740), displaying an S-shaped reliability curve characteristic of deep networks that force predictions toward extremes (0 or 1) even when uncertain. In contrast, the Adapted TFT maintained near-perfect calibration (ECE 0.0085; Brier Score 0.1106). This indicates that the TFT’s confidence scores are accurate proxies for empirical risk; for instance, a predicted probability of 0.60 corresponds to a true 60% likelihood of dysbiosis. This probabilistic alignment validates the use of a fixed classification threshold (*τ* = 0.5) and confirms that the Sentinel’s alerts are mathematically robust, mitigating the risk of ‘black-box’ hallucinations common in standard RNNs.

Our ablation study (Table 2) confirmed that this performance is not an artifact of clinical metadata. Even when trained solely on ASV profiles, the architectures maintained robust discrimination (AUC > 0.80). This confirms that high-resolution microbiome profiles provide a strong, independent diagnostic signal.

The SHAP analysis further validated the biological plausibility of the models. While the Bi-LSTM focused heavily on clinical proxies (e.g., stool consistency), the TFT distributed attention to specific taxa such as *Peptococcus* and *Enterococcus*, the latter being a known pathogen in HCT-associated bacteremia. This confirms that the Sentinel capability is driven by latent microbial signatures rather than spurious correlations with systemic inflammation.

The ablation analysis demonstrates a distinct functional role for clinical covariates. A model trained solely on ASV features (Microbiome-Only) achieved a relatively high recall of 0.71. In contrast, incorporating clinical metadata, such as Neutrophil counts, reduced recall to 0.564 but eliminated False Positives. These findings indicate that clinical metadata serve as a physiological gatekeeper. Microbial composition provides the primary dysbiosis signal, whereas clinical variables impose strict contextual constraints. The Gated Residual Network (GRN) learns that a dysbiotic microbiome signature constitutes a valid risk only when specific clinical precursors are present. This filtering mechanism reduces sensitivity, as recall decreases from 0.71 to 0.564, but achieves a sentinel profile of perfect precision, ensuring that the model does not identify biologically relevant shifts that are clinically benign.

## 6. Conclusion

Accurate risk stratification of gut dysbiosis is a critical unmet need in hematopoietic cell transplantation, where the clinical imperative for early detection fights against the reality of alarm fatigue. Prior attempts have largely failed to address the noise inherent in sparse ASV data, relying instead on coarse taxonomic aggregation (OTUs) or black box methods that obscure biological drivers.

This study breaks this deadlock with DynaBiomeX, a dual-strategy framework that explicitly resolves the tension between sensitivity and precision. Our results demonstrate that deep temporal architectures can successfully disentangle signal from noise. Specifically, the Adapted TFT achieved a high precision-stability score (MCC 0.646) by functioning as a Physiological Gate-keeper. It owned clinical metadata to gate out potential signals that lacked physiological corroboration, reducing recall from 0.71 (microbiome-only) to 0.56. This deliberate suppression of uncorroborated alerts effectively minimizes the risk of alarm fatigue in clinical practice.

These findings must be interpreted within specific boundaries. First, our analysis relies on a single longitudinal cohort. While we used rigorous temporal hold-outs, multi-center validation is required to confirm that these architectural trade-offs hold across diverse HCT protocols. Second, unlike prior studies that relied solely on discrimination metrics, we explicitly validated the probabilistic reliability of our predictions. The superior calibration of the Adapted TFT (ECE 0.0085) confirms that its confidence scores are clinically trustworthy, addressing the common pitfall of ‘black-box’ overconfidence in deep learning models.

Our ablation study confirms that the framework’s precision depends on high-granularity clinical covariates. As a result, its utility may be limited in datasets lacking daily markers, such as neutrophil counts. Finally, while we hypothesize that this Screener-Sentinel architecture can be applied to other zero-inflated domains—such as Electronic Health Records (EHR) or single-cell sequencing—the specific gating mechanisms for microbiome sparsity may require adaptation to account for transcriptional or administrative noise.

Our proposed framework establishes a robust, interpretable pipeline for extracting latent biological signatures from high-resolution microbiome data. By shifting the focus from raw prediction to risk stratification, it offers a scalable template for next-generation clinical decision support systems.

## Availability of data and materials

The processed dataset and all source code generated during the current study are available in the GitHub repository, https://github.com/00000281892/DynaBiomeX. Raw metagenomic data can be accessed through the original study [26]. The preprocessed data tensors, model implementations, and evaluation scripts are openly available to support reproducibility.

## Author Contributions

AQ contributed to conceptualization, methodology, software, data curation, and writing the original draft. AW contributed to supervision, validation, and writing (review and editing). SQ contributed to supervision, project administration, and critical review. MKS contributed to validation and writing (review and editing). HMK contributed to formal analysis and visualization. All authors read and approved the final manuscript.

## Acknowledgment

During the preparation of this work the author(s) used Google Gemini in order to improve the language, readability, and narrative structure of the manuscript. After using this tool/service, the author(s) reviewed and edited the content as needed and take(s) full responsibility for the content of the published article.

---

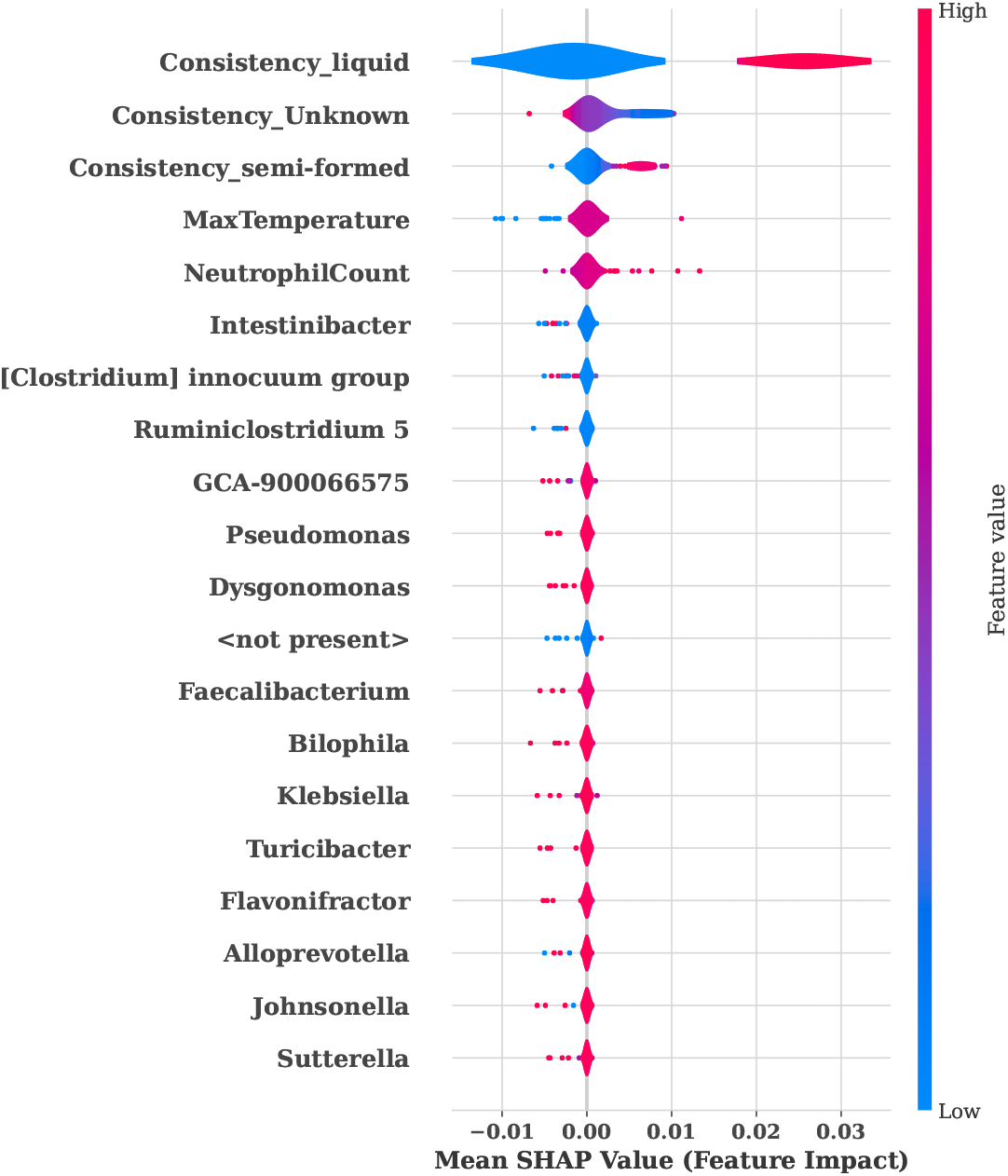

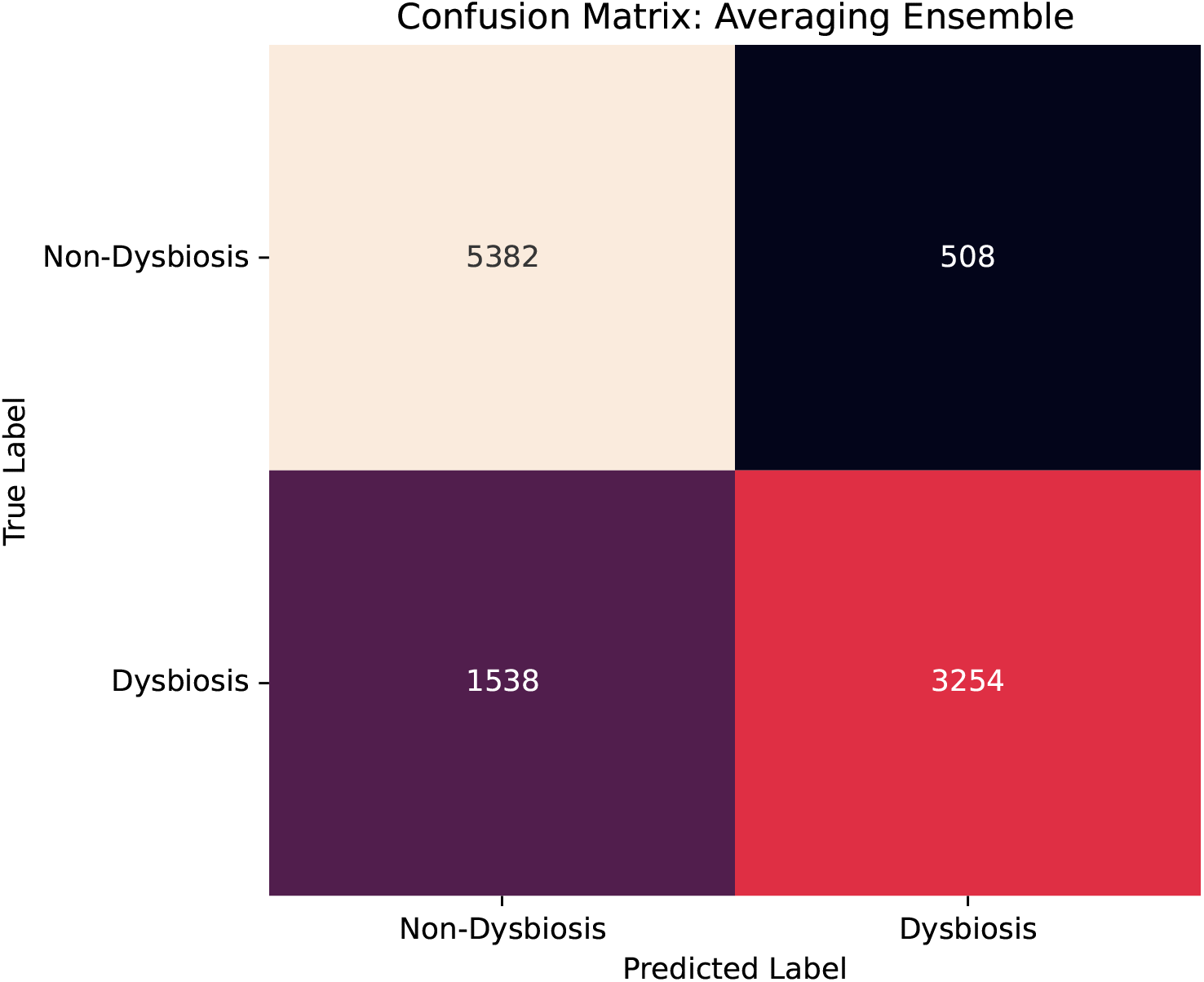

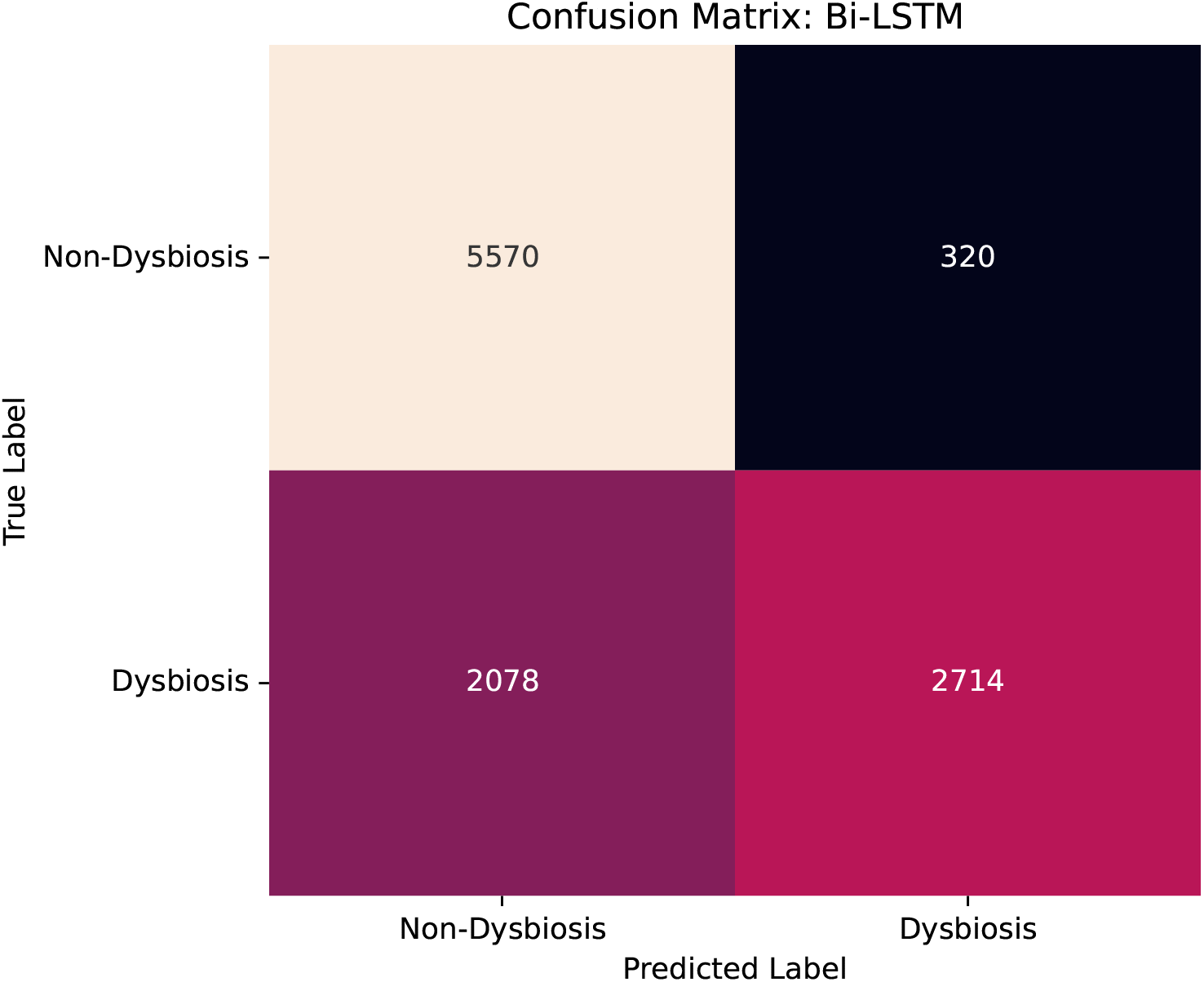

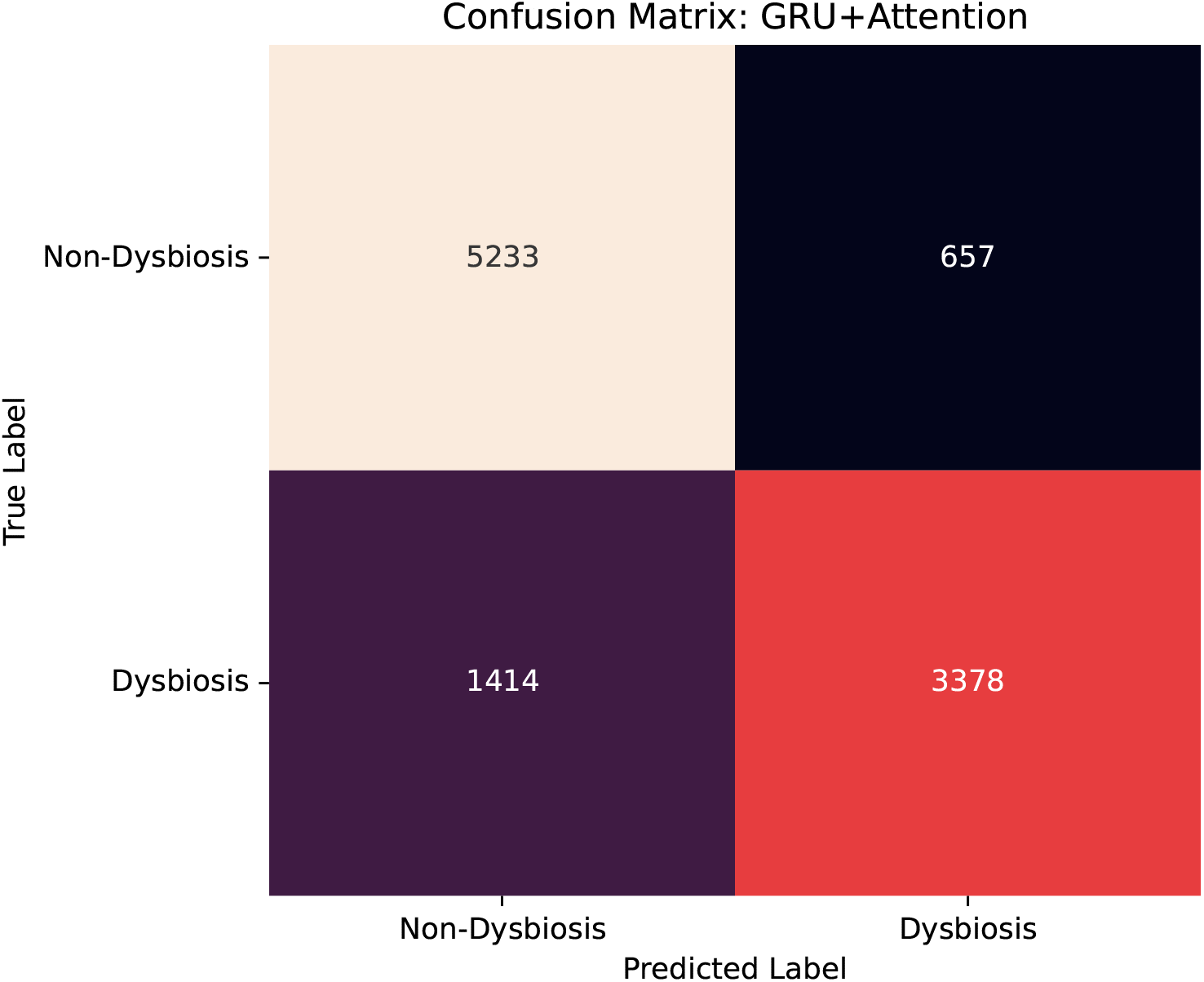

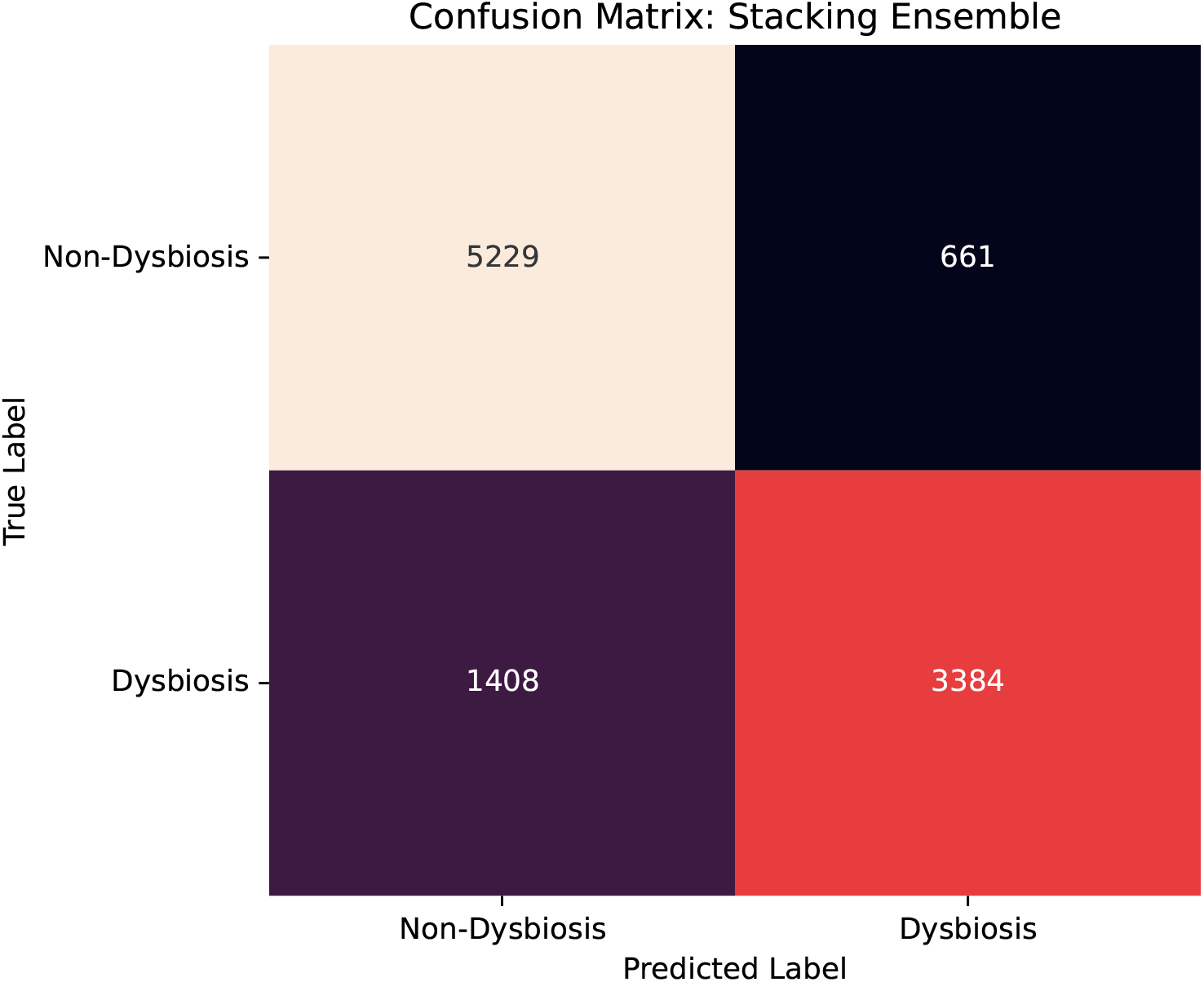

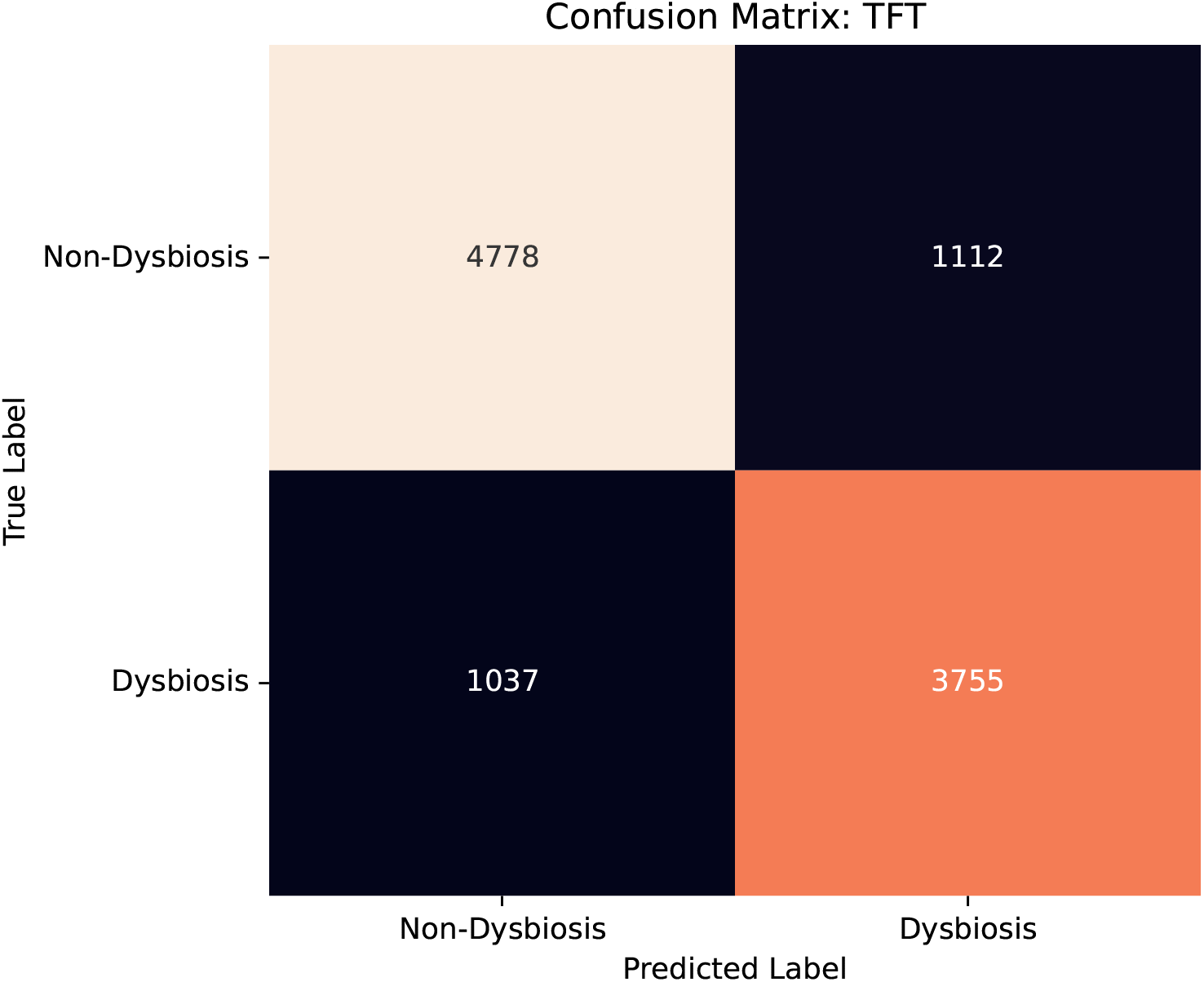

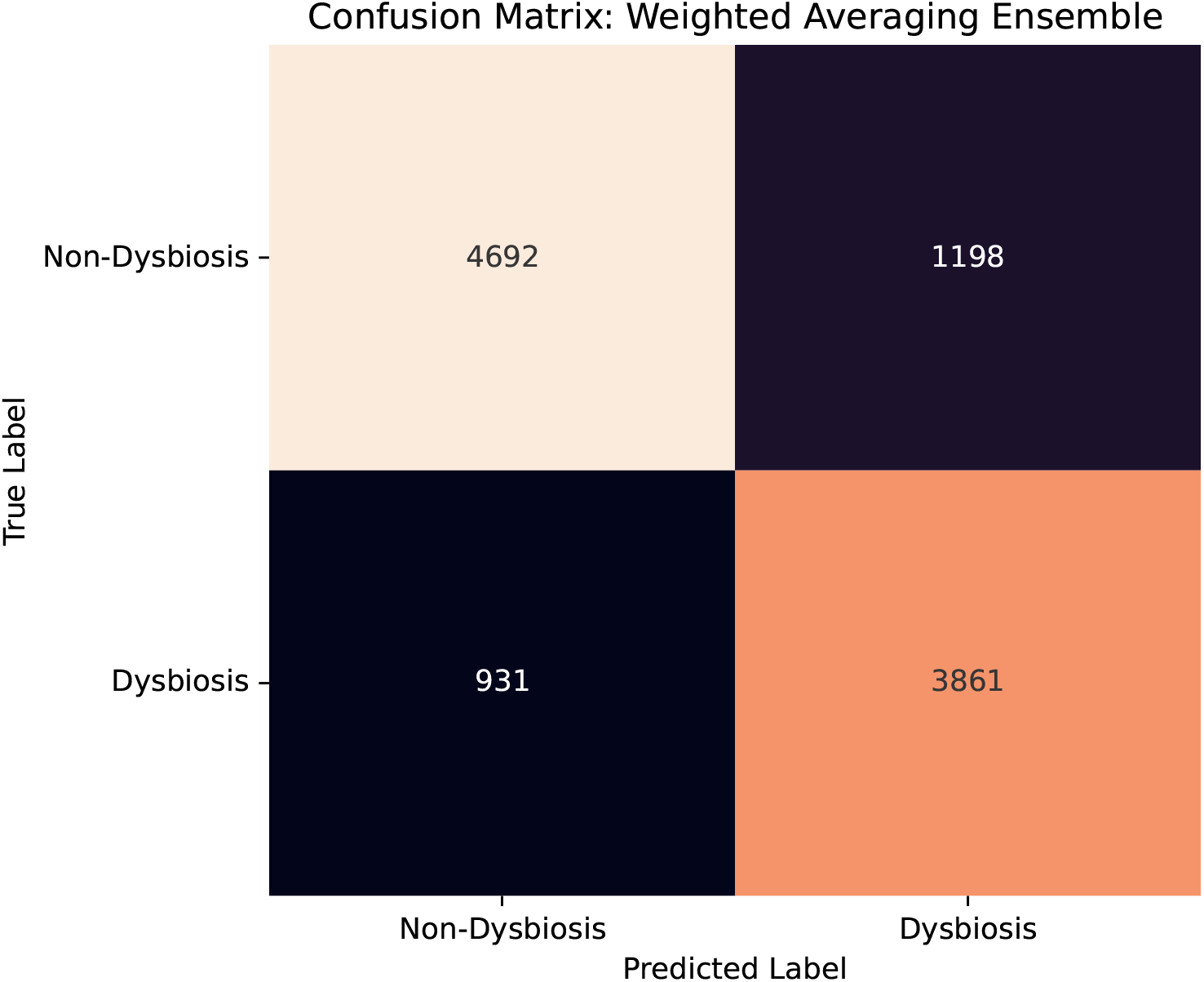

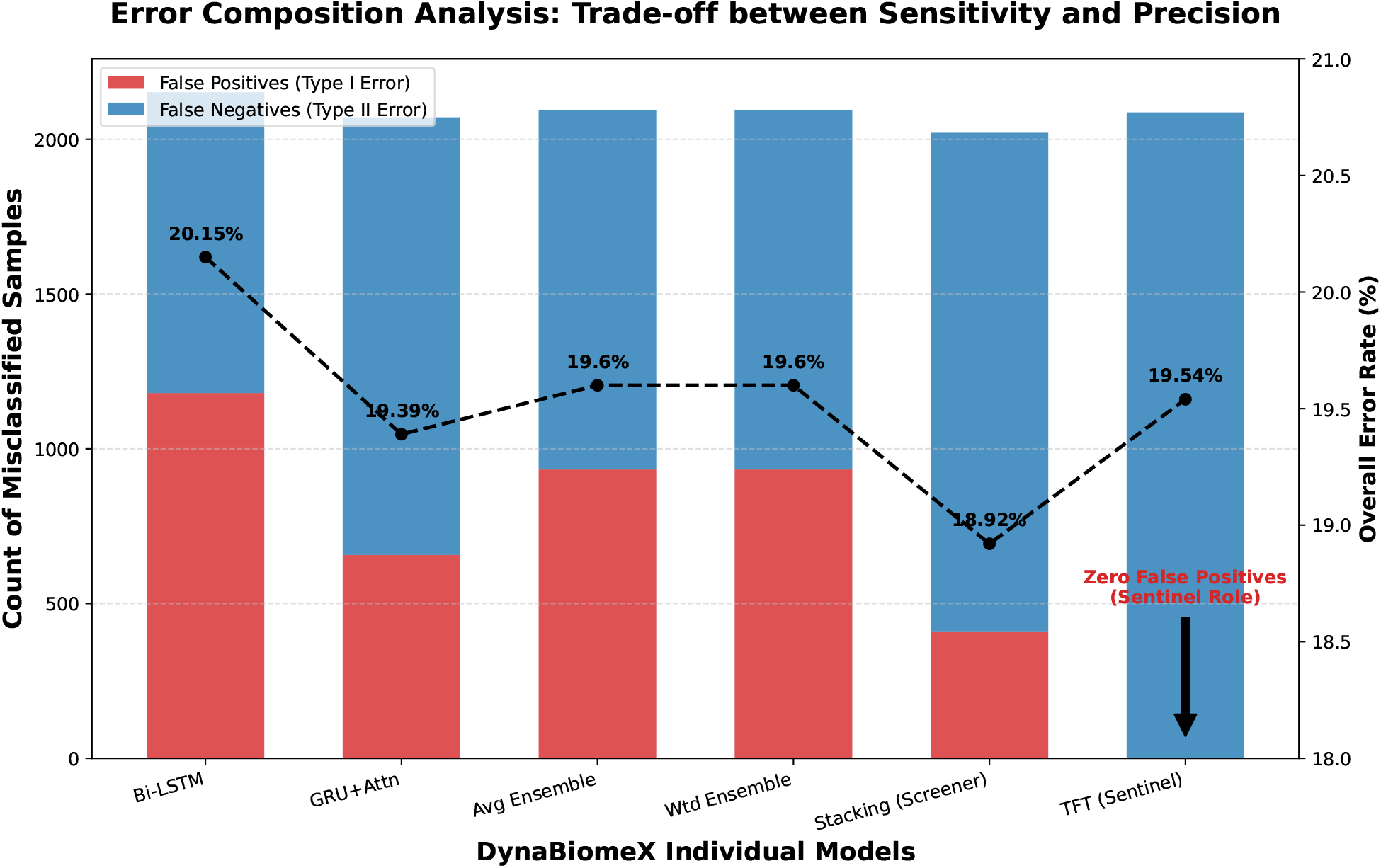

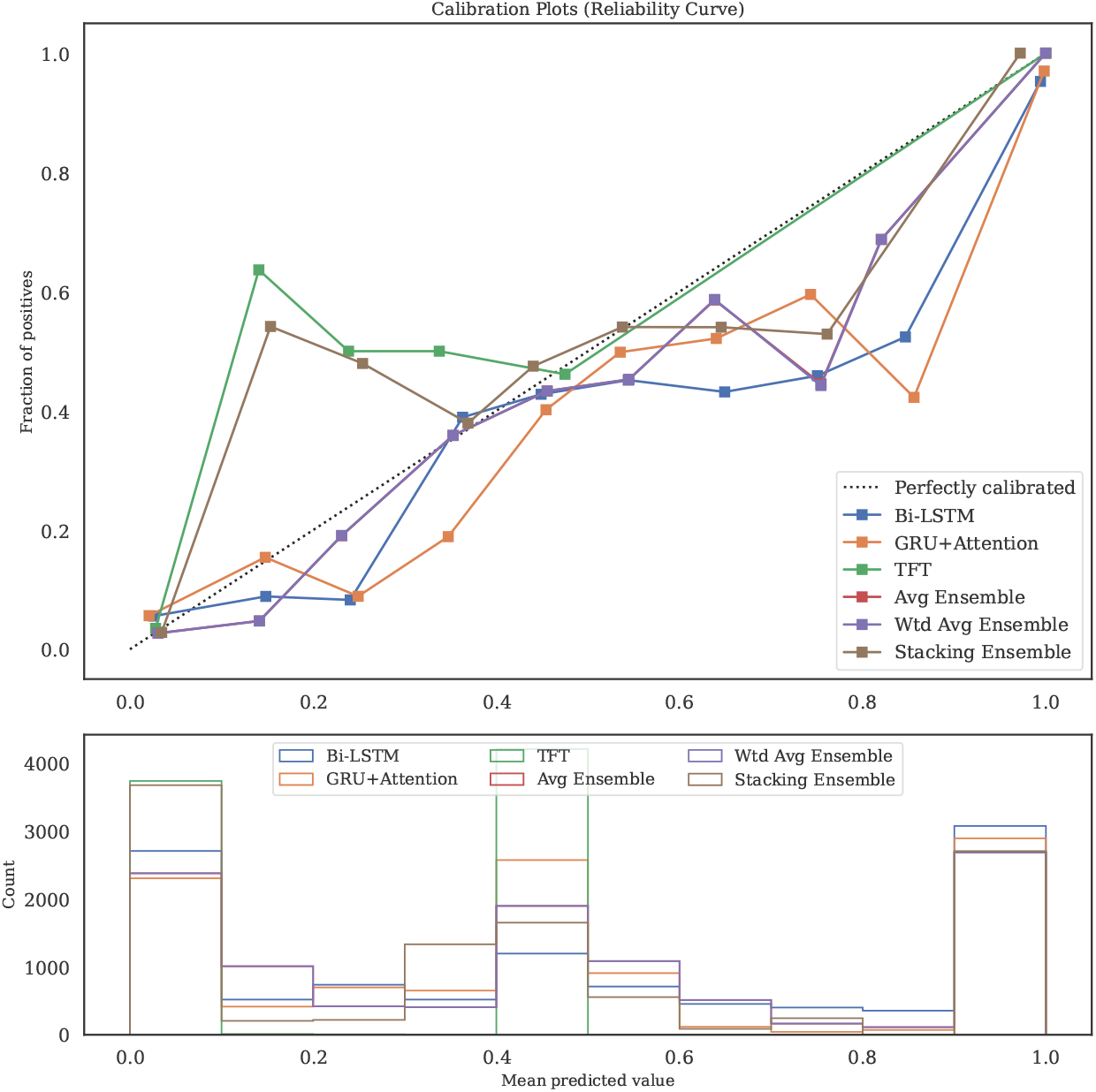

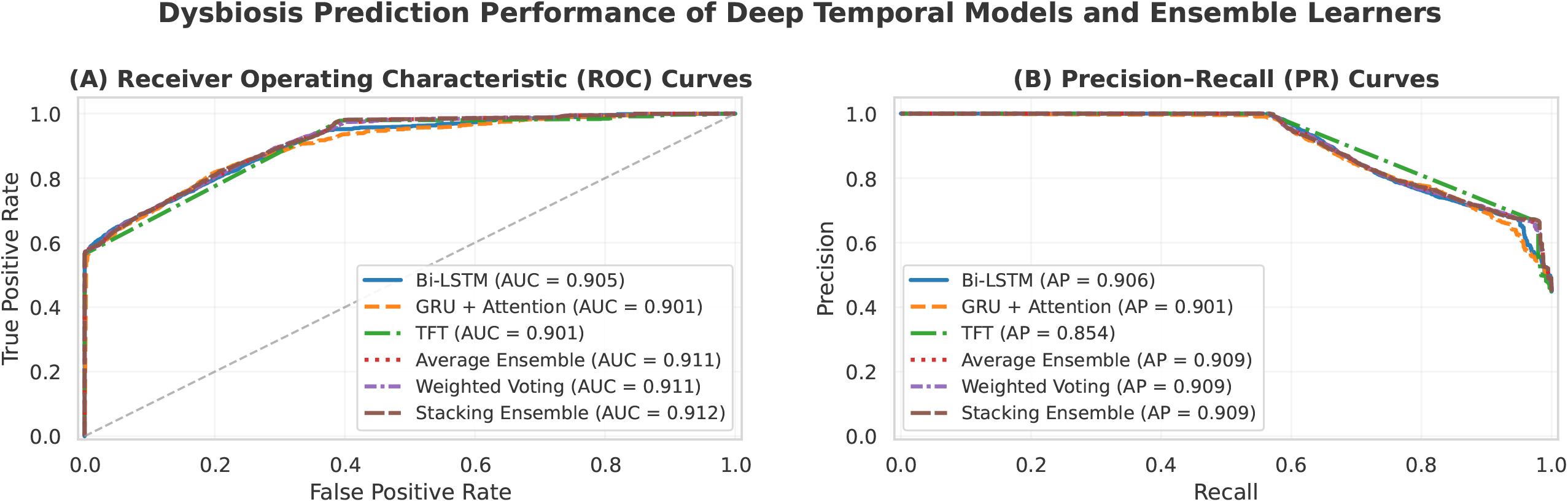

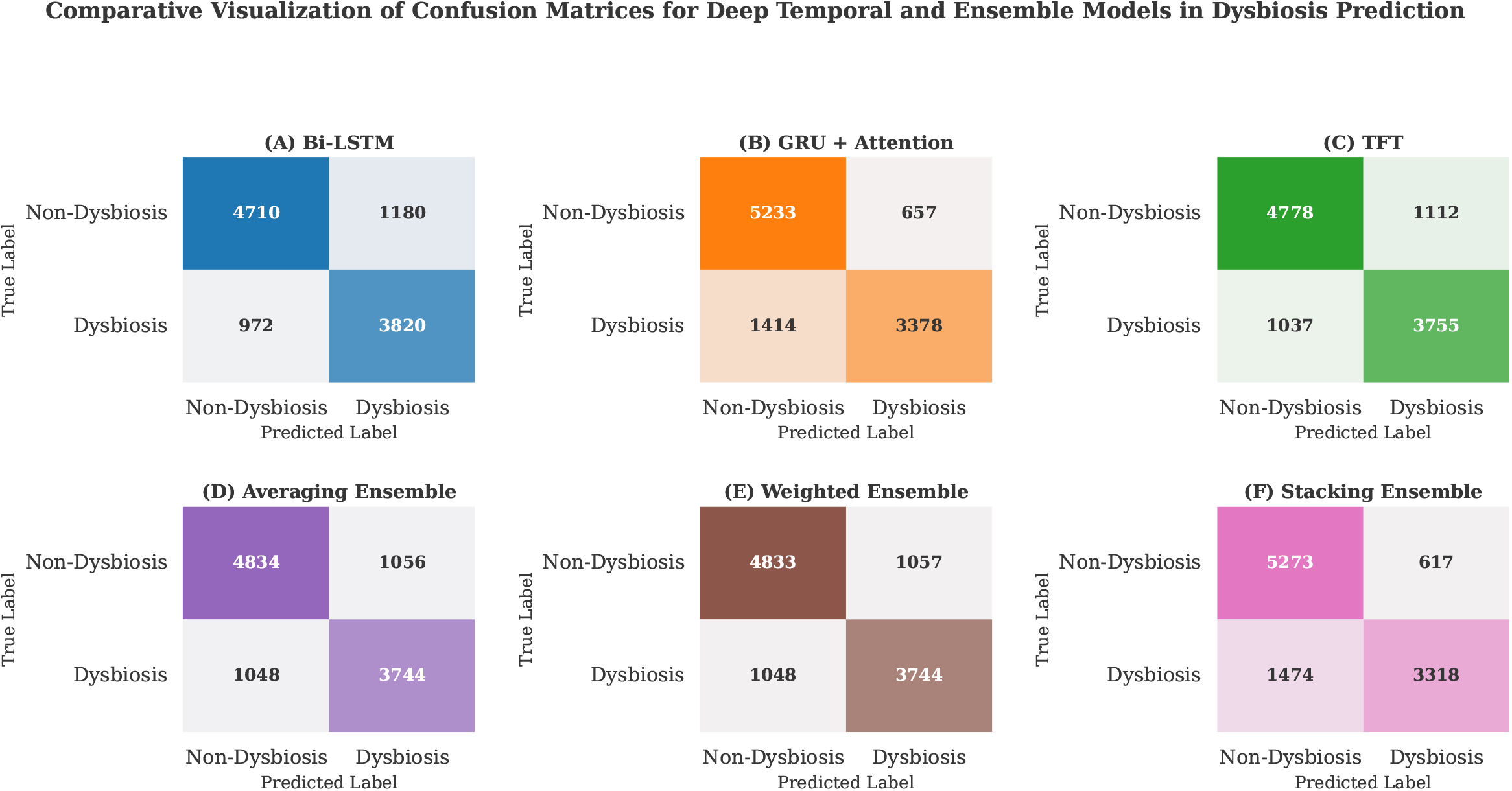

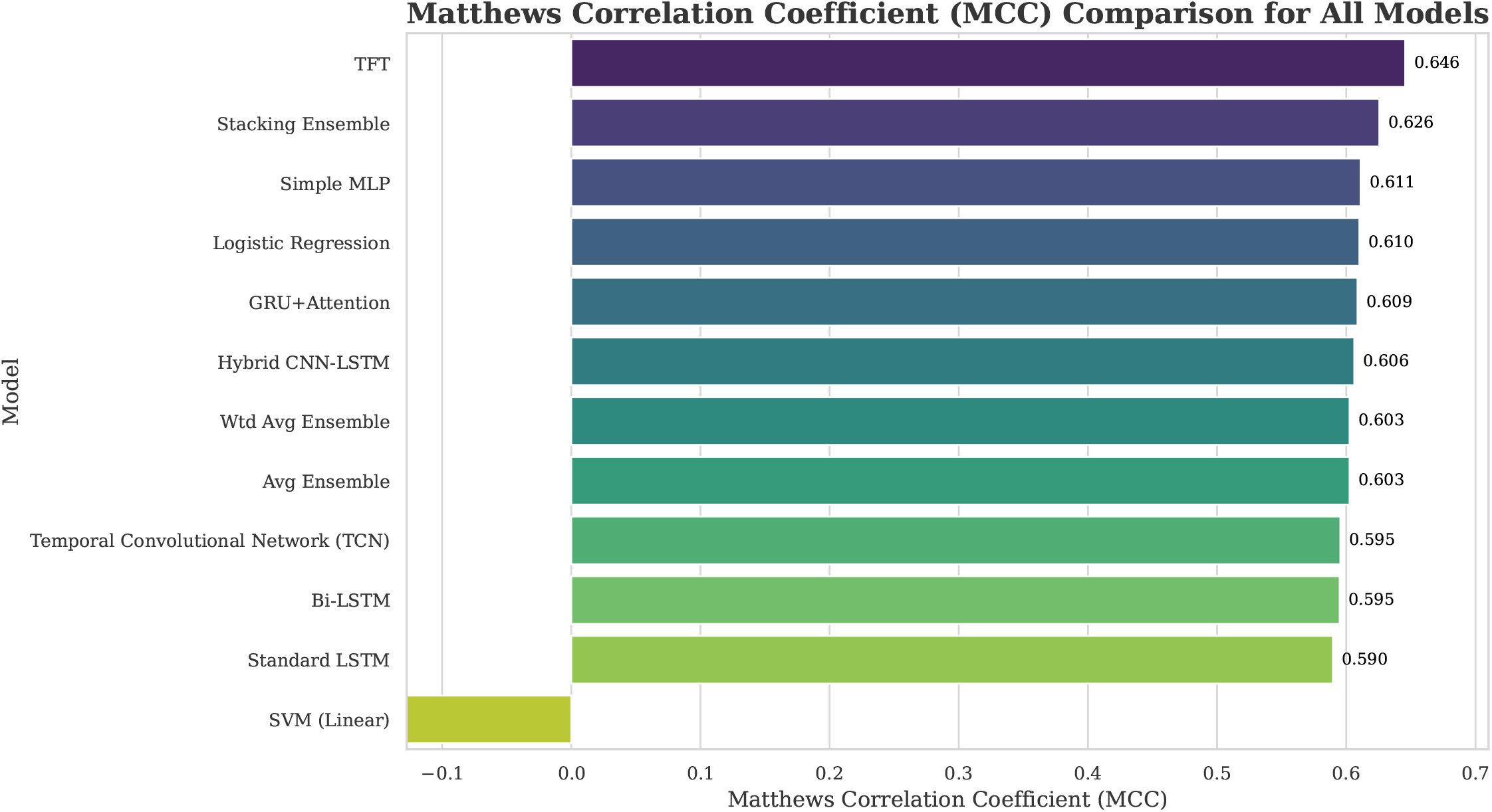

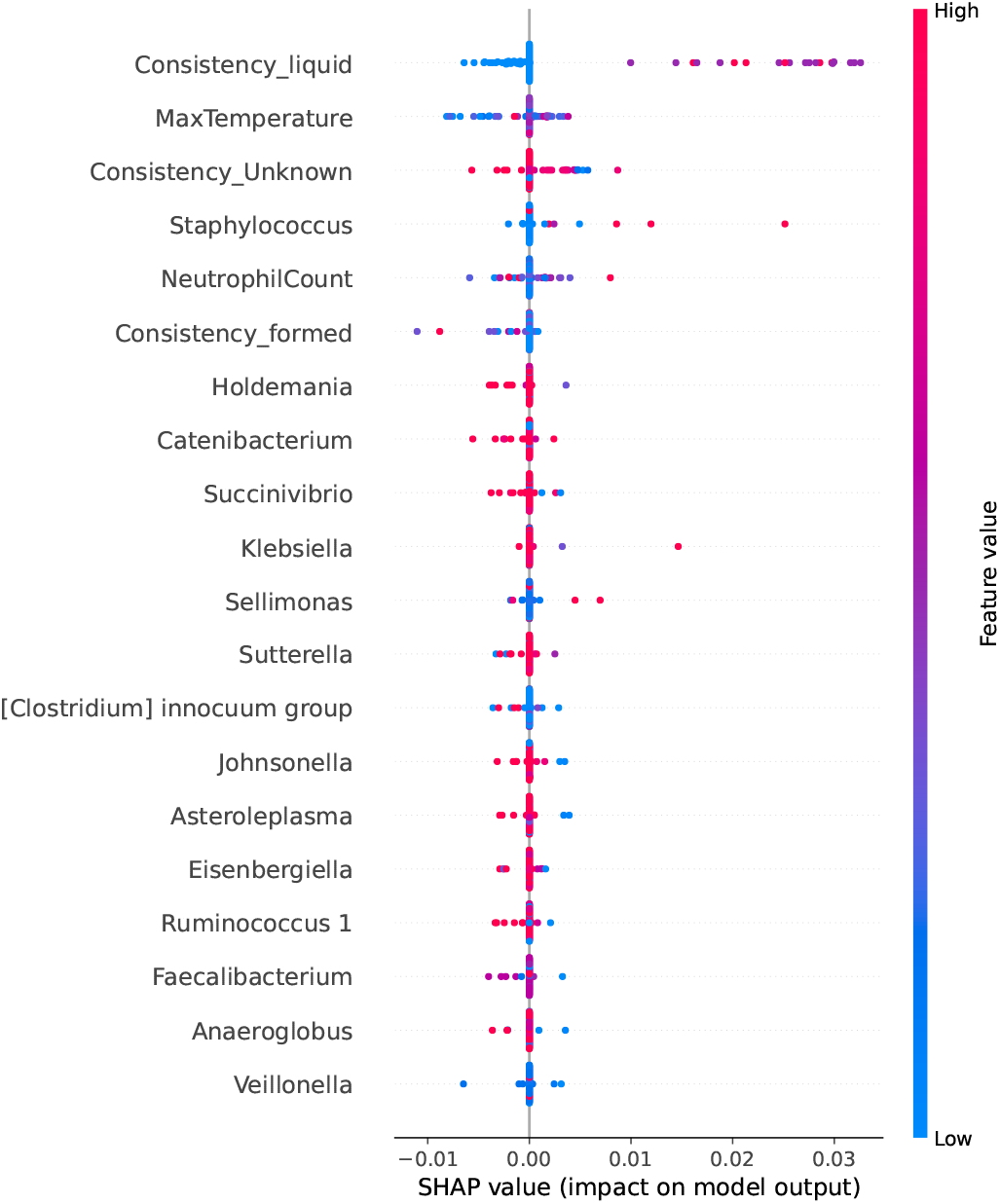

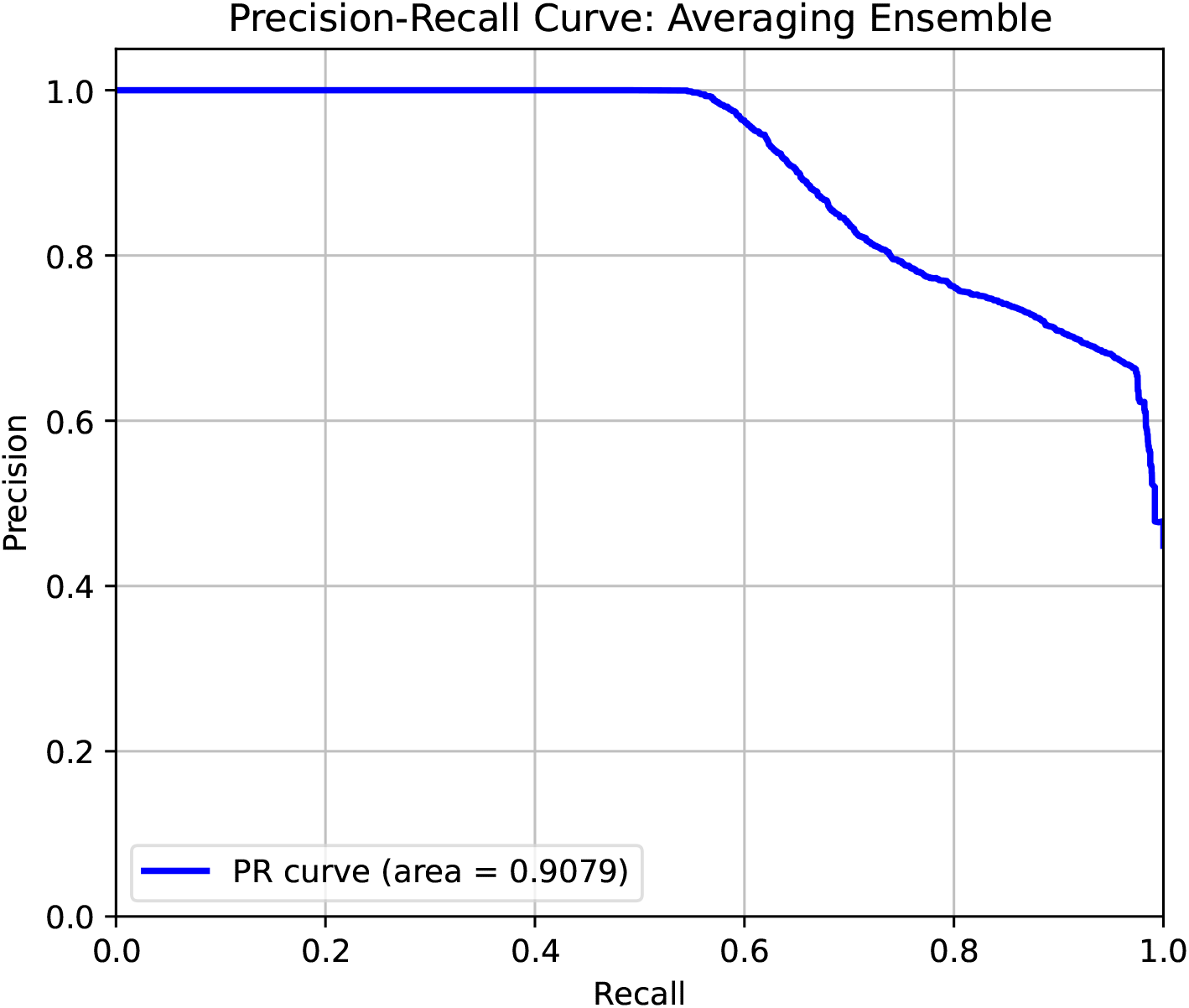

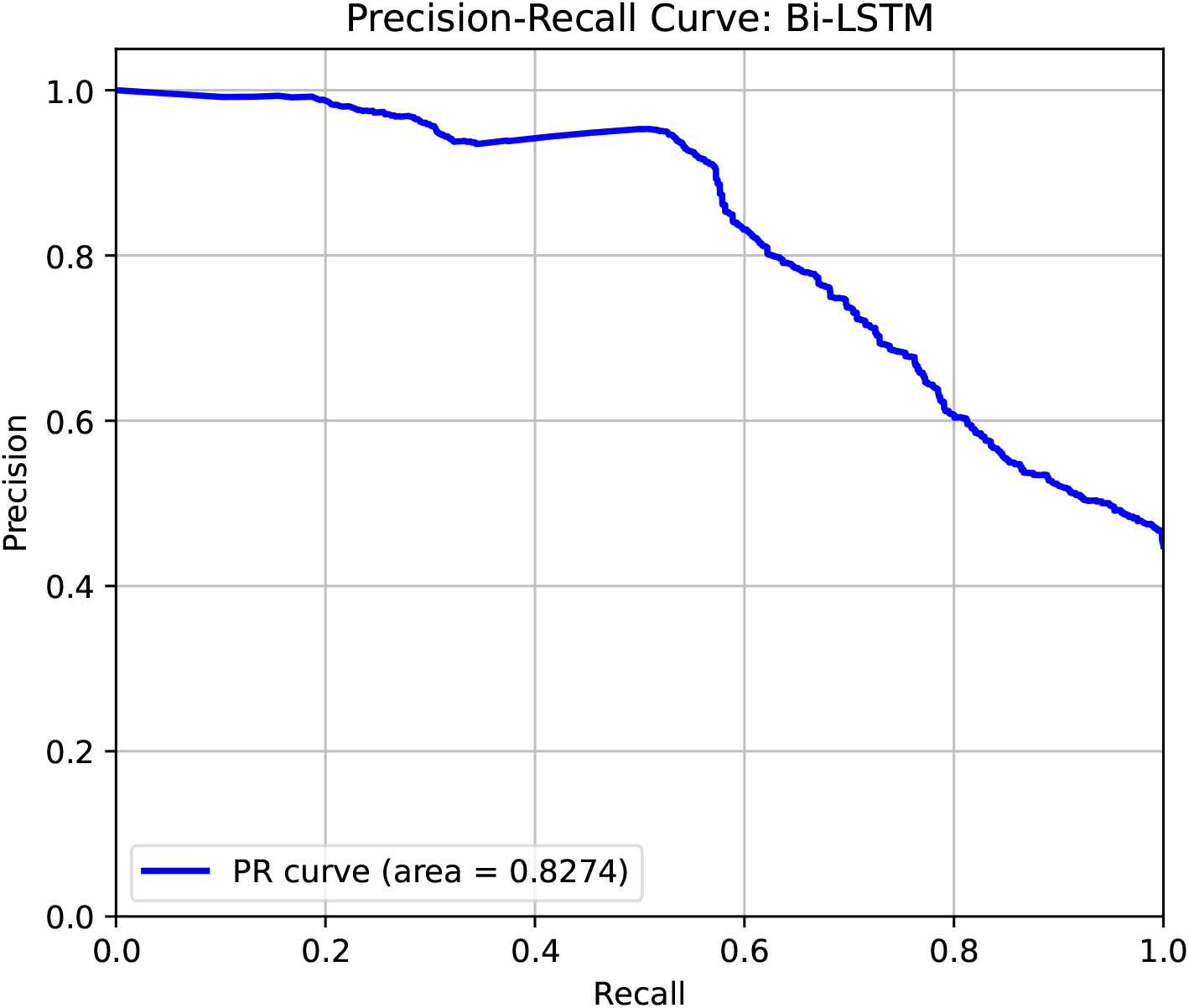

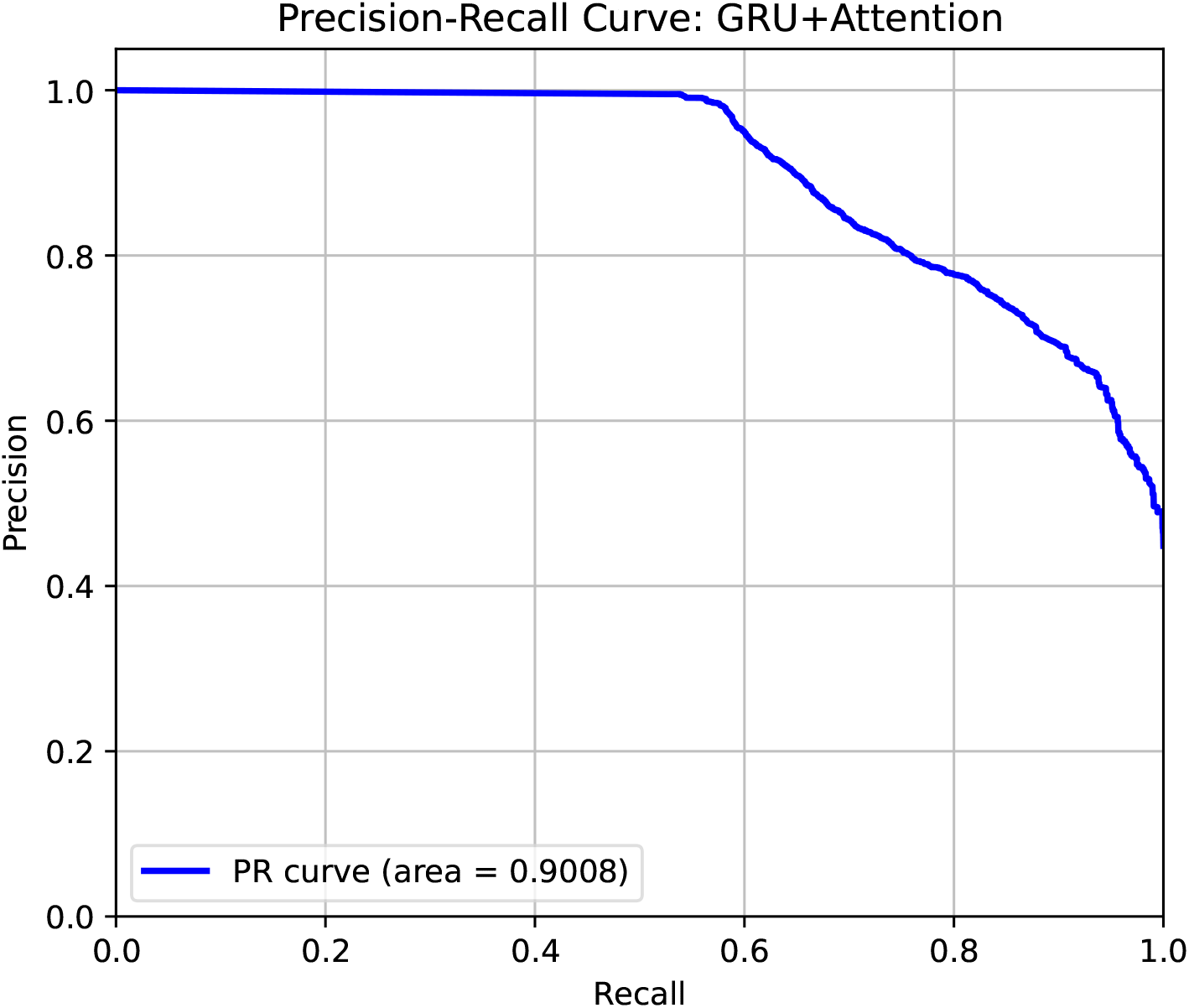

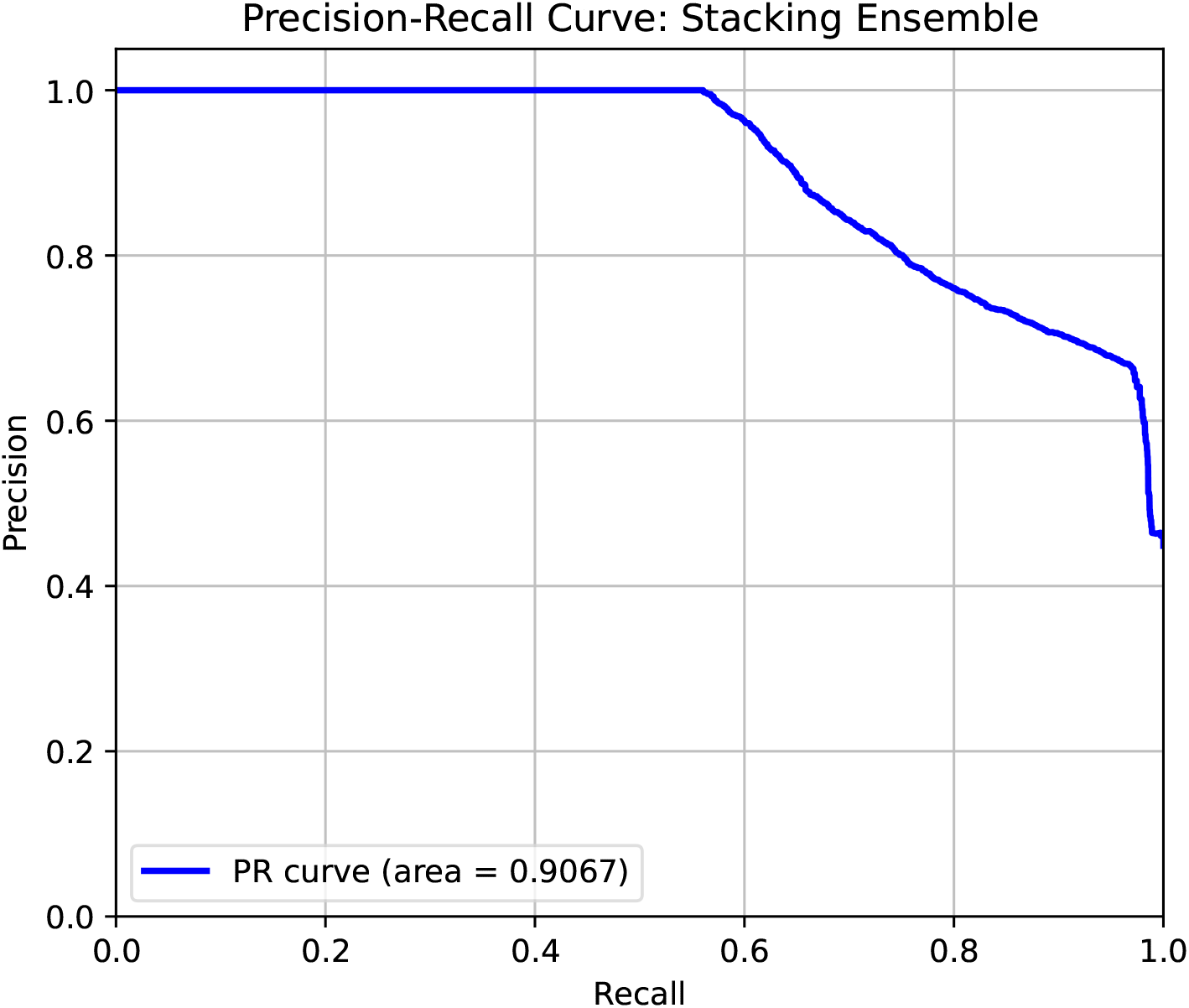

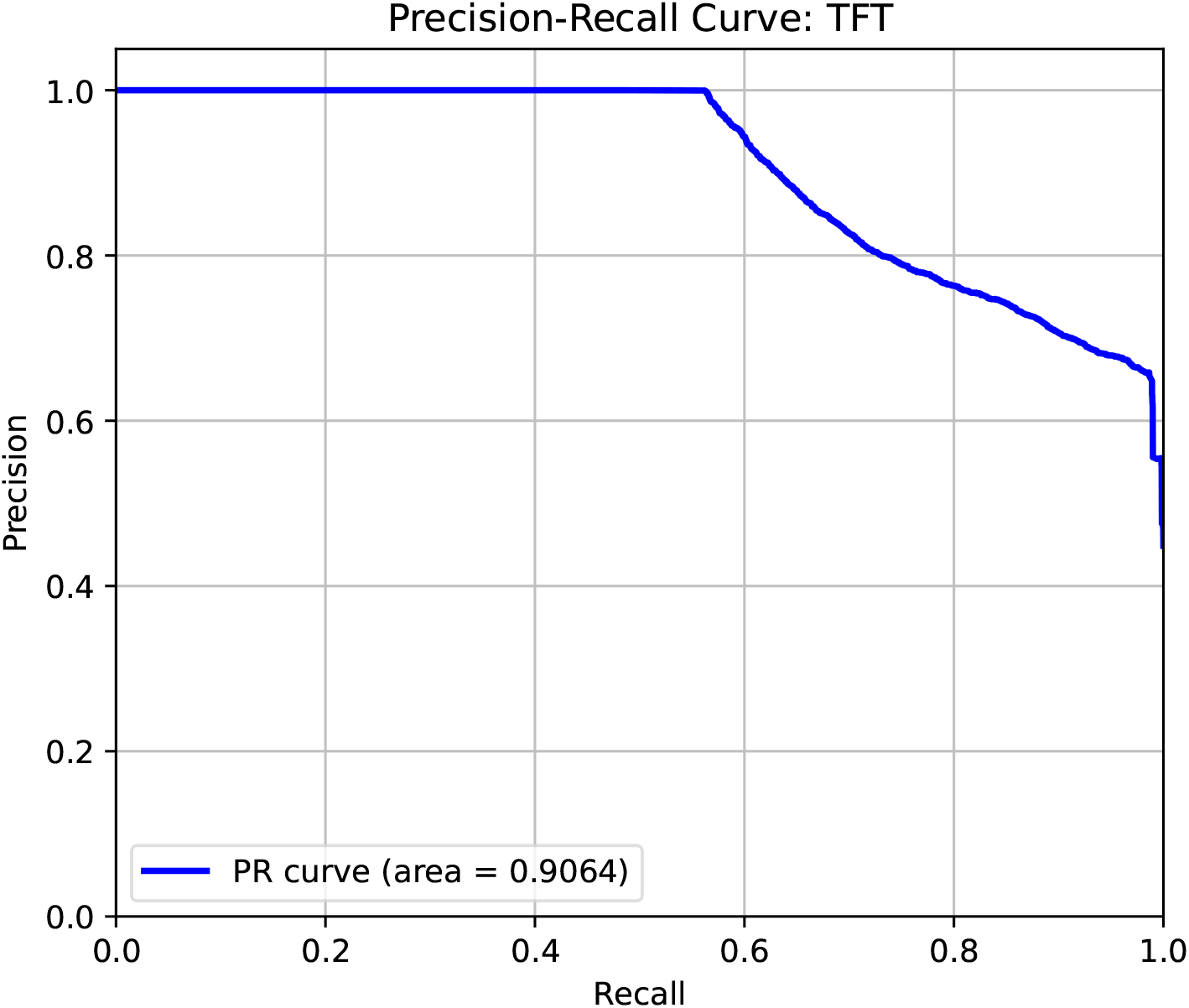

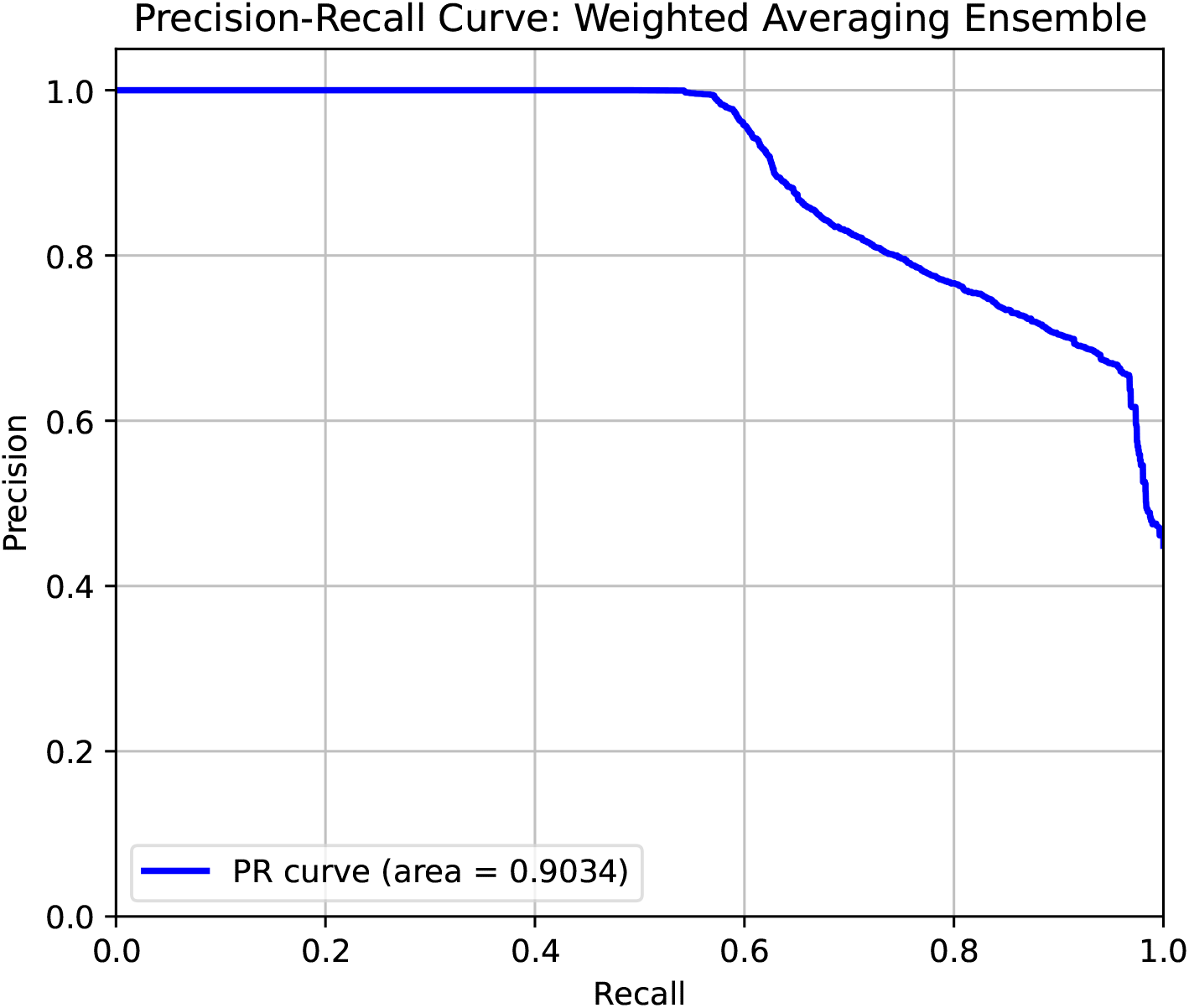

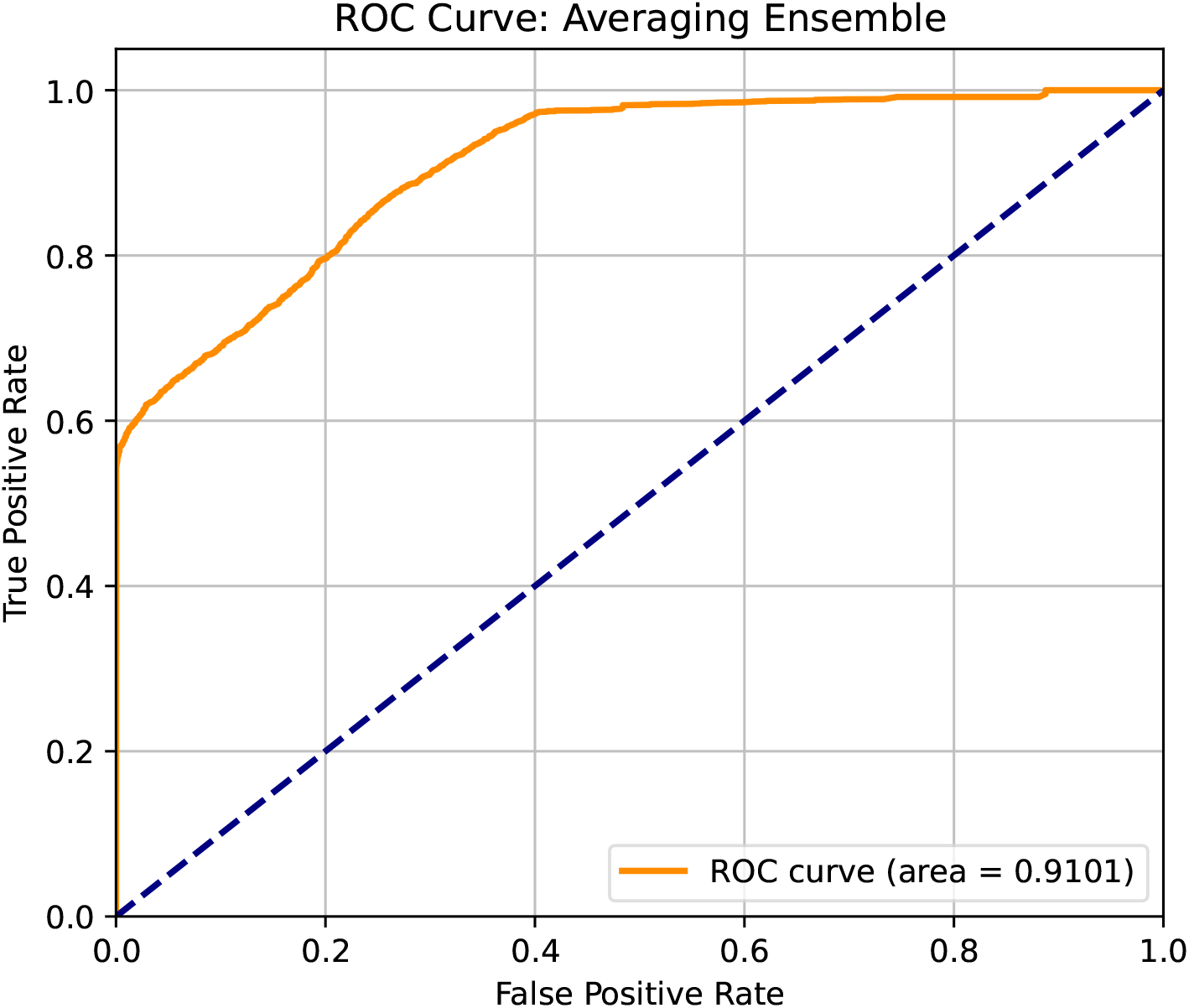

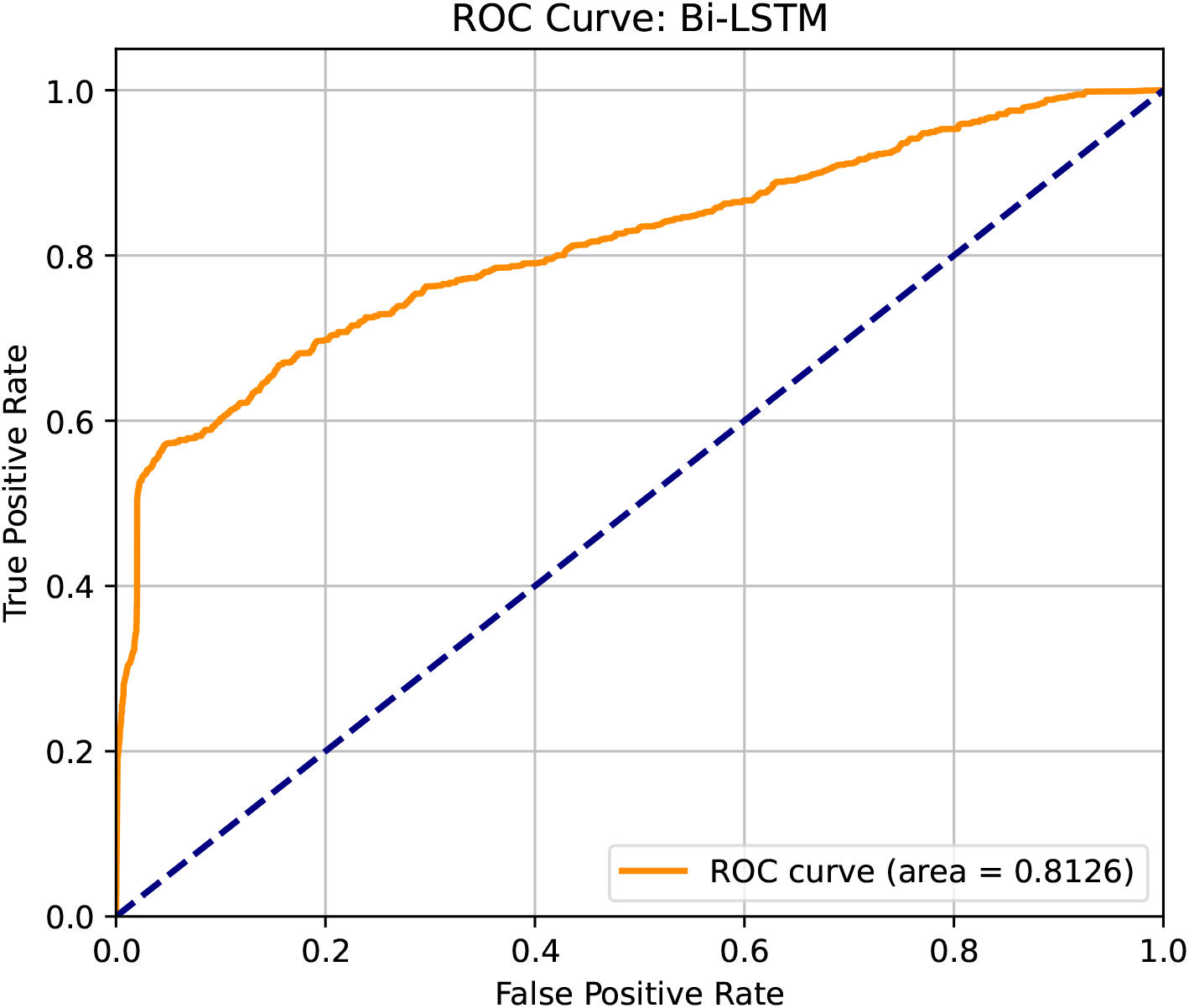

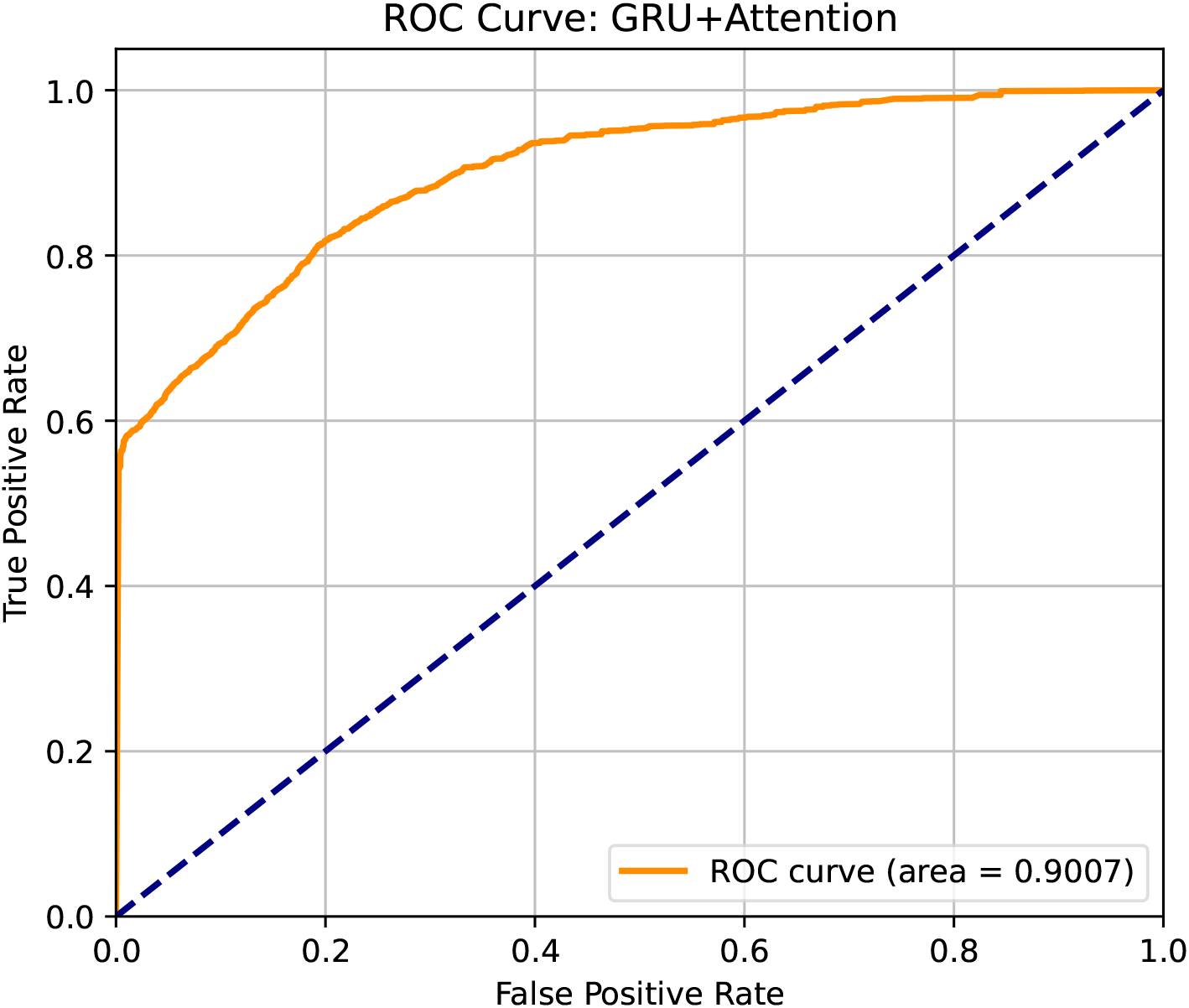

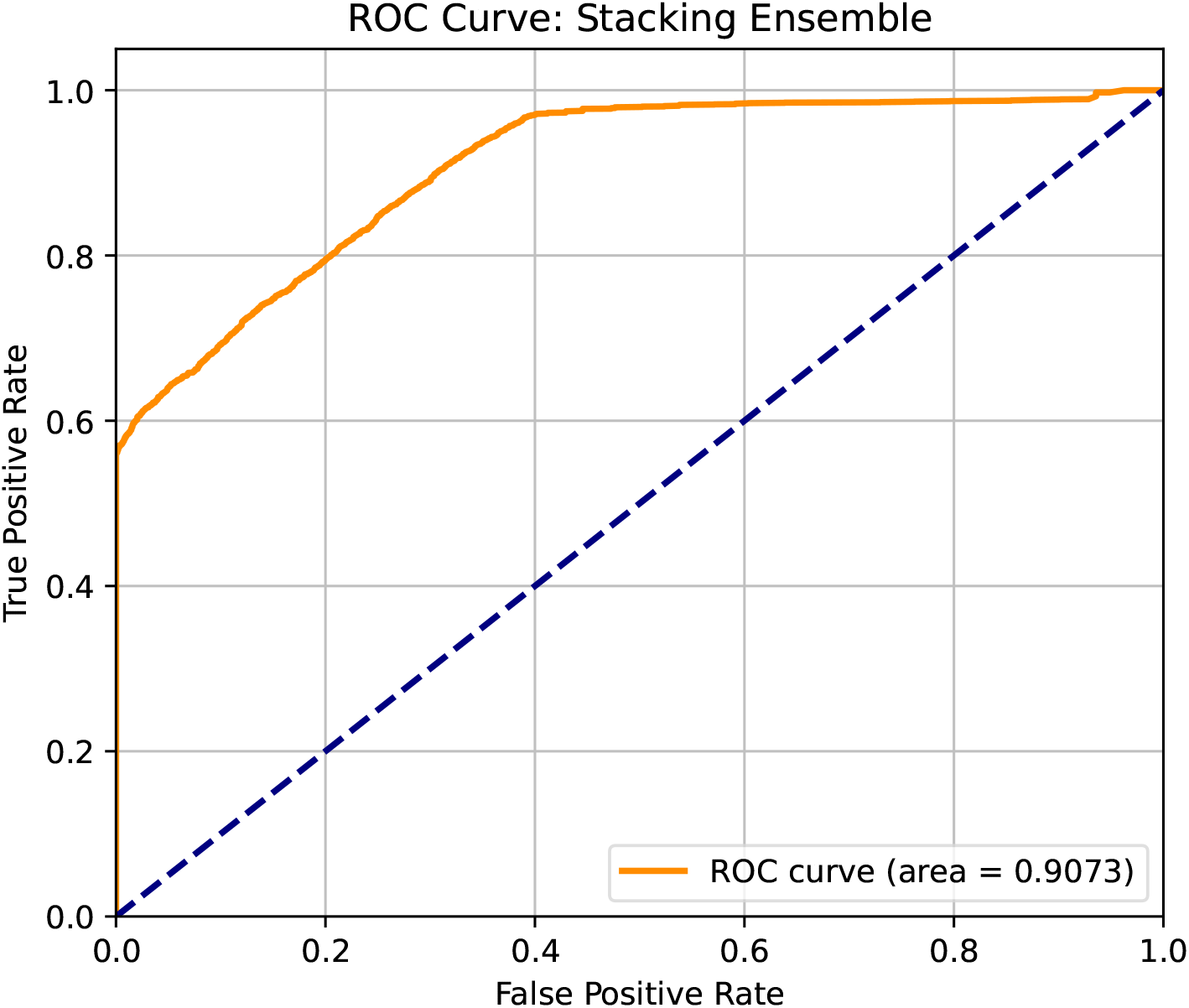

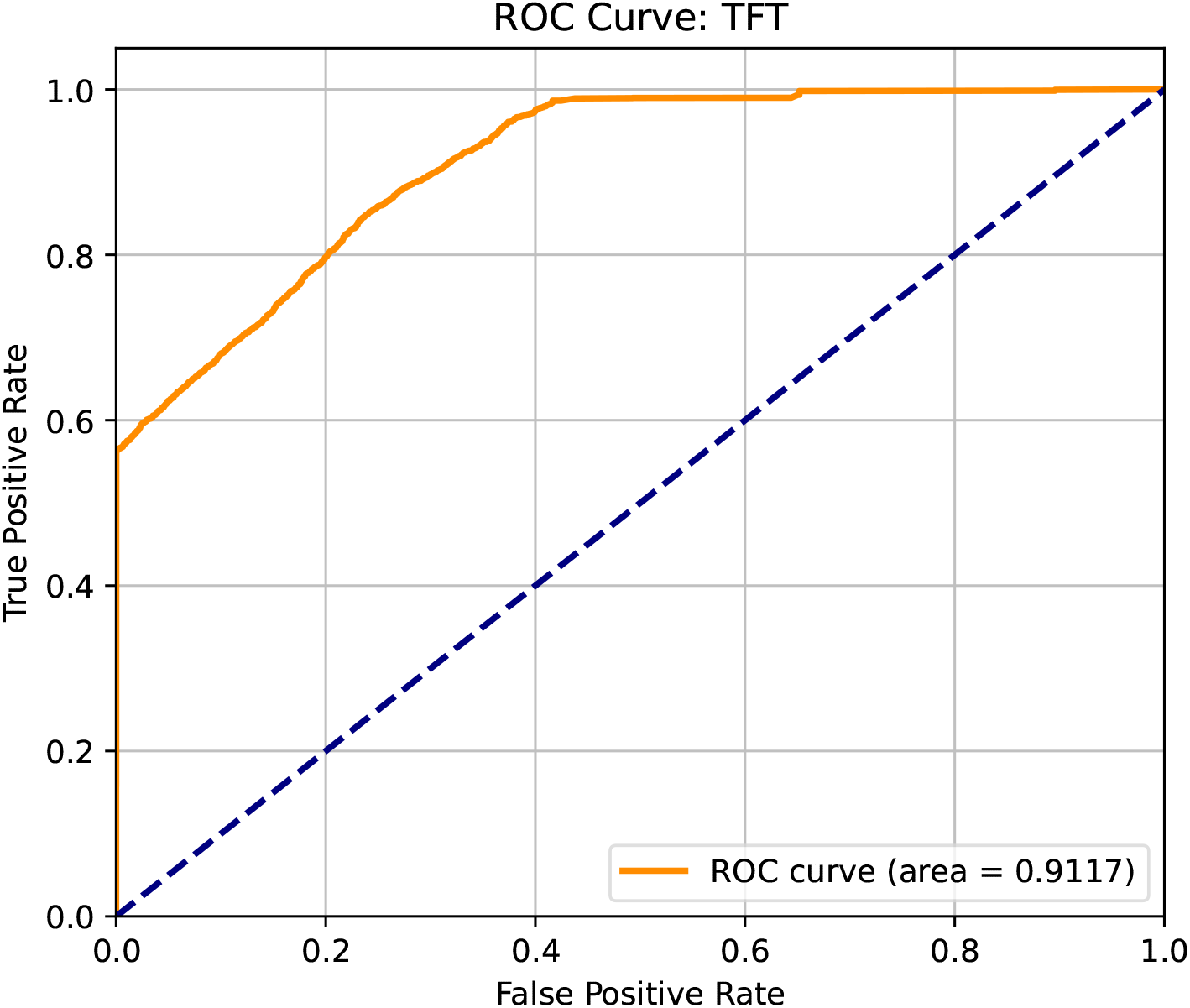

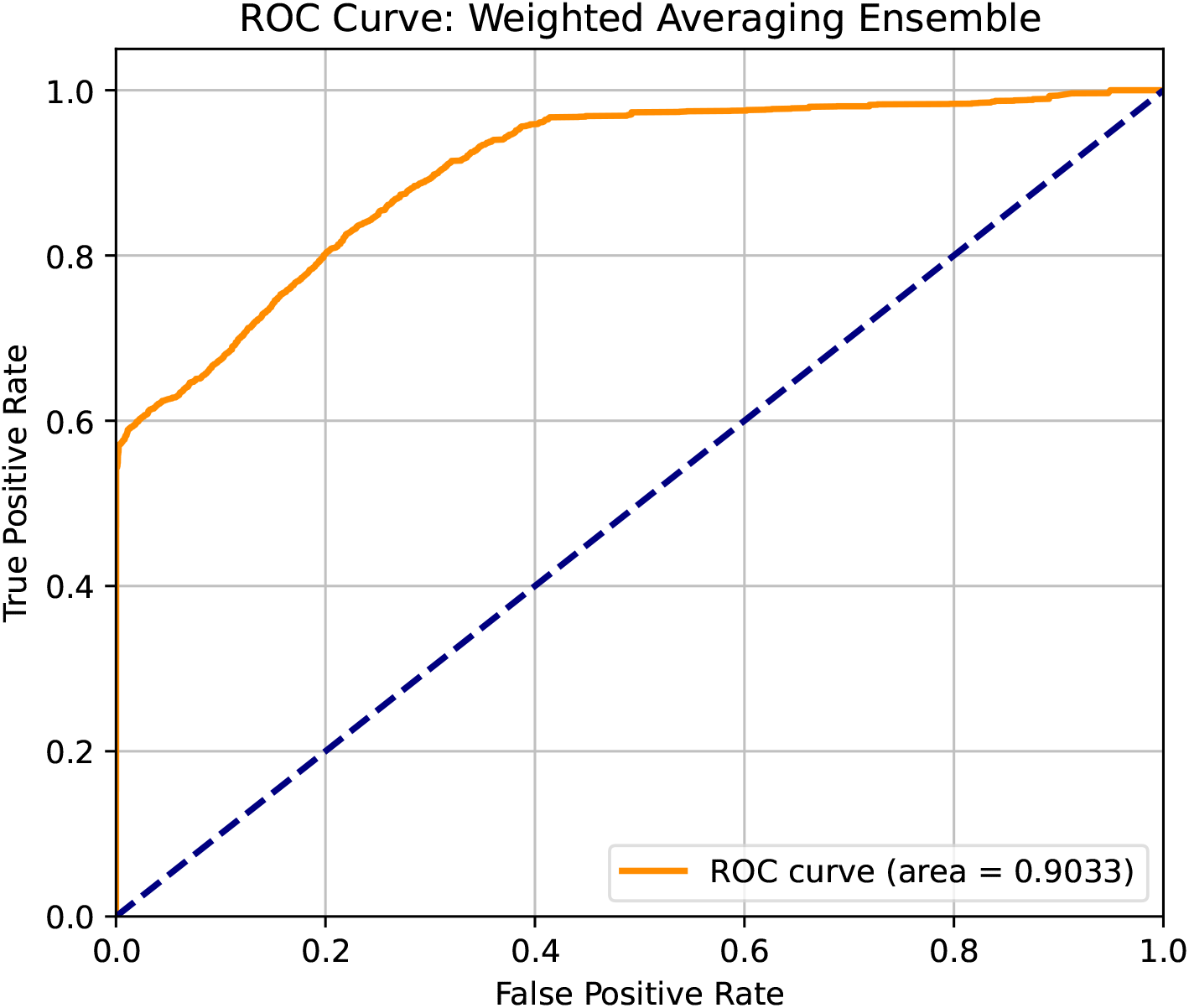

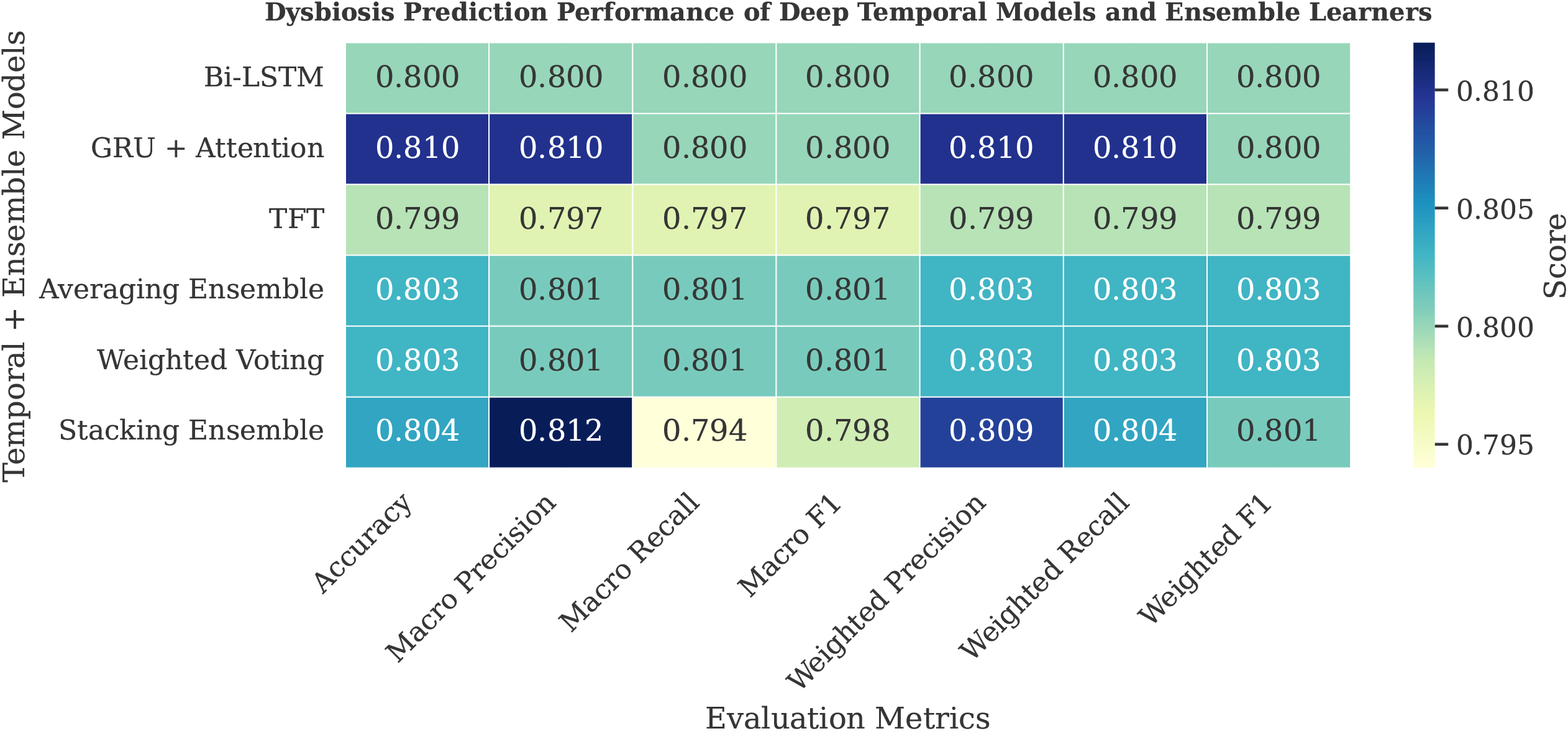

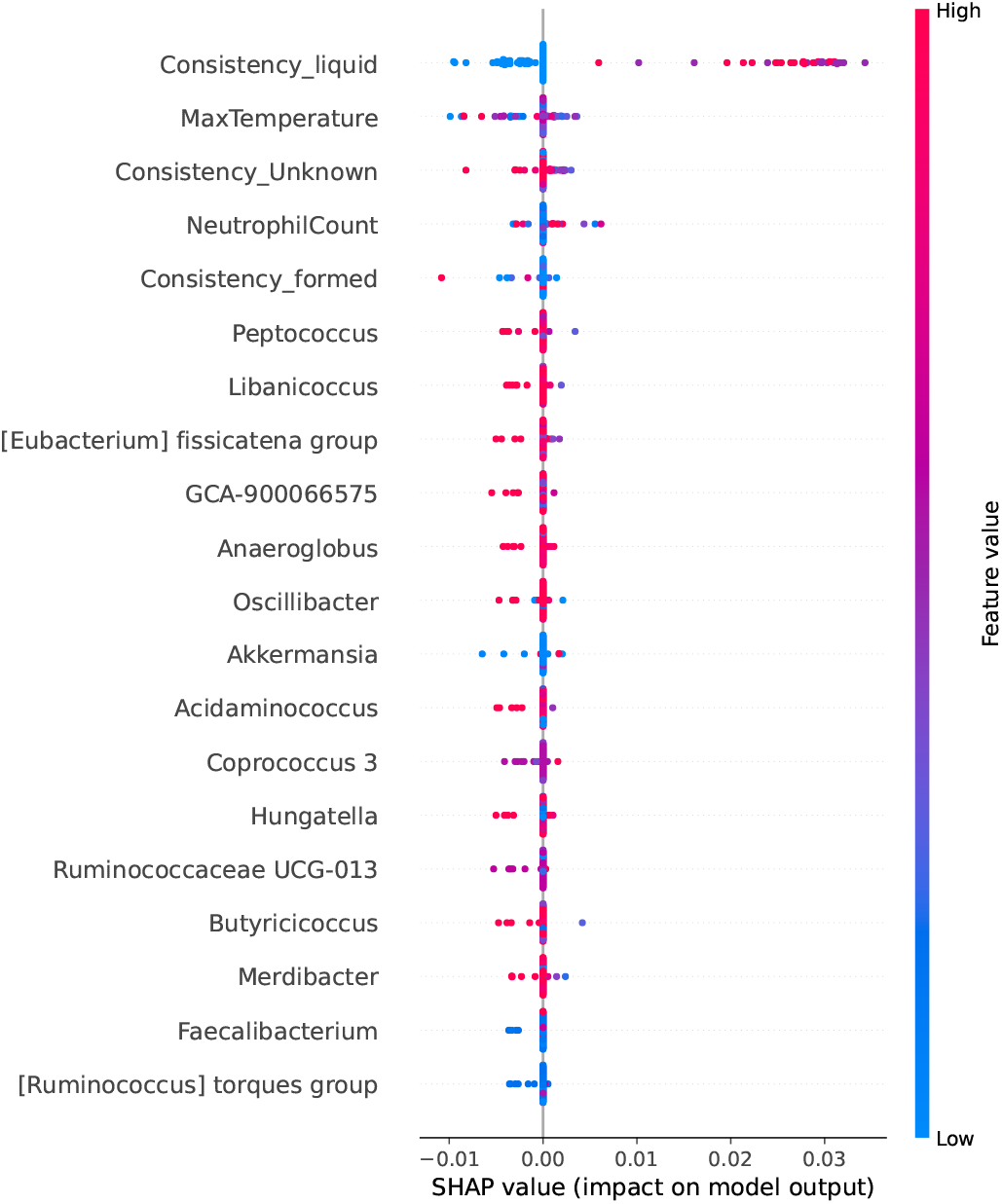

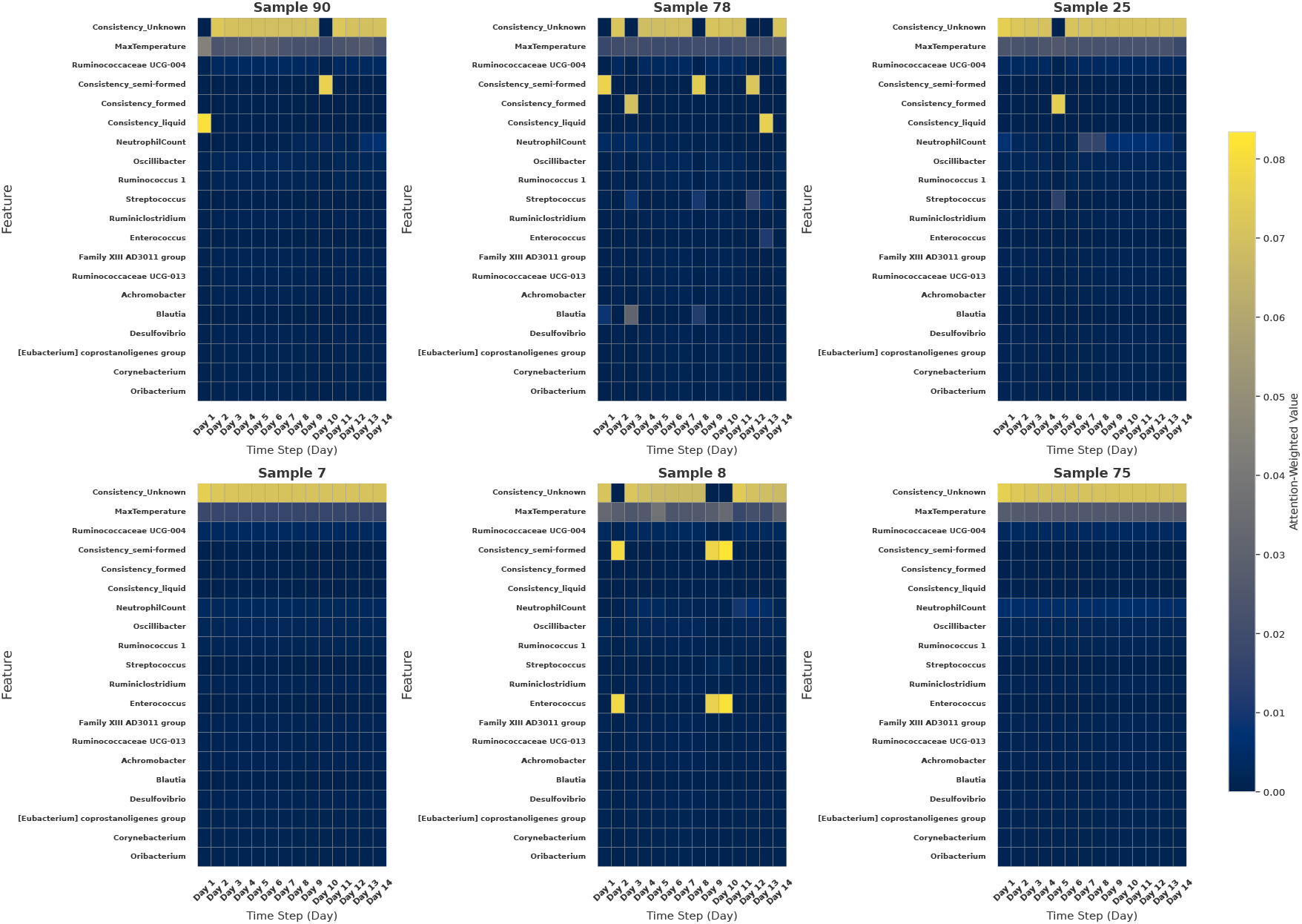

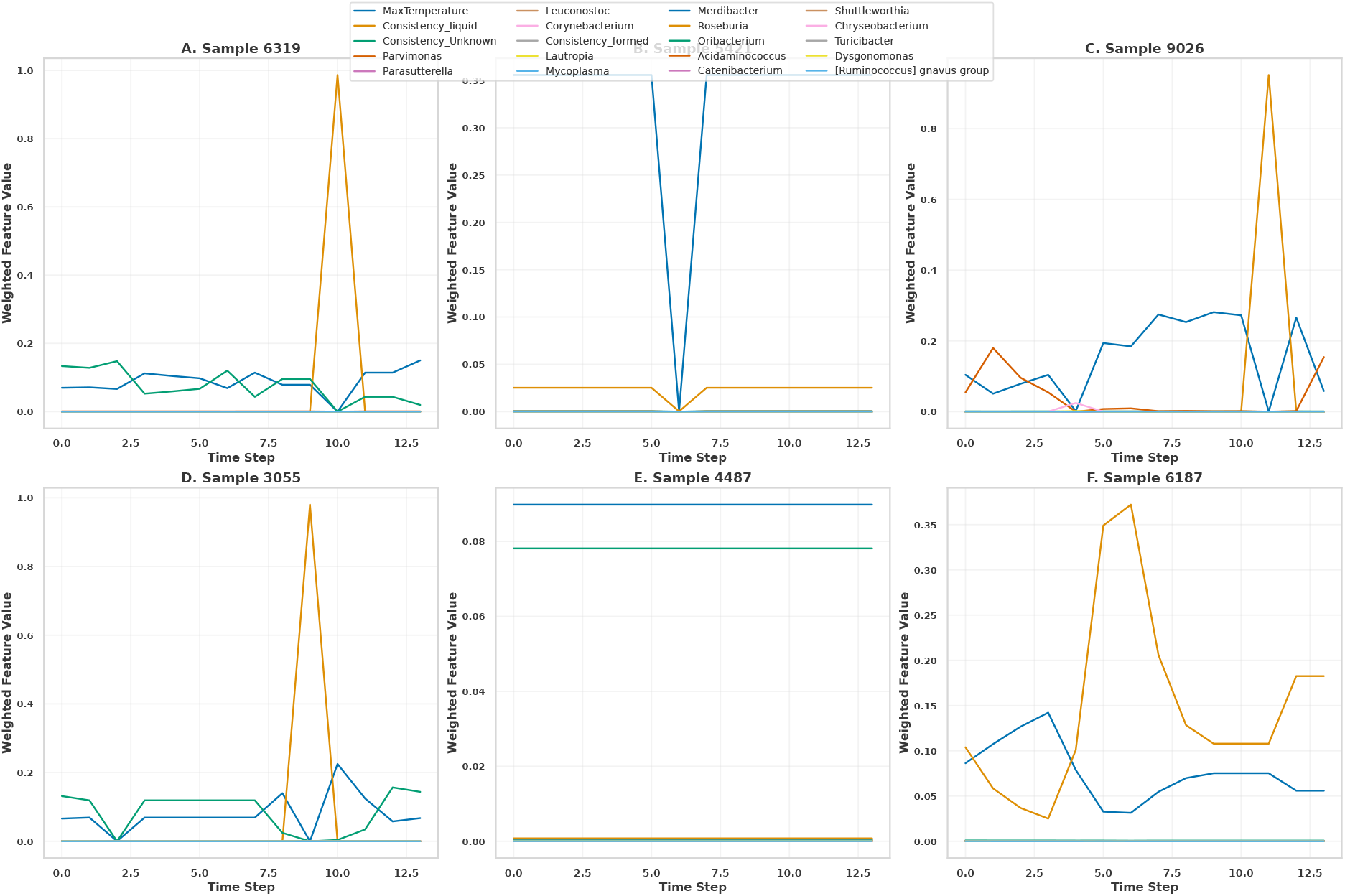

## Notes

### Competing Interest Statement

The authors have declared no competing interest.

### Summary of Updates

New Graphs have been added and also the Abstract and Discussion Sections are revised

