## Supplementory for "DynaBiomeX: An Interpretable Dual-Strategy Deep Learning Framework for Architectural Noise Filtration in Sparse Longitudinal Microbiome Data"

#### S1. Detailed Model Algorithms

##### *A. Preprocessing Pipeline*

**Algorithm S1** Microbiome Time-Series Preprocessing Pipeline

**Input:** Raw microbiome dataset with metadata
**Output:** Preprocessed sequences for time-series modeling

**Handle NeutrophilCount**
Define function `parse_neutrophil`:
**if** value == '$<$ 0.1' **then**
  Convert to 0.05
**else if** value is not a float **then**
  Convert to NaN
**end if**

**One-Hot Encode Consistency**
Apply one-hot encoding to 'Consistency' column

**Label Dysbiosis**
For each row, compute `RowLabel` based on temperature, neutrophil count, and consistency
Define function `patient_dysbiosis`:
**if** patient has $\geq$ 2 consecutive days with `RowLabel` = 1 **then**
  Assign `DysbiosisLabel` = 1
**else**
  Assign `DysbiosisLabel` = 0
**end if**
Apply function grouped by 'PatientID'

**Pivot Genus Abundances**
Reshape DataFrame: index = ('PatientID', 'SampleID', 'DayRelativeToNearestHCT'), columns = 'Genus', values = 'RelativeAbundance'
Fill missing genus entries with 0

**Merge Metadata and Apply CLR Transformation**
Merge pivoted genus abundances with selected metadata
Apply CLR transformation with pseudocount = 1 to genus columns

**Build Sequences and Split Data**
Define `build_sequences_with_labels`:
Create sliding windows per patient
Label = dysbiosis status at last timestep
Perform patient-level stratified split into train, validation, and test sets
Build sequences for each split

**Apply Variance Thresholding**
On training set genus features:
Remove features with low variance
Exclude same features from validation and test sets

**Scale Features**
Fit `MinMaxScaler` on training data
Transform training, validation, and test sets

**Return:** Preprocessed sequences ready for time-series modeling

##### *B. Bi-LSTM Architecture*

**Algorithm S2** Bi-LSTM Model Building and Training with Keras Tuner

**Input:** Preprocessed time-series data $(X_{\text{train}}, y_{\text{train}})$ and $(X_{\text{val}}, y_{\text{val}})$
**Output:** Trained Bi-LSTM model with optimized hyperparameters
**Procedure:** BuildBiLSTMModel($hp$)
$lstm\_units \leftarrow hp.\texttt{Int}(32, 256, step = 32)$
$dropout\_rate \leftarrow hp.\texttt{Float}(0.1, 0.5, step = 0.1)$
$learning\_rate \leftarrow hp.\texttt{Choice}([10^{-2}, 10^{-3}, 10^{-4}])$
Create Sequential model with input shape $(14, 139)$
Add Bidirectional LSTM layer with $lstm\_units$, return sequences
Add Dropout($dropout\_rate$), BatchNormalization
Add Bidirectional LSTM layer with $lstm\_units/2$, no return sequences
Add Dropout($dropout\_rate$), BatchNormalization
Add Dense layers with tunable units and ReLU activation
Add Dropout layers
Add final Dense(1) with sigmoid activation
Compile model with Adam($learning\_rate$), binary crossentropy, accuracy, and AUC
**return** model

**Hyperparameter Tuning Phase:**
Initialize Keras Tuner with RandomSearch on `BuildBiLSTMModel`
Set objective to maximize validation AUC
Set max trials = 10, executions per trial = 1
Apply EarlyStopping on val_AUC with patience = 3
Run tuner.search on training data for 10 epochs
Retrieve best hyperparameters $hp^*$

**Final Training Phase:**
Build final model using `BuildBiLSTMModel`($hp^*$)
Apply EarlyStopping and ModelCheckpoint on val_AUC
Train model on $(X_{\text{train}}, y_{\text{train}})$ with validation on $(X_{\text{val}}, y_{\text{val}})$
Set epochs = 50, batch size = 32
**return** Trained Bi-LSTM model

##### *C. GRU Architecture*

**Algorithm S3** Optimized GRU–Attention Model with Hyperparameter Tuning

**Require:** Training data $(X_{\text{train}}, y_{\text{train}})$, Validation data $(X_{\text{val}}, y_{\text{val}})$, Sequence length $L$, Feature dimension $F$, Maximum trials $T$, Epochs $E$
**Ensure:** Tuned GRU–Attention model $M^*$ and best hyperparameters $\theta^*$

**Step 1: Define Custom Attention Layer**
Initialize trainable weights $W \in \mathbb{R}^{F \times 1}$, bias $b \in \mathbb{R}^{L \times 1}$
**for** each timestep $t = 1$ to $L$ **do**
    $e_t \leftarrow \tanh(X_t W + b_t)$ {Compute attention score}
**end for**
$\alpha_t \leftarrow \dfrac{\exp(e_t)}{\sum_{k=1}^{L} \exp(e_k)}$ {Softmax normalization}
$c \leftarrow \sum_{t=1}^{L} \alpha_t X_t$ {Context vector}
**return** $c \in \mathbb{R}^F$

**Step 2: Define Hyperparameter Search Space**
Tune $\theta = \{$ GRU units $u \in [64, 256]$, Dropout rates $d_i \in [0.2, 0.5]$, Dense units $h_1 \in [32, 128]$, $h_2 \in [16, 64]$, Learning rate $\eta \in \{10^{-4}, 10^{-3}, 10^{-2}\}$ $\}$

**Step 3: Model Construction and Training (per trial $t$)**
**for** $t = 1$ to $T$ **do**
    Sample hyperparameters $\theta_t$ from search space
    Initialize model $M_t$:
        Input shape: $(L, F)$
        GRU($u$, `return_sequences=True`)
        Dropout($d_1$), BatchNormalization
        GRU($u$, `return_sequences=True`)
        Dropout($d_2$), BatchNormalization
        Apply Attention layer $\rightarrow c$
        Dense($h_1$, ReLU), Dropout($d_3$)
        Dense($h_2$, ReLU), Dropout($d_4$)
        Dense(1, Sigmoid)
    Compile $M_t$ with Adam($\eta$), loss = binary cross-entropy, metrics = {accuracy, AUC}
    Train $M_t$ on $(X_{\text{train}}, y_{\text{train}})$ with validation $(X_{\text{val}}, y_{\text{val}})$
    Apply EarlyStopping on `val_AUC` (patience = 3)
    Record best validation AUC as $\text{AUC}_t$
**end for**

**Step 4: Model Selection and Final Training**
$\theta^* \leftarrow \arg\max_{\theta_t} \text{AUC}_t$
Build final model $M^*$ using $\theta^*$
Train $M^*$ on full training set for $E$ epochs with EarlyStopping and checkpointing
Save $M^*$ and training history

**Output:** Best model $M^*$, hyperparameters $\theta^*$, and attention weights $\{\alpha_t\}$ for interpretability

##### *D. TFT Architecture*

**Algorithm S4** Adapted Temporal Fusion Transformer (TFT) with Noise Filtering

**Require:** Time-series microbiome matrix $\mathbf{X} \in \mathbb{R}^{T \times F}$; static covariates $\mathbf{s}$
**Ensure:** Predicted dysbiosis probability $\hat{y} \in [0, 1]$

**Step 1: Component Initialization**
Define Gated Linear Unit: $\text{GLU}(\gamma) = \sigma(\mathbf{W}_1\gamma + \mathbf{b}_1) \odot (\mathbf{W}_2\gamma + \mathbf{b}_2)$
Define Gated Residual Network (GRN) with ELU activation:
  $\eta_1 = \text{ELU}(\mathbf{W}_{\text{in}}\mathbf{x} + \mathbf{b}_{\text{in}})$ *// Matches code dense1*
  $\eta_2 = \mathbf{W}_{\text{out}}\eta_1 + \mathbf{b}_{\text{out}}$ *// Matches code dense2*
  $\text{GRN}(\mathbf{x}) = \text{LayerNorm}(\mathbf{x} + \text{GLU}(\eta_2))$

**Step 2: Variable Selection Network (VSN)**
*// Weighs importance of specific microbial features*
Compute feature weights: $\mathbf{v}_t = \text{Softmax}(\mathbf{W}_v \cdot \text{GRN}(\mathbf{X}_t))$
Apply weights: $\tilde{\mathbf{X}}_t = \mathbf{X}_t \odot \mathbf{v}_t$
Project to hidden dim: $\mathbf{E}_t = \text{Dense}(\tilde{\mathbf{X}}_t)$

**Step 3: Local Temporal Processing (Bi-LSTM/GRU)**
$\mathbf{H}_t = \text{GRU}(\mathbf{E}_t, \mathbf{H}_{t-1})$
Apply Gate & Residual: $\tilde{\mathbf{H}}_t = \text{LayerNorm}(\mathbf{H}_t + \text{GLU}(\mathbf{H}_t))$

**Step 4: Temporal Self-Attention (Global Context)**
Compute Attention: $\mathbf{A}_t = \text{Softmax}\left(\frac{\mathbf{QK}^\top}{\sqrt{d_k}}\right)\mathbf{V}$
Apply Gate & Residual: $\mathbf{C}_t = \text{LayerNorm}(\tilde{\mathbf{H}}_t + \text{GLU}(\mathbf{A}_t))$

**Step 5: Non-Linear "Sentinel" Filtering**
Apply final GRN to context vector: $\mathbf{O}_t = \text{GRN}(\mathbf{C}_t)$

**Step 6: Classification Head Adaptation**
Global Average Pooling: $\mathbf{z} = \frac{1}{T}\sum_{t=1}^{T}\mathbf{O}_t$ *// Matches GlobalAveragePooling1D*
Final Prediction: $\hat{y} = \sigma(\mathbf{W}_{\text{final}} \cdot \text{ReLU}(\mathbf{z}) + \mathbf{b}_{\text{final}})$

**Step 7: Optimization**
Minimize Binary Cross-Entropy Loss $\mathcal{L}$ with Adam optimizer
**return** Dysbiosis probability $\hat{y}$

#### S2. Statistical Validation

TABLE S1
Pairwise Statistical Comparison of Top-Performing Models (Bootstrapped Paired t-test, n=2000)

| Model A | Model B | Metric | Mean Diff | p-value | Verdict |
|---|---|---|---|---|---|
| Stacking Ensemble | Adapted TFT | ROC-AUC | +0.0109 | $< 0.001$ | Ensemble Superior |
| Stacking Ensemble | Bi-LSTM | ROC-AUC | +0.0069 | $< 0.001$ | Ensemble Superior |
| Adapted TFT | Stacking Ensemble | MCC | +0.0200 | $< 0.001$ | TFT Superior |
| Adapted TFT | Bi-LSTM | MCC | +0.0507 | $< 0.001$ | TFT Superior |
| Bi-LSTM | Adapted TFT | F1 (Dysbiosis) | +0.0585 | $< 0.001$ | Bi-LSTM Superior |

### S3. EXTENDED VISUALIZATION: FEATURE HEATMAPS

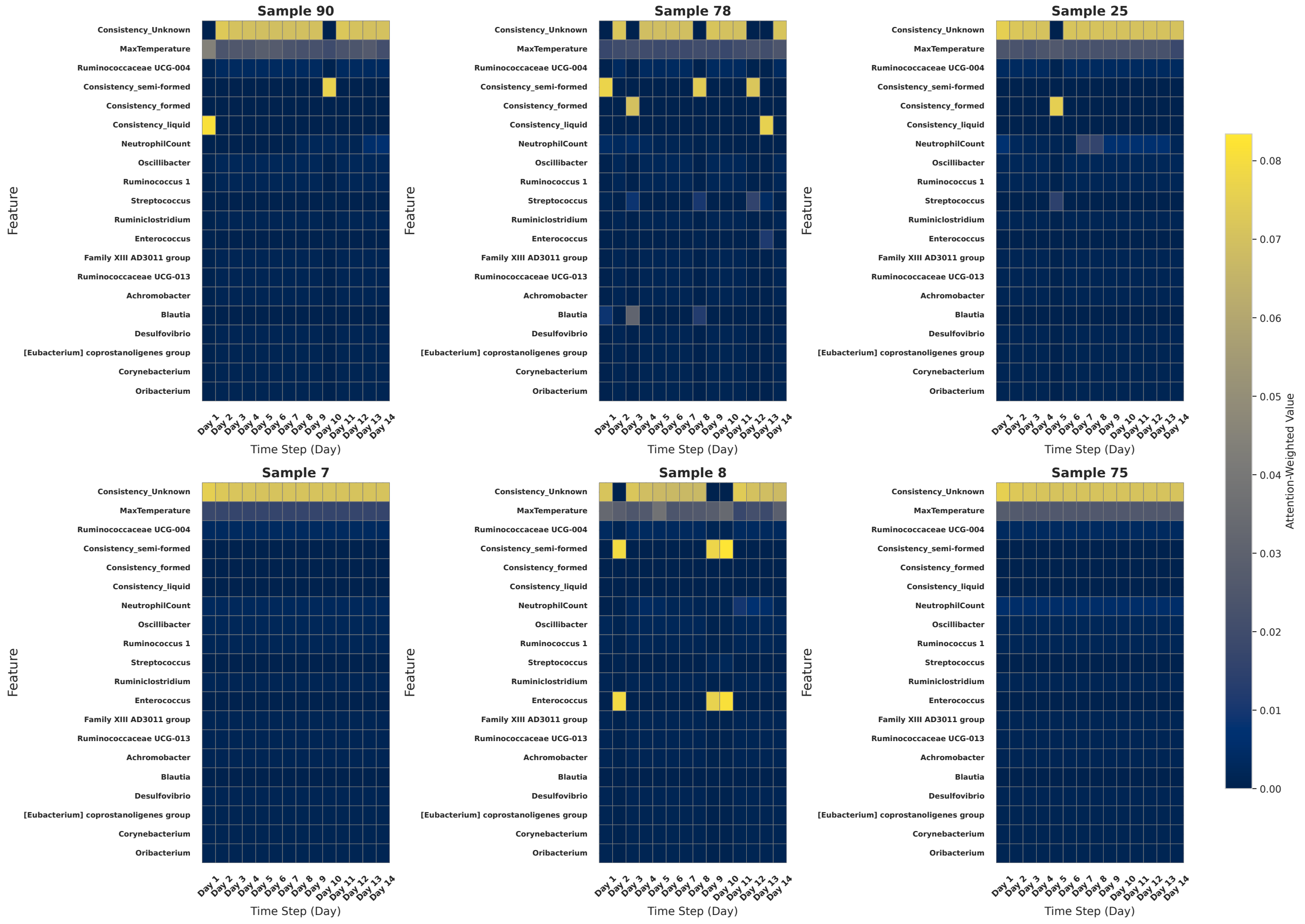


Fig. S1. Feature-level attention visualizations from the GRU + Attention model across six representative samples. Each heatmap displays attention weights assigned to clinical and microbial features over a 14-day sequence. Notably, features such as liquid stool consistency, neutrophil count, and elevated temperature consistently receive higher attention in dysbiosis sequences, while microbial genera like *Enterococcus* and *Streptococcus* emerge as key contributors. These patterns highlight the model's ability to temporally and biologically localize predictive signals.

#### S4. Extended Visualization: VSN Weights

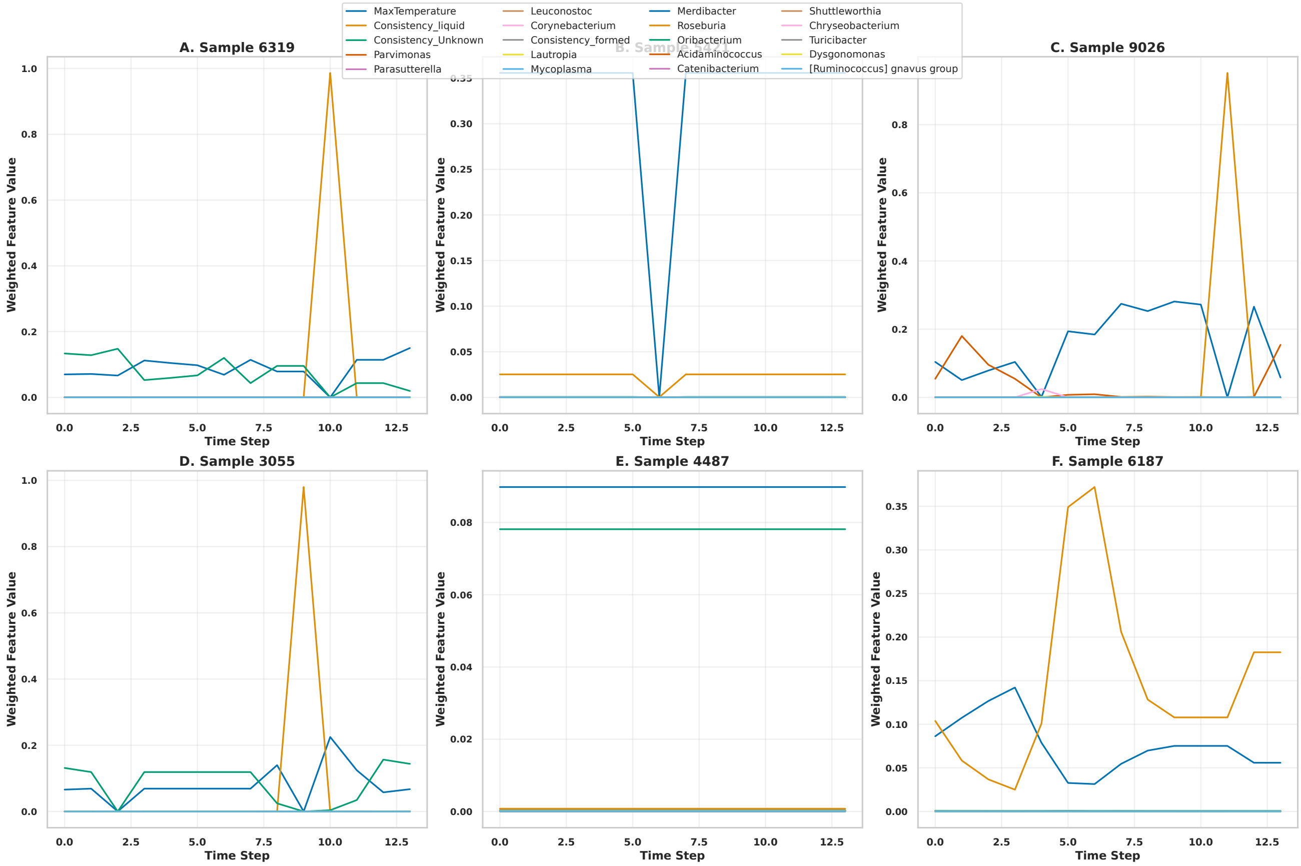


Fig. S2. Temporal Variable Selection Network (VSN) weights reveal patient-specific feature importance in detecting gut dysbiosis for ALLO-HCT patients. VSN weights from the Temporal Fusion Transformer show the top 20 features across six patient samples over 12-13 time steps. Samples 6319 (A) and 3055 (D) show acute dysbiosis with single dominant features. Samples 5421 (B) and 6187 (F) display moderate, distributed patterns. Sample 9026 (C) shows intermediate dynamics. Sample 4487 (E) exhibits stable, low-weight patterns indicating minimal dysbiosis risk.

#### S5. Detailed Tables

TABLE S2
Summary of key pairwise model comparisons across evaluation metrics using bootstrapped paired t-tests. All reported differences are statistically significant at $\alpha = 0.05$.

| **Model 1** | **Model 2** | **Metric** | **Mean Diff** | **$t$-statistic** | **$p$-value** | **Model 1 > Model 2** |
|---|---|---|---|---|---|---|
| Bi-LSTM | GRU+Attention | ROC-AUC | −0.0493 | −483.29 | < 0.001 | False |
| Bi-LSTM | GRU+Attention | PR-AUC | −0.0241 | −311.07 | < 0.001 | False |
| Bi-LSTM | GRU+Attention | MCC | −0.2960 | −928.12 | < 0.001 | False |
| Bi-LSTM | GRU+Attention | F1 Score | −0.0942 | −525.22 | < 0.001 | False |
| Bi-LSTM | TFT | ROC-AUC | −0.0603 | −576.53 | < 0.001 | False |
| Bi-LSTM | TFT | PR-AUC | −0.0297 | −381.80 | < 0.001 | False |
| Bi-LSTM | TFT | MCC | −0.2814 | −904.31 | < 0.001 | False |
| Bi-LSTM | TFT | F1 Score | −0.1064 | −664.17 | < 0.001 | False |
| GRU+Attention | TFT | ROC-AUC | −0.0111 | −181.29 | < 0.001 | False |
| GRU+Attention | TFT | PR-AUC | −0.0056 | −93.48 | < 0.001 | False |
| GRU+Attention | TFT | MCC | 0.0146 | 61.74 | < 0.001 | True |
| GRU+Attention | TFT | F1 Score | −0.0123 | 88.58 | < 0.001 | False |
| GRU+Attention | Stacking Ensemble | ROC-AUC | −0.0099 | −231.97 | < 0.001 | False |
| GRU+Attention | Stacking Ensemble | MCC | −0.0146 | −74.79 | < 0.001 | False |
| TFT | Stacking Ensemble | ROC-AUC | 0.0012 | 32.75 | < 0.001 | True |
| TFT | Stacking Ensemble | MCC | −0.0292 | −143.79 | < 0.001 | False |
| Averaging Ensemble | Stacking Ensemble | ROC-AUC | −0.0013 | −37.79 | < 0.001 | False |
| Averaging Ensemble | Stacking Ensemble | MCC | −0.0428 | −176.14 | < 0.001 | False |

#### S6. Baseline Model Implementation Details

To ensure a rigorous evaluation, all benchmark models were optimized using the same validation protocol as the DynaBiomeX framework.

##### A. Static Baselines

- **Logistic Regression (LR):** Implemented using *scikit-learn* with $L_2$ regularization. A grid search was conducted for the regularization parameter $C \in \{0.001, 0.01, 0.1, 1, 10\}$. The optimal configuration used $C = 1.0$ with class weighting.
- **Support Vector Machine (Linear SVM):** Implemented with a linear kernel. $C$ was tuned over $\{0.1, 1, 10, 100\}$. The final model utilized $C = 1.0$ and calibrated probabilities via Platt scaling.
- **Simple MLP:** A feed-forward network consisting of two dense layers (128 and 64 units) with ReLU activation, followed by a Dropout layer ($p = 0.3$).

##### B. Temporal Baselines

- **Standard LSTM:** A vanilla LSTM architecture with a single LSTM layer (128 units), BatchNormalization, and a dense head.
- **Temporal Convolutional Network (TCN) [21]:** Modeled using dilated causal convolutions. Architecture: 3 residual blocks, dilation factors $d \in \{1, 2, 4\}$, kernel size $k = 3$, 64 filters.
- **Hybrid CNN-LSTM (phyLoSTM) [5]:** A domain-specific architecture featuring a 1D Convolutional layer (filters=64, kernel=3) followed by an LSTM layer (units=100).

TABLE S4
DOMINANCE ANALYSIS BY CLINICAL OBJECTIVE: IDENTIFYING THE SUPERIOR ARCHITECTURE FOR SPECIFIC METRICS.

| Target Metric | Comparison (Model A vs. Model B) | Mean Diff. | $\mathbf{P_{holm}}$ | Statistical Verdict |
|---|---|---|---|---|
| ***Discriminative Performance (ROC-AUC)*** | | | | |
| | **Stacking Ensemble** vs. Adapted TFT | +0.0109 | < 0.001 | **Ensemble Superior** |
| | **Stacking Ensemble** vs. Bi-LSTM | +0.0069 | < 0.001 | **Ensemble Superior** |
| | **Stacking Ensemble** vs. GRU + Attn | +0.0113 | < 0.001 | **Ensemble Superior** |
| ***Classification Reliability (MCC)*** | | | | |
| | **Adapted TFT** vs. Stacking Ensemble | +0.0200 | < 0.001 | **TFT Superior** |
| | **Adapted TFT** vs. Bi-LSTM | +0.0507 | < 0.001 | **TFT Superior** |
| | **Adapted TFT** vs. GRU + Attn | +0.0368 | < 0.001 | **TFT Superior** |
| ***Clinical Sensitivity (F1-Score Dysbiosis)*** | | | | |
| | **Bi-LSTM** vs. Stacking Ensemble | +0.0213 | < 0.001 | **Bi-LSTM Superior** |
| | **Bi-LSTM** vs. Adapted TFT | +0.0585 | < 0.001 | **Bi-LSTM Superior** |
| | **Bi-LSTM** vs. GRU + Attn | +0.0147 | < 0.001 | **Bi-LSTM Superior** |

*Note:* $P_{holm}$ denotes the p-value adjusted using the **HolmBonferroni method**. 'Mean Diff.' represents the average performance gap over 2,000 bootstrap iterations. The results confirm the **Adapted TFT's** dominance in reliability (MCC) versus the **Stacking Ensemble's** dominance in overall discrimination (ROC-AUC).

#### S7. FINAL HYPERPARAMETER CONFIGURATIONS

TABLE S3
HYPERPARAMETER SEARCH SPACE AND OPTIMAL VALUES

| Model | Hyperparameter | Search Space | Selected Value |
|---|---|---|---|
| **Bi-LSTM** | LSTM Units | $\{32, 64, 128, 256\}$ | 128 |
| | Dropout Rate | $[0.1, 0.5]$ | 0.3 |
| | Learning Rate | $\{1e^{-2}, 1e^{-3}, 1e^{-4}\}$ | $1e^{-3}$ |
| | Batch Size | $\{16, 32, 64\}$ | 32 |
| **GRU + Attention** | GRU Units | $\{32, 64, 128\}$ | 128 |
| | Attention Dim | Fixed (Sequence Length) | 14 |
| | Dense Units | $\{32, 64\}$ | 64 |
| | Dropout Rate | $[0.1, 0.5]$ | 0.25 |
| **Adapted TFT** | Hidden Dimension | $\{64, 128, 256\}$ | 128 |
| | Attention Heads | $\{2, 4, 8\}$ | 4 |
| | Dropout Rate | $[0.1, 0.4]$ | 0.2 |
| | Gradient Clipping | Fixed | 1.0 |

#### S8. COMPUTATIONAL ENVIRONMENT

- **Hardware:** Google Colab Pro+ (TPU v2-8 Runtime, 334 GB RAM).
- **Software:** Python 3.10.12, TensorFlow 2.15.0, Keras 2.15.0.
- **Libraries:** Pandas 2.0.3, NumPy 1.25.2, SHAP 0.44.1.

#### S9. MCC

#### S10. ERROR ANALYSIS

#### S11. ERROR ANALYSIS

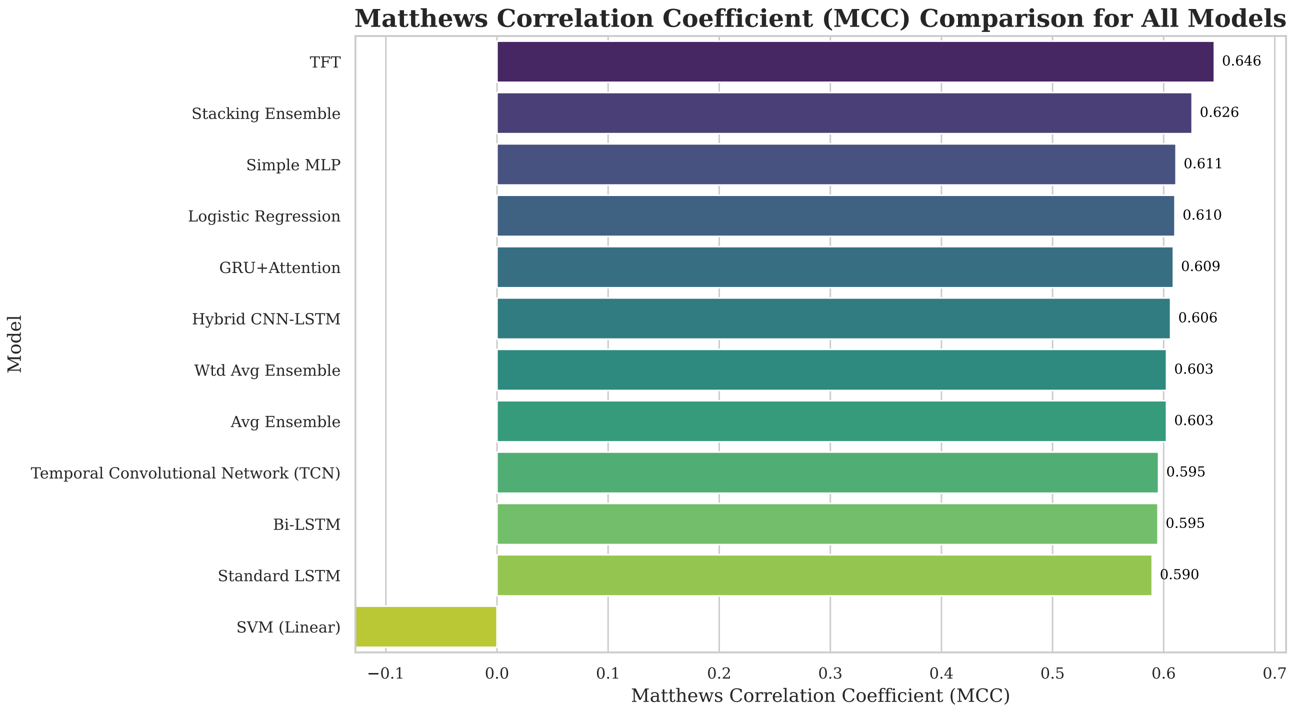


Fig. S3. Matthews Correlation Coefficient (MCC) comparison across all evaluated architectures, ranked by classification reliability. The Adapted TFT achieves the highest stability (MCC 0.646), validating the efficacy of Gated Residual Networks in filtering noise from high-dimensional microbiome data. While the Stacking Ensemble (MCC 0.626) and Simple MLP (MCC 0.611) perform competitively, the TFT is the only individual architecture to surpass the 0.64 threshold, acting as a robust sentinel against False Positives. In contrast, the Linear SVM fails to capture temporal dependencies, resulting in a negative MCC (-0.142) indicative of inverse prediction.

TABLE S5

**Supplementary Table S3:** Comparison of State-of-the-Art Deep Learning Models for Longitudinal Microbiome Analysis. The table highlights the pervasive reliance on legacy data structures (OTUs/Genera) and the lack of explicit architectural mechanisms for handling zero-inflation in sparse ASV data.

| Study | Model | Input Data | Interpretability | Critical Limitation / Gap |
|---|---|---|---|---|
| ***Group 1: Recurrent Neural Networks (RNNs)*** | | | | |
| Metwally et al. [?] | LSTM (Sparse Autoencoder) | Genus | Feature Selection (mRMR) | **Low Resolution:** Relies on genus-level aggregation, losing strain-level specificity. Latent representations failed to capture sparse signals. |
| Chen et al. [?] | GRU / Bi-GRU | OTU | None (Black-box) | **Legacy Constraints:** Relies on OTU clustering, ignoring the ASV standard. Validated only on binary classification. |
| Sharma & Xu [?] | CNN-LSTM (PhyloSTM) | OTU | None (Black-box) | **Artificial Structure:** Phylogenetic sorting imposes artificial spatial correlations on sparse data. Does not explicitly handle zero-inflation. |
| Karambelkar et al. [?] | Neural CDE | Genus | None (Black-box) | **Data Resolution:** Discards ASV-level variance via aggregation. Fails to address zero-inflation inherent in high-res data. |
| Sun & Zhou [?] | LSTM | Taxa (Relative Abundance) | Prediction Intervals | **Generalization:** Performance degrades significantly under environmental perturbations. Remains a black-box regarding biological drivers. |
| Dai et al. [?] | Ensemble (RNNs) | OTU | Permutation Importance | **Overfitting:** Advanced RNN variants overfitted on sparse data compared to simple NNs. Requires complete trajectories (imputation dependent). |
| ***Group 2: Generative & Imputation Models*** | | | | |
| Choi et al. [?] | GAN (DeepMicroGen) | OTU | None | **Regularity Assumption:** Struggles with irregular sampling intervals common in clinical data. Validated on OTUs, not ASVs. |
| Seki et al. [?] | Diffusion (CSDI) | OTU/Species | None | **Signal Attenuation:** Smoothing layers attenuate significant biological fluctuations (signal loss). High error rates for high-abundance taxa. |
| ***Group 3: Transformers & Proposed Framework*** | | | | |
| Myers et al. [?] | Transformer (TRPCA) | ASV | SHAP | **Modest Gains:** Statistical improvements over conventional ML are marginal, with overlapping uncertainty intervals. |
| **DynaBiomeX (Ours)** | **Ensemble + TFT** | **ASV (Sparse)** | **SHAP + Attention** | **Solution:** Explicitly models *structural* vs. *sampling* zeros using a dual-strategy (Screener-Sentinel). Achieved 100% Precision (0 False Positives). |

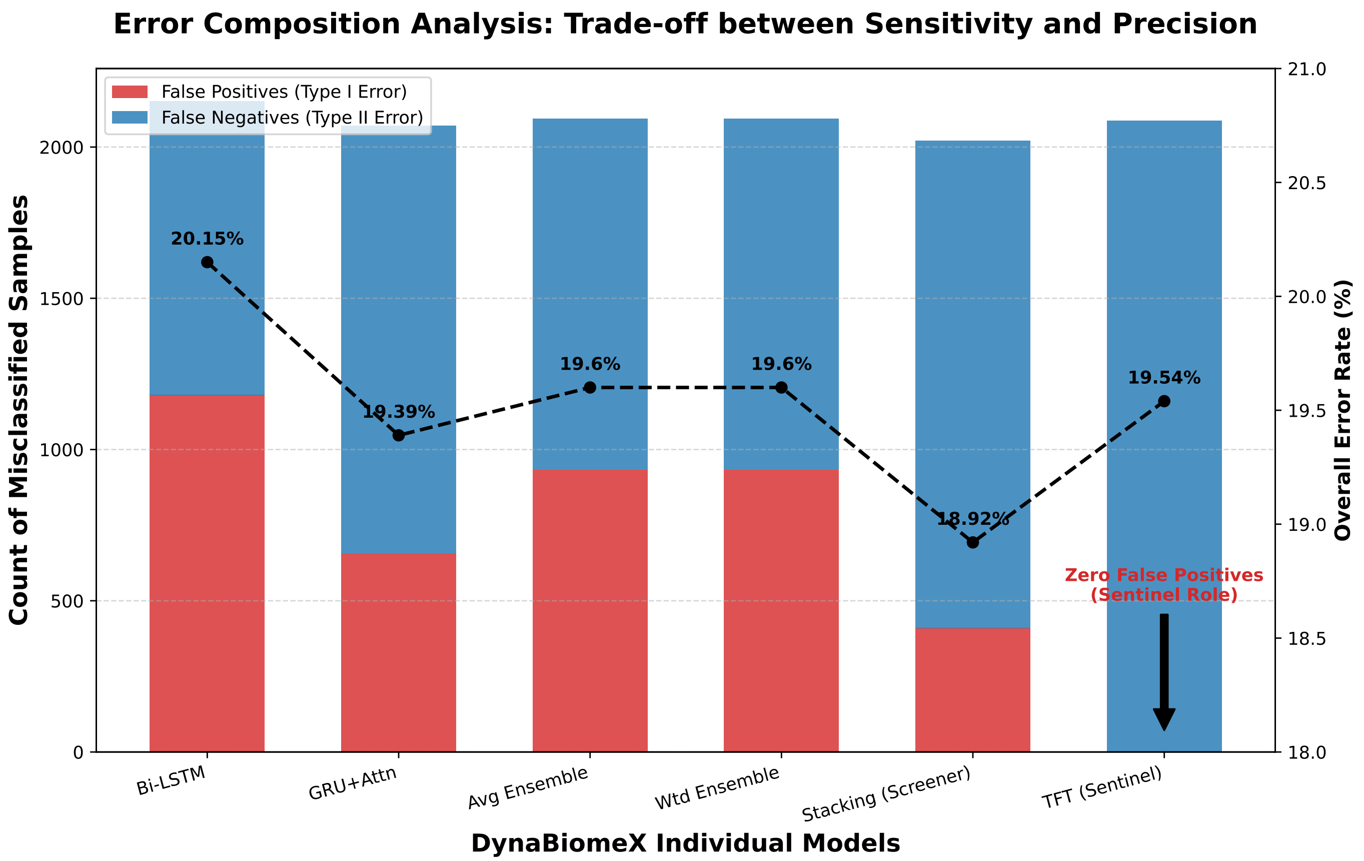


Fig. S4. Error Composition Analysis across Deep Learning Architectures. The stacked bars represent the total misclassified samples, decomposed into False Positives (Type I Error, Red) and False Negatives (Type II Error, Blue). The black dashed line indicates the overall Error Rate (%). This visualization highlights the architectural trade-off: the Stacking Ensemble minimizes total error (Screener), while the Adapted TFT eliminates False Positives entirely (Sentinel).
